# Loss of Carnitine Palmitoyltransferase 1a Reduces Docosahexaenoic Acid-Containing Phospholipids and Drives Sexually Dimorphic Liver Disease in Mice

**DOI:** 10.1101/2023.08.17.553705

**Authors:** Mikala M. Zelows, Corissa Cady, Nikitha Dharanipragada, Anna E. Mead, Zachary A. Kipp, Evelyn A. Bates, Venkateshwari Varadharajan, Rakhee Banerjee, Se-Hyung Park, Nathan R. Shelman, Harrison A. Clarke, Tara R. Hawkinson, Terrymar Medina, Ramon C. Sun, Todd A. Lydic, Terry D. Hinds, J. Mark Brown, Samir Softic, Gregory A. Graf, Robert N. Helsley

**Author notes:** <u>Corresponding author:</u> **Robert N. Helsley, PhD** Assistant Professor 741 S. Limestone St., BBSRB B347 Lexington, KY 40536.

## Abstract

**Background and Aims:** Genome and epigenome wide association studies identified variants in carnitine palmitoyltransferase 1a (CPT1a) that associate with lipid traits. The goal of this study was to determine the impact by which liver-specific CPT1a deletion impacts hepatic lipid metabolism.

**Approach and Results:** Six-to-eight-week old male and female liver-specific knockout (LKO) and littermate controls were placed on a low-fat or high-fat diet (HFD; 60% kcal fat) for 15 weeks. Mice were necropsied after a 16 hour fast, and tissues were collected for lipidomics, matrix-assisted laser desorption ionization mass spectrometry imaging (MALDI-MSI), kinome analysis, RNA-sequencing, and protein expression by immunoblotting. Female LKO mice had increased serum alanine aminotransferase (ALT) levels which were associated with greater deposition of hepatic lipids, while male mice were not affected by CPT1a deletion relative to male control mice. Mice with CPT1a deletion had reductions in DHA-containing phospholipids at the expense of monounsaturated fatty acids (MUFA)-containing phospholipids in both whole liver and at the level of the lipid droplet (LD). Male and female LKO mice increased RNA levels of genes involved in LD lipolysis (*Plin2*, *Cidec*, *G0S2*) and in polyunsaturated fatty acid (PUFA) metabolism (*Elovl5, Fads1, Elovl2*), while only female LKO mice increased genes involved in inflammation (*Ly6d, Mmp12, Cxcl2*). Kinase profiling showed decreased protein kinase A (PKA) activity, which coincided with increased PLIN2, PLIN5, and G0S2 protein levels and decreased triglyceride hydrolysis in LKO mice.

**Conclusions:** Liver-specific deletion of CPT1a promotes sexually dimorphic steatotic liver disease (SLD) in mice, and here we have identified new mechanisms by which females are protected from HFD-induced liver injury.

**Graphical Summary:** 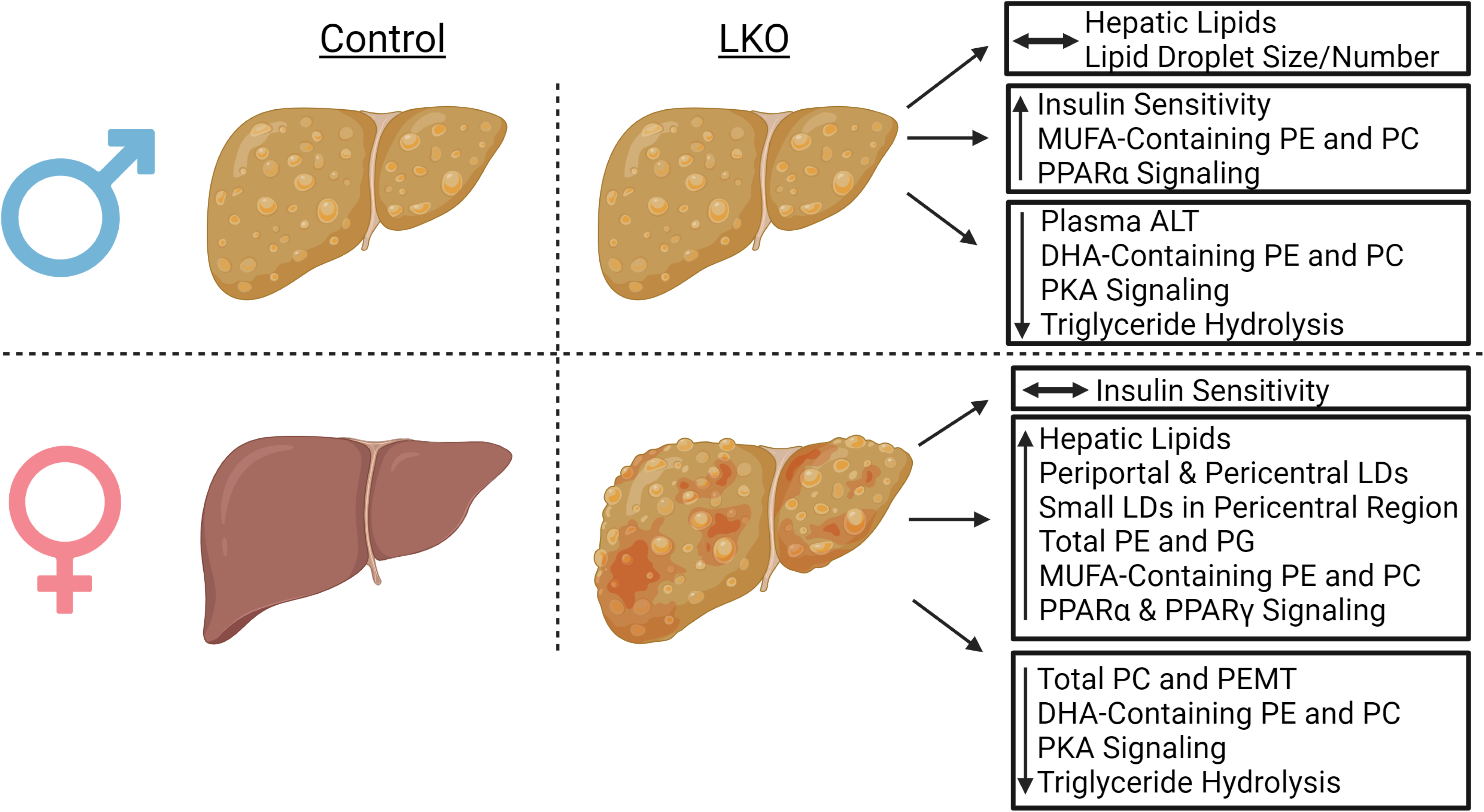

## INTRODUCTION

Steatotic liver disease (SLD) is a spectrum of disease etiologies ranging from simple steatosis to metabolic dysfunction-associated steatohepatitis (MASH) (1, 2). Over 64 million people in the United States are projected to have SLD, which is associated with a decrease in lifespan (3) and is expected to become the leading cause of liver-related morbidity and mortality over the next 15 years (4). Intriguingly, SLD is largely considered a sexually dimorphic disease with a greater prevalence in men (5), but women exhibit a greater risk of progressing to more severe SLD (6). Evidence from rodent and human studies has shown that pre-menopausal women are protected from SLD, in part, due to their ability to partition fatty acids towards oxidation and away from the secretory pathway (7, 8). While it is widely appreciated that estrogen elicits anti-inflammatory and pro-fatty acid oxidation (FAO) properties (9), less is known about downstream signaling pathways that elicit protection from SLD.

The carnitine palmitoyltransferase 1 (CPT1) gene family consists of three isoforms (1a, 1b, 1c) that differ in subcellular localization, tissue specificity, and function (10). The CPT1a and CPT1b isoforms are considered to be primarily expressed in the liver and muscle, respectively, where they reside on the outer mitochondrial membrane to convert long-chain acyl-CoAs to acylcarnitines for entry into the mitochondria and subsequent fatty acid oxidation (FAO) (10). Genome- and epigenome-wide associations studies have identified loss-of-function variants in CPT1a that associate with lower triglycerides (11-13), ω3-polyunsaturated fatty acids (PUFAs) including docosahexaenoic acid (DHA), and increased circulating monounsaturated fatty acids (MUFAs) (14). However, little is known about the mechanisms explaining these associations.

We characterized the contribution of hepatic CPT1a to hepatic lipid metabolic phenotypes in male and female mice. Using a CPT1a liver-specific deficient mouse model, we show female mice depend on CPT1a for protection from the high-fat diet (HFD)-induced liver dysfunction. Male mice are more susceptible to HFD-induced liver dysfunction than female control mice, a condition that is not worsened by the absence of CPT1a. Utilizing a combination of lipidomic approaches, we demonstrate that hepatic CPT1a-deficiency alters the total phospholipid pool and promotes the incorporation of monounsaturated (MUFA)-containing phospholipids at the expense of docosahexaenoic acid (DHA)-containing phospholipids in lipid droplets (LD). The shift in MUFA from DHA-containing phospholipids within LDs associates with increases in PUFA-biosynthetic and peroxisome proliferator activated receptor (PPAR)- target gene expression. Mechanistically, kinase profiling of liver samples shows reductions in protein kinase A (PKA) activity, which coincided with impaired triglyceride hydrolysis in CPT1a-deficient livers. These studies reveal new mechanisms by which impairments in long-chain fatty acid oxidation drive sexually dimorphic SLD and alter LD dynamics in mice.

## EXPERIMENTAL PROCEDURES

### Experimental Design and Study Subjects

Animal protocols were in accordance with NIH guidelines and approved by the Institutional Animal Care and Use Committee at the University of Kentucky. Mice were housed in individually ventilated cages at 20-22°C on a 14 h light/10 h dark cycle with ad libitum access to food and water. CPT1a-floxed mice were obtained from the laboratory of Dr. Peter Carmeliet (15), backcrossed >10 generations on the C57BL/6 background, and then were bred with the Albumin-Cre (Jackson Laboratory) transgenic mice to achieve liver-specific knockdown of CPT1a. Six to 8-week-old littermate controls (floxed & LKO) were fed either a semipurified low-fat diet (10% kcal from fat; Research Diets, D12450K) or a high-fat diet (60% kcal from fat; Research Diets, D12492) for 15-weeks prior to necropsy. Mice were necropsied after a 16-hour (4 PM – 8 AM) fast to induce adipose tissue lipolysis and concomitant hepatic FAO. Indirect calorimetry was completed on mice after 7 weeks of HFD feeding using the Promethion metabolic cages (Sable Systems International). Magnetic resonance relaxometry (Echo-MRI-100TM, Echo Medical System, Houston, TX) was used to assess total body fat, lean mass, and fat mass in mice at baseline, 7, and 15 weeks on the diet.

For intraperitoneal glucose tolerance tests, mice were fasted overnight and were injected into the peritoneal cavity with a 20% glucose solution at a volume of 10 µl per gram of body weight. For insulin tolerance tests, baseline glucose levels were measured in the fed state (at 8 AM), then mice were then injected into the peritoneal cavity with 1 U/kg of insulin. Serum glucose levels were measured using the INFINITY glucometer and glucose strips (ADW). Serum insulin (Crystal Chem) and β-hydroxybutyrate (Cayman Chem) levels were quantified using commercial assays according to manufacturers’ instructions.

### Quantification of Hepatic Lipids

Extraction of liver lipids and subsequent quantification of hepatic triglycerides, total cholesterol, and phosphatidylcholine were conducted using enzymatic assays as previously described (16, 17). Liver samples were delipidated in 4 mL of 2:1 chloroform to methanol (v/v) overnight. The organic solvent was dried under a constant stream of N_2_ prior to adding 6 mL of 2:1 chloroform to methanol. To separate the phases, 1.2 mL of 0.05% H_2_SO_4_ was added, samples were vortexed then centrifuged at 2000 rpm for 15 minutes. The bottom (organic) phase was recorded, and 0.5 mL of the organic phase was added to 1 mL of 1% TritonX-100 in chloroform. The samples were dried under N_2_ and 0.5 mL H_2_O was added prior to quantification of triglycerides (Wako), total cholesterol (Pointe Scientific), and choline-containing phospholipids (Sigma), per manufacturer’s instructions. All standards and blanks were prepared in a similar fashion, and data were presented normalized to initial liver weights.

### Histological Analysis

Hematoxylin and eosin (H&E) staining of formalin-fixed paraffin-embedded liver sections was performed as previously described (16, 17). LD number and size were calculated by blinded members of the laboratory. In short, 2 individuals (independent of one another) captured 3 fields surrounding the portal vein and 3 fields surrounding the central vein at 100x magnification on a Nikon AR15. A total of 6 fields were captured per section, per mouse, for a total of 240 fields (e.g., 2 sections/per mouse for 20 total mice). The diameter was calculated for each individual droplet using Nikon Elements. The total number of LDs was normalized to the total nuclei in a given field.

### Serum ALT Levels

To determine the degree of liver injury in control and LKO mice, alanine aminotransferase (ALT) levels were measured in serum using enzymatic assays (Sekisui Diagnostics, Lexington, MA, USA), as previously described (16, 17).

### Shotgun Lipidomics

#### Lipid extraction

Liver samples on dry ice were spiked with 10 microliters of an internal standard and calibration mixture consisting of 500 μM each of di-myristoyl phospholipids (PG, PE, PS, PA, PC), 500 μM SM (30:1) and 50 μM TG (14:1/14:1/14:1). To each sample, 300 microliters of -20°C chilled 75% methanol containing 0.01% BHT (butylated hydroxytoluene) were added along with 0.5 mm zirconium oxide beads. Samples were homogenized briefly in a Bullet Blender tissue homogenizer and placed on ice. Sixty microliters of methanol and 1 mL of MTBE were added to each sample, and samples were then vortexed for 60 minutes at room temperature. Water (170 μL) was added, and the samples were vortexed for an additional 15 minutes and then centrifuged for 15 minutes. The supernatants were collected to new test tubes and precipitated proteins were re-extracted as above. Pooled extracts were dried overnight in a speedvac, and resuspended in 400 μL of isopropanol containing 0.01% BHT.

#### Lipidomics analysis by Orbitrap high resolution/accurate mass spectrometry

Immediately prior to analysis, aliquots of each lipid extract were diluted 50-fold in isopropanol: methanol (2:1, v:v) containing 20 mM ammonium formate. Full scan MS spectra at 100,000 resolution (defined at m/z 400) were collected on a Thermo Scientific LTQ-Orbitrap Velos mass spectrometer in both positive and negative ionization modes. Scans were collected from m/z 200 to m/z 1200. For each analysis, 10 µL of sample was directly introduced by flow injection (18, 19) at 10 µL/min using an electrospray ionization source equipped with a fused silica ESI needle to minimize intrasource accumulation of triglycerides. A Shimadzu Prominence HPLC with thermostatted autosampler served as the sample delivery unit. The sample and injection solvent were 2:1 (v: v) isopropanol: methanol containing 20 mM ammonium formate. The spray voltage was 4.5 kV, ion transfer tube temperature was 275 °C, the S-lens value was 50 percent, and the Orbitrap fill time was 100 ms. The autosampler was set to 4 °C. After two minutes of MS signal averaging, the LC tubing, autosampler, and ESI source were flushed with 1 mL of isopropanol, prior to injection of the next sample. Samples were analyzed in random order, interspersed by solvent blank injections, extraction blank injections, and pooled QC samples derived from all study samples. Following MS data acquisition, offline mass recalibration was performed with the "Recalibrate Offline" tool in Thermo Xcalibur software according to the vendor’s instructions, using the theoretical computed masses for the internal calibration standards and several common endogenous mammalian lipid species. MS/MS confirmation and structural analysis of lipid species identified by database searching were performed using higher-energy collisional dissociation (HCD) MS/MS at 60,000 resolution and a normalized collision energy of 25 for positive ion mode, and 60 for negative ion mode. MS/MS scans were triggered by inclusion lists generated separately for positive and negative ionization modes.

#### Lipid Peak Finding, Identification, and Quantitation

Lipids were identified using the Lipid Mass Spectrum Analysis (LIMSA) v.1.0 software linear fit algorithm, in conjunction with an in-house database of hypothetical lipid compounds, for automated peak finding and correction of ^13^C isotope effects as previously described (20). Peak areas of found peaks were quantified by normalization against an internal standard of a similar lipid class. The top ∼300 most abundant peaks in both positive and negative ionization mode were then selected for MS/MS inclusion lists and imported into Xcalibur software for structural analysis on the pooled QC sample as described above. For this untargeted analysis, no attempt was made to correct for differences in lipid species ionization due to the length or degree of unsaturation of the esterified fatty acids. Therefore, lipid abundance values are inherently estimates rather than true ‘absolute’ values, and so all data are presented as % of the total pool of lipid.

### Spatial Lipidomics

High-performance liquid chromatography (HPLC)-methanol, HPLC-grade water and N-(1-Naphthyl)ethylenediamine dihydrochloride (NEDC) matrix were obtained from Sigma-Aldrich. Slides were prepared similar as previously described (21). After desiccation for one hour, slides were sprayed with 14 passes of 7 mg/mL NEDC matrix in 70% methanol, applied at 0.06 mL/min with a 3mm offset and a velocity of 1200mm/min at 30°C and 10psi using the M5 Sprayer with a heated tray of 50°C. Slides were used immediately or stored in a desiccator until use. For the detection of lipids, a Bruker timsTOF QTOF high-definition mass spectrometer was used. The laser was operating at 10000 Hz with 60% laser energy, with 300 laser shots per pixel and spot size of 50µm at X and Y coordinates of 50µm with mass range set at 50 – 2000m/z in negative mode. Data acquisition spectrums were uploaded to Scils Software (Bruker Corporation) for the generation of small molecule and lipid images. Regions of interest (ROIs) were drawn around the whole tissue. For all pixels defined within a ROI, peak intensities were averaged and normalized by total ion current.

### LD Isolation from Mouse Liver

Male and female control and LKO mice were fed a HFD for 15-weeks. After 15-weeks of feeding, mice were necropsied in the fasted state and LDs were isolated by sucrose gradient centrifugation as we have previously described (17). LD PC and PE lipids were extracted and quantified using the targeted LC-MS/MS method below (17).

### Relative Quantitation of Phosphatidylcholine (PC) and Phosphatidylethanolamine (PE) Lipid Species in LD Fractions

A targeted lipidomic assay for PC and PE lipids was developed using HPLC on-line electrospray ionization tandem mass spectrometry (LC/ESI/MS/MS), as previously described (17).

#### Standard Solutions and Lipid Extraction

The standards (PC_36:4 and PE_34:2) and the isotope labeled internal standards (PC_33:1-d7 and PE_31:1-d7) were purchased from Avanti Polar Lipids (Alabaster, Alabama, USA). The Standard solution at concentrations of 0, 10, 50, 200, 1000, 5000 and 20000 ng/ml were prepared in 80% methanol containing the internal standard at the concentration of 500 ng/ml. Tissue homogenate with a volume of 20 μl was mixed with 80 μl methanol containing 2 internal standards at the concentration of 625 ng/ml, vortexed for 30 sec and then centrifuged at 18000 rcf, 4 C for 12 min. After centrifugation, 50 ul supernatant was transferred into a HPLC vial for injection. The volume of 5 μl standard solution and extracted sample was injected into the Vanquish UHPLC system (Thermos Fisher Scientific, Waltham, MA, USA) for lipid separation.

#### LC/MS/MS Parameters

A C18 column (2.1 x 150mm, Gemini, 3 μm, Phenomenex,) was used for the separation of the lipid species. Mobile phases were A (water containing 0.1% acetic acid and 0.3% ammonium hydroxide) and B (methanol/acetonitrile, 1/1 (v/v)) containing 0.1% acetic acid and 0.3% ammonium hydroxide). Mobile phase B at 80 % was used from 0 to 2 min at the flow rate of 0.3 ml/min and then a linear gradient from 80 % B to 100 % B from 2 to 8 min, kept at 100 % B from 8 to 20 min, 100 % B to 80 % B from 20 to 20.1 min, kept 80 % B from 20.1 to 29 min. The HPLC eluent was directly injected into a triple quadrupole mass spectrometer (TSQ Quantiva, Thermos Fisher Scientific, Waltham, MA, USA) and both the PC and PE lipids were ionized at the positive mode. The PC species were monitored using Selected Reaction Monitoring (SRM) and the SRM transitions were the mass to charge ratio (m/z) of molecular cation to the daughter ion m/z 184, the specific phosphocholine group. The PE species were monitored using Selected Reaction Monitoring (SRM) and the SRM transitions were the mass to charge ratio (m/z) of molecular cation to the daughter ion at m/z minus 141, the specific ethanolamine group.

#### Data Analysis

Software Xcalibur was used to get the peak area for all the lipid species. The internal standard calibration curves were used to calculate the relative concentration of all the PC and PE species in the samples.

### PamGene PamStation Sample Preparation

Kinase activity was measured using serine-threonine kinases (STK) PamChip4 porous 3D microarrays and measured using the PamStation12 (PamGene International, ’s-Hertogenbosch, The Netherlands). Substrates contained in each array are listed in **Supplemental Table 1**. Mouse livers were pooled and measured in triplicate across three chips simultaneously for STK, as previously described in (22-24). This approach effectively deals with large batch effects across samples. It allows for the characterization of kinase activity only in the context of analytical variance. The pooled samples were lysed using M-PER Mammalian Extraction Buffer (Thermo Fischer Scientific, CAT#78503), Halt Phosphatase Inhibitor (Thermo Fischer Scientific, CAT#78428), and Protease Inhibitor Cocktail (Sigma, CAT#P2714). The samples were homogenized using TissueLyser LT (Qiagen). The protein concentration was measured in triplicate using Pierce BCA Protein assay (Thermo Fischer Scientific, CAT#23225). Samples were diluted to a final protein concentration of 2.5 μg/μl. Each array contained 1 μg of protein per sample for the STK chips. In the presence of adenosine triphosphate (ATP), kinase phosphorylation activity is quantified using fluorescently labeled antibodies to detect differential phosphorylation of 144 (STK) reporter peptides between experimental and control conditions, as previously described (25). Evolve (PamGene) software uses a charge-coupled device (CCD) camera and light-emitting diode (LED) imaging system to record relative phosphorylation levels of each unique consensus phosphopeptide sequence every 5 minutes for 60 minutes as measured by peptide signal intensities recorded across 10, 20, 50, and 100 millisecond exposure times. Raw imaging data were exported for further data analysis and kinase mapping.

### PamGene PamStation Kinase Data Analysis

The images taken during the run were analyzed using BioNavigator (PamGene). Signal ratios are used to calculate fold change (FC) for each phosphopeptide sequence averaged across three replicates. Minimum threshold values were selected using cutoffs cited in previous literature (25-28). These thresholds require differential phosphopeptide signals greater than or equal to 30% (FC ≥ 1.30 or FC ≤ 0.70) for differential phosphorylation to be considered. Linear regression slopes provide phosphorylation intensity signals used in differential analyses (e.g., experimental vs. control). Undetectable and/or nonlinear (R2 < 0.80) phosphopeptides are excluded from subsequent analyses. We performed upstream Kinase identification using Kinome Random Sampling Analyzer (K.R.S.A.) (29) and Upstream Kinase Analysis (U.K.A.) (30), as previously described in (25). The kinase scores from the K.R.S.A. and U.K.A. are included in **Supplemental Tables 2 (males) and 3 (females)**. MEOW (measurements extensively of winners) plots, as described in (31), were used to interrogate individual kinase activities on substrates considering the confidence of the experimental versus the control groups using the equation: [Log2 Fold Change (FC) of kinase substrates * Δconfidence (experimental hits/mean hits of 2000 random sampling iterations)]. Data input files for STK runs available on the GitHub repository at the following (https://github.com/The-Hinds-Lab/Liver-Specific-Cpt1a-KO-Kinome-Analysis).

### Real-Time PCR Analysis of Gene Expression

RNA extraction, cDNA synthesis, and quantitative real-time PCR was performed as previously described (16, 17). A small portion of frozen livers (∼20 mg) were homogenized in 1 mL of QIAzol (Qiagen 79306). After the addition of 200 µL of chloroform, phase separation occurred by centrifugation and the top layer was mixed with 400 µL of 75% ethanol prior to running the sample over a RNeasy spin column (Qiagen, RNeasy Mini Kit). 500 ng of RNA was used as a template to synthesize cDNA (High Capacity cDNA Reverse Transcription Kit, Applied Biosystems). QPCR reactions were carried out using 2X SYBR green (Cowin Biosciences) and mRNA expression levels were calculated using the ΔΔ- Ct method on an Applied Biosystems (ABI) QuantStudio 7 Flex Real-Time PCR System. Primers used for qPCR are listed in **Supplemental Table 4**.

### Bulk RNA-Sequencing

RNA was isolated using the RNeasy Mini kit (Qiagen) from livers of male and female, WT and LKO mice fed HFD for 15-weeks. The quantity and quality of the samples were determined using the Cytation 5 (BioTek) plate reader and Agilent 4150 Tape Station System, respectively, prior to submission to Novogene. The mRNA-seq libraries were prepared and sequenced on an Illumina HiSeq2500 platform, at a depth of 20 M read pairs (or 10M/sample) by Novogene. After sequencing, the paired-end clean reads were aligned to the reference genome using Hisat2 v2.0.5. FeatureCounts v1.5.0-p3 was used to count the read numbers mapped to each gene, and the fragments per kilobase of transcript (FPKM) of each gene was calculated based on the length of the gene and read counts mapped to the gene.

Differential expression analysis was performed using the DESeq2 R package (1.20.0). The resulting p-values were adjusted using the Benjamini and Hochberg’s approach for controlling the false discovery rate. Genes with an adjusted p-value ≤0.05 found by DESeq2 were assigned as differentially expressed. Gene ontology (GO) enrichment analysis of differentially expressed genes was implemented by the clusterProfiler R package. GO terms with corrected p-values <0.05 were considered significantly enriched by differential expressed genes. The clusterProfiler R package was also used to test the statistical enrichment of differentially expressed genes in KEGG pathways (http://www.genome.jp/keg/).

### Immunoblotting

Whole liver (∼20 mg) homogenates were solubilized in 1 mL of 1X ice-cold RIPA buffer (Cell Signaling) supplemented with 1X protease and phosphatase inhibitor cocktail (bimake.com). After two sequential centrifugation steps (13,000 rpm for 10 min), the supernatant was collected and protein was quantified using the Pierce^TM^ Bicinchoninic Acid Protein (BCA) assay (ThermoFisher). For SDS-PAGE, 10 µg of protein was loaded and separated on 4-15% criterion TGX gels. Proteins were blocked with 5% milk and probed with antibodies listed in **Supplemental Table 5**. Images were taken on a ChemiDoc MP Imaging System (Bio-Rad) and quantification of blots were performed using ImageJ software (NIH).

### TAG Hydrolysis Measurements

Triacylglycerol hydrolysis experiments were conducted as previously described (32). A small piece of liver tissue was first homogenized in assay buffer containing 20 mM Tris-HCl, 150 mM NaCl, and 0.05% Triton X-100 (pH=8.0). Lysates were spun at 15,000 x *g* for 15 minutes and the infranatant was collected for a Pierce BCA protein assay. 50 μg of protein lysate was then brought up to 180 uL with assay buffer and added to a black, 96 well microtiter place on ice. The substrate resorufin ester (Sigma D7816) was prepared in 0.3 mg/mL of assay buffer, and 20 uL was then added to all samples. Kinetic readings were taken every 2 minutes at 530 nm excitation and 590 emission. Moles of resorufin ester hydrolyzed are calculated from a standard curve generated using free resorufin.

### Transmission Electron Microscopy

A 1 mm x 3 mm piece of liver was fixed with 2.5% glutaraldehyde/4% paraformaldehyde in 0.2 M cacodylate buffer overnight at 4°C. The sample was then washed 3 times with sodium cacodylate buffer (0.2M, pH 7.3) at 5 min/wash. The buffer was removed, the samples were post-fixed with 1% Osmium Tetroxide (in H2O) for 60 min at 4°C. The samples were washed 2 times with sodium cacodylate buffer for 5 min/wash, then rinsed with 2% Maleate buffer (pH 5.1) for 5 min. The buffer was changed to 1% uranyl acetate in Maleate buffer and stained for 60 min at room temperature (RT). After staining, the uranyl acetate was removed and washed with maleate buffer 3 times at 5 min each. Dehydration is followed by different concentrations of cold ethanol 30%, 50%, 75%, 95% 1X, 5 min/wash followed by 100% ethanol at RT 3 times for 10 min/wash. For the infiltration step, the 100% ethanol was removed, and 1:1 ethanol/eponate12 medium was added at RT overnight. The media was then removed and changed to pure eponate 12 medium for 4-6 hrs at RT. For embedding, the sample was placed into pure eponate 12 in a rubber mold and allowed to polymerize for 24 h in an oven at 62°C. Ultra-thin sections (85nm) were cut with diamond knife, stained with 10% uranyl acetate and lead citrate, and then observed with a Tecnai G2 SpiritBT electron microscope operated at 80 kV.

### Statistical Analysis

All graphs and statistical analyses were completed using GraphPad Prism 9.3.0. Data are expressed as ±SEM, unless otherwise noted in the figure legends. Differences were computed using two-way ANOVAs followed by a Tukey’s multiple comparison *post hoc* analysis. When comparing two groups, a Student’s *t*-test was utilized. P-values <0.05 were considered statistically significant. All figure legends contain the statistical analysis used for each panel of data.

## RESULTS

### Liver-Specific Deletion of CPT1a Does Not Influence Body Weight or Adiposity

To examine the mechanisms by which excess dietary fatty acids alter hepatic lipid metabolism in a sex- and CPT1a-dependent manner, we fed 6–8 week old male and female, *Cpt1a*-liver specific knockout (LKO) and littermate *Cpt1a*^F/F^ (Control) mice a semipurified low-fat (LFD; 10% kcal fat) or high-fat diet (HFD; 60% kcal fat) for 15 weeks. An experimental scheme has been provided in **Fig. S1A**. Mice were necropsied after a 16-hour fast to induce adipose tissue lipolysis and concomitant hepatic FAO (**Fig. S1A**) (33). To confirm successful deletion of *Cpt1a* in the liver, we measured *Cpt1a* RNA and protein levels in livers collected from LFD- and HFD-fed mice. Real-Time PCR (qPCR) analysis revealed a 70-80% reduction in *Cpt1a* RNA in male and female LKO mice fed LFD (**Fig. S1B**) and HFD (**Fig. 1A**). Immunoblotting followed by densitometry confirmed CPT1a protein levels were significantly reduced in LKO mice (**Fig. 1B-C, Fig. S1C-D**). Previous reports have shown that another Cpt1 isoform, *Cpt1b*, may compensate for the loss of *Cpt1a* (34). We then measured the relative expression of *Cpt1b* by qPCR and found it to be elevated with the loss of *Cpt1a* in male mice fed either a LFD (**Fig. S1E**) or a HFD (**Fig. 1D**). Female LKO mice exhibited increased *Cpt1b* gene expression with LFD feeding (**Fig. S1E**) but was reduced in LKO mice fed a HFD, relative to control mice (**Fig. 1D**). Despite relative compensatory changes in *Cpt1b* measured by qPCR, total read counts of the *Cpt1b* transcript by RNA-sequencing showed it is ∼10,000x lower than *Cpt1a* (**Fig. S2A, B**) and is undetectable by western blot (**Fig. S2C**). Consistent with loss of the predominant Cpt1 isoform in the liver, male and female LKO mice displayed a 77% and 62% reduction in fasting serum β-hydroxybutyrate levels, a surrogate for hepatic FAO (35), upon LFD and HFD feeding, respectively (**Fig. 1E, Fig. S1F**).

**Figure 1.**
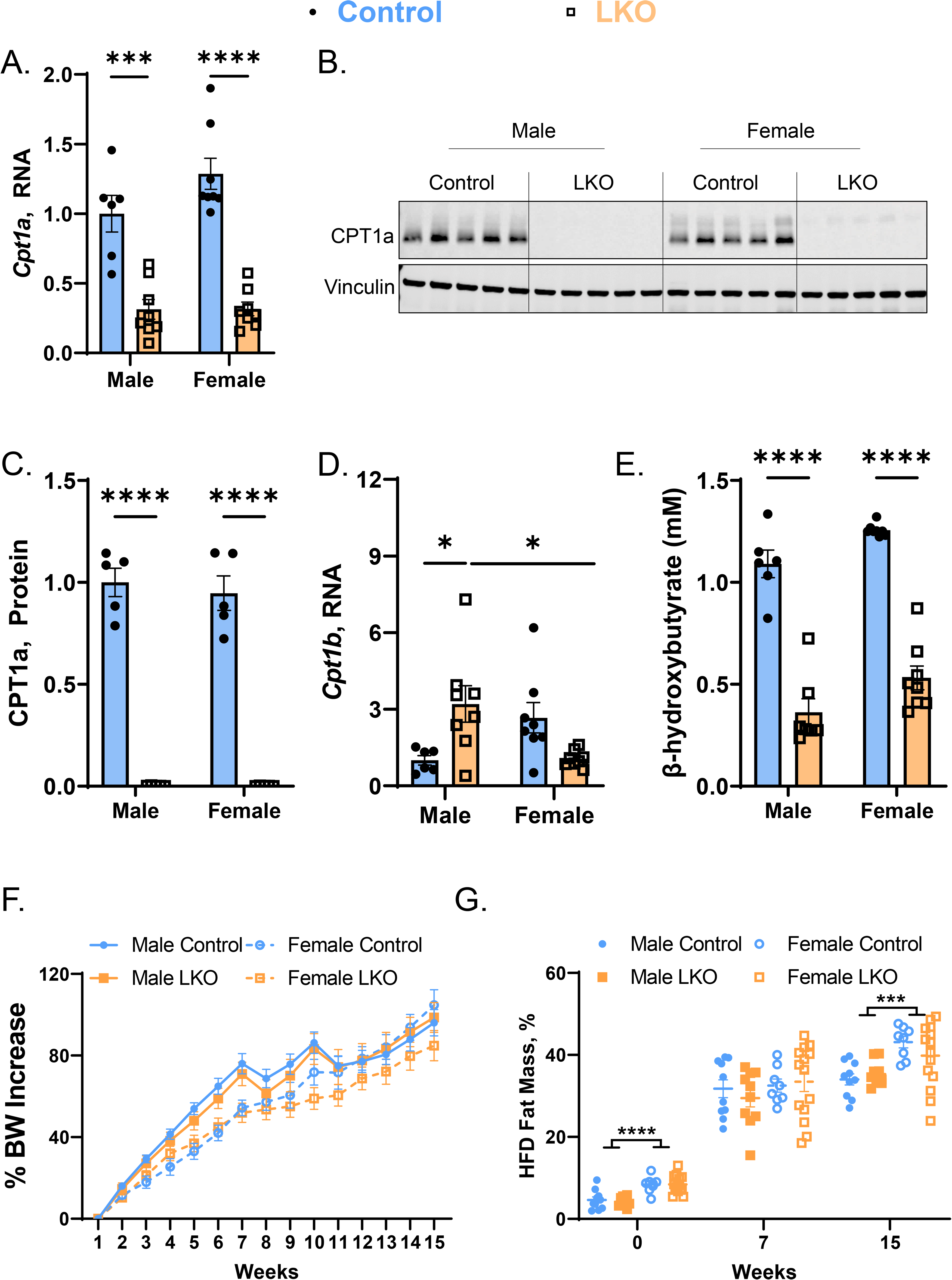
Liver-specific CPT1a Deletion Has No Effect on Body Weight in Response to HFD-Feeding. Male and female control and LKO mice were fed a HFD for 15-weeks. (**A**-**C**) Liver CPT1a RNA and protein levels were measured by qPCR (**A**; n=6-8) and western blot (**B**; n=5) followed by densitometry (**C**; n=5), respectively. Vinculin is used as a loading control. (**D**) *Cpt1b* RNA levels were measured by qPCR from livers of HFD-fed male and female control and LKO mice (n=6-8). For all qPCR analyses, housekeeping genes *Tbp* and *Hprt* were averaged and used for normalization. (**E**) β-hydroxybutyrate levels were measured from the plasma of fasted mice (n=6-8). (**F**, **G**) Percent body weight (**F**) and fat (**G**) mass were recorded throughout the duration of the study (n=8-14). Significance was determined by two-way ANOVA with Tukey’s multiple comparison *post hoc* analysis. *P<0.05; ***P<0.001; ****P<0.0001.

To determine if LKO mice exhibited changes in overall adiposity prior to dietary manipulation, we measured baseline adiposity in 6-8 week old control and LKO mice by Echo MRI. Baseline body weights were elevated in males compared to females, however loss of CPT1a in the liver had no impact on overall body weight (**Fig. S3A**), fat mass (**Fig. S3B**), or lean mass (**Fig. S3C**) compared to their respective controls. Next, we measured body weights and adiposity in response to LFD- and HFD-feeding for 15-weeks. Control and LKO mice fed LFD had comparable increases in body weight (∼23-35% increase from baseline) and percent fat mass throughout the 15-week study, regardless of sex or genotype (**Fig. S3D, E**). When challenged with HFD, male and female mice increased their body weight by ∼100% but no significant differences were observed across genotypes (**Fig. 1F**). Consistent with baseline (**Fig. S3A-C**), female mice had greater fat mass (∼41% fat mass) than male mice (∼34% fat mass) at the end of 15-weeks of HFD-feeding, independent of hepatic CPT1a (**Fig. 1G**). Taken together, LKO mice had reduced hepatic CPT1a protein and ketone levels with fasting; however, there were no observable differences in adiposity or body weight gain in response to LFD- or HFD-feeding.

### Liver-Specific CPT1a Deficiency Alters Food Intake, Total Activity, and Energy Expenditure in Female Mice

Alterations in both substrates (free fatty acids; FFAs) and products (long chain acylcarnitines) of CPT1a have been associated with insulin resistance (36, 37). This prompted us to investigate whether liver-specific deletion of *Cpt1a* would affect systemic glucose tolerance and insulin sensitivity in mice. Following 10 weeks on diet, mice were fasted overnight and underwent a glucose tolerance test (GTT). In response to LFD, male and female LKO mice exhibited similar fasting glucose levels as control mice (**Fig. S4A**) and no differences were observed in glucose tolerance or insulin sensitivity in male and female LKO mice (**Fig. S4A-D**). Similar to LFD-feeding, HFD-fed LKO mice showed similar fasting glucose levels as control mice (**Fig. S5A**); however, male LKO mice had significant lower fasting insulin levels (0.83±0.39 ng/mL) as compared to male control mice (2.16±1.0 ng/mL; **Fig. S5B**). These reductions in fasting insulin levels drove improvements in the homeostatic model assessment for insulin resistance (HOMA-IR; **Fig. S5C**), a surrogate marker for insulin resistance (38).

We then utilized indirect calorimetry to examine the effects of liver-specific deletion of *Cpt1a* on metabolic parameters in male and female mice in response to HFD-feeding. Notably, control and LKO mice were challenged with an overnight fast during night 4 of the five-day experiment. Over the course of the 5-day experiment, male control and LKO mice exhibited no differences in cumulative food intake during the light and dark cycles (**Fig. S6A-C**). Female LKO mice, however, consumed ∼10% more calories than female control mice, which was observed across the light and dark cycles (significance by genotype = 0.0018; **Fig. S6D-F**). Consistent with changes in food intake, female LKO mice were ∼20% more active and expended more energy (significance by genotype=0.0122) as compared to female control mice (**Fig. S7D-F**; **S8D**). Taken together, female LKO had enhanced caloric intake which associated with increased total activity and energy expenditure, leaving the mice isocaloric compared to their littermate controls throughout the duration of the study (**Fig. 1F, G**).

### Female CPT1a LKO Mice Exhibit Panlobular Microvesicular Steatosis and Exacerbation of Liver Injury in Response to HFD-Feeding

Hepatic steatosis, a hallmark feature of SLD, is defined by the accumulation of triacylglycerol and cholesterol esters in the core of cytosolic lipid droplets (39, 40). Hepatic lipid droplet accumulation can be driven by prolonged fasting (e.g. fasting-induced steatosis) and by overnutrition (e.g. HFD-feeding). We first assessed the impact of liver-specific *Cpt1a*-deficiency on fasting-induced steatosis with LFD-feeding and found that male and female LKO mice had increased liver weights and hepatic triglycerides, as compared to their respective controls (**Fig. S9A, B**) (41). Meanwhile, hepatic cholesterol levels were significantly increased only in female LKO mice (**Fig. S9C**), while no differences were observed in serum alanine aminotransferase (ALT) levels, a marker of liver injury (**Fig. S9D**). In response to HFD, female control mice were protected against diet-induced liver injury and had reduced liver weights (34.5%), hepatic triglycerides (38.9%), and serum ALT levels (67.9%) as compared to male control mice (**Fig. 2A-D**). Female LKO mice, however, completely lost this protection characterized by increased liver weights (53.9%), hepatic triglycerides (141.3%) and cholesterol (129.1%), and serum ALT levels (67.9%), as compared to female control mice (**Fig. 2A-D**). Notably, male LKO mice had comparable liver weights and hepatic lipids (triglycerides, cholesterol), as well as reduced ALT levels (-42.6%) as compared to male control mice (**Fig. 2A-D**).

**Figure 2.**
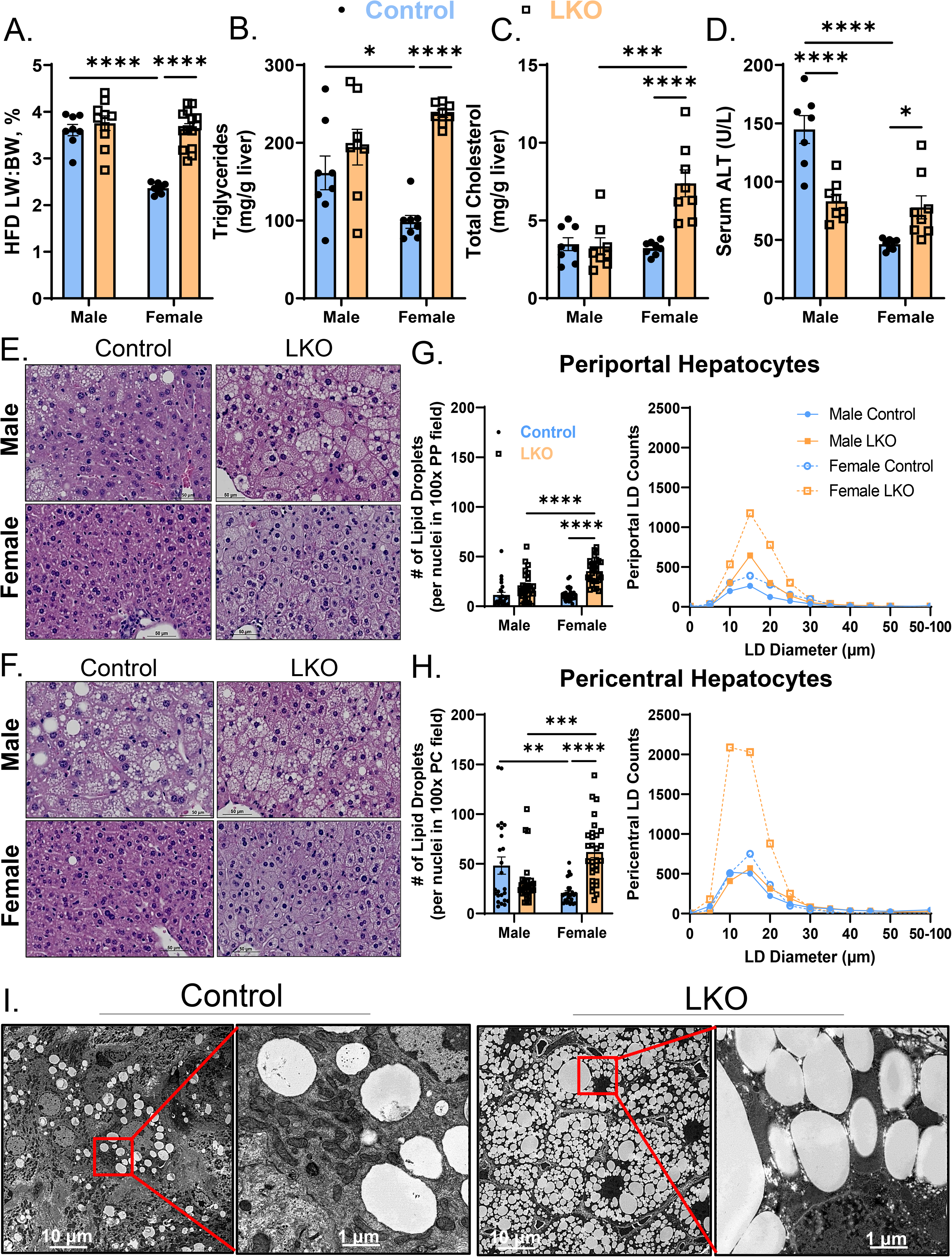
Female LKO Mice Develop Exacerbated Steatosis and Liver Dysfunction in Response to HFD-Feeding. Male and female control and LKO mice were fed a HFD for 15-weeks. (**A**-**C**) Liver weights normalized to body weight (**A**; LW:BW, %; n=8-14), and hepatic triglycerides (**B**; n=8) and total cholesterol (**C**; n=8) levels were quantified enzymatically. (**D**) Serum ALT levels were quantified (n=7-8). (**E, F**) Representative Oil Red O staining to assess neutral lipid accumulation across groups. The scale bar (50 µm) is embedded within the lumen of the portal (**E**) or central (**F**) veins in the corner of each image. Total number of lipid droplets (# lipid droplets/nuclei in 100X field) and their respective diameters were quantified in periportal (**G**) and pericentral hepatocytes (**H**). A total of 17,473 lipid droplets were quantified across 19 mice. (**I**) Transmission electron microscopy was completed on livers collected from HFD-fed female control and LKO mice (scale bars = 10 µm on left image and 1 µm on right image). Significance was determined by two-way ANOVA with Tukey’s multiple comparison *post hoc* analysis. *P<0.05; **P<0.01; ***P<0.001; ****P<0.0001.

The liver lobule can be broken down into anatomical zones based off their metabolic function. Hepatocytes occupying the periportal region (zone 1) are exposed to oxygen-rich blood flow and undergo greater rates of FAO (42, 43). Hepatocytes occupying the pericentral region (zone 3) are exposed to more hypoxic-like conditions and undergo more triglyceride synthesis (42, 43). The accumulation of LDs observed in SLD typically affects pericentral hepatocytes (zone 3), which are distinct from portal-based diseases such as viral hepatitis (44). Given FAO is highly active in periportal hepatocytes, we asked whether we could drive LD accumulation in periportal hepatocytes. To assess this, we stained liver sections with hematoxylin and eosin (**Fig. 2E, F**) and blindly quantified the size and total number (per nuclei) of 17,473 LDs across the periportal-pericentral axis. Male control mice displayed significant pericentral steatosis (**Fig. 2H**) but were largely free from lipid droplet accumulation in periportal hepatocytes (**Fig. 2G**). Male LKO mice displayed slight (albeit not significant) periportal steatosis, while pericentral steatosis was unaffected relative to male control mice (**Fig. 2E-H**). Female control mice were protected from diet-induced LD accumulation as compared to male control mice; however, LKO mice displayed diffuse, panlobular microvesicular steatosis across the entire periportal-pericentral axis (**Fig. 2E-H**). Consistently, female LKO mice had increased total LD number (**Fig. 2G, H**) and a greater propensity to accumulate smaller LDs (37% of LDs within 5-10 μm) in the pericentral zone, as compared to female control mice (27.5% of LDs within 5-10 μm; **Fig. S10A, B**). Further, transmission electron microscopy revealed significant LD accumulation in female LKO mice as compared to female control mice (**Fig. 2I**). Taken together, female mice are protected from diet-induced SLD in a *Cpt1a*-dependent manner, while male mice are largely unaffected by *Cpt1a*-deletion in response to HFD-feeding.

### Choline- and DHA-Containing Phospholipids are Selectively Reduced in Livers from Female LKO Mice

The LD core is surrounded by a phospholipid monolayer consisting of primarily (>90%) phosphatidylethanolamine (PE) and phosphatidylcholine (PC) (45, 46). Given that female LKO mice accumulated many small LDs (**Fig. 2I**), we reasoned that phospholipid levels would also be affected in these mice. Utilizing an untargeted lipidomics approach by liquid chromatography-tandem mass spectrometry (LC-MS/MS), we observed similar levels of all major phospholipid species (PE, PC, phosphatidylserine [PS], phosphatidylinositol [PI], phosphatidylglycerol [PG], and phosphatidic acid [PA]) in male control and LKO mice (**Fig. S11A**). To our surprise, however, female LKO mice had decreased PC levels at the expense of both PE and PG in the liver (**Fig. 3A**). This lipidomics approach allows for semi-quantitation of lipid species, so we validated our untargeted analysis by quantifying the total mass of PC in livers from HFD-fed mice using a Folch-based extraction followed by an enzymatic assay containing PC standards. Consistently, female LKO mice had 9.9±1.74 (mg PC/g liver) relative to female control mice 13.9±3.02 (mg PC/g liver), a 28.3% reduction in the total mass of hepatic PC (**Fig. S11B**). The ratio of cellular PC to PE is critically important for maintaining LD growth and stability, and is tightly controlled by the phosphatidylethanolamine N-methyltransferase (PEMT) enzyme which facilitates PC synthesis from PE (45, 47, 48). We observed a significant 35% reduction in the ratio of total PC to PE (**Fig. 3B**), which is associated with reduced PEMT gene (1.5 fold) and protein (4-fold) levels (**Fig. 3C-E**) in female LKO mice relative to control mice. These data are consistent with reduced hepatic PC synthesis and accumulation of PE in female LKO mice.

**Figure 3.**
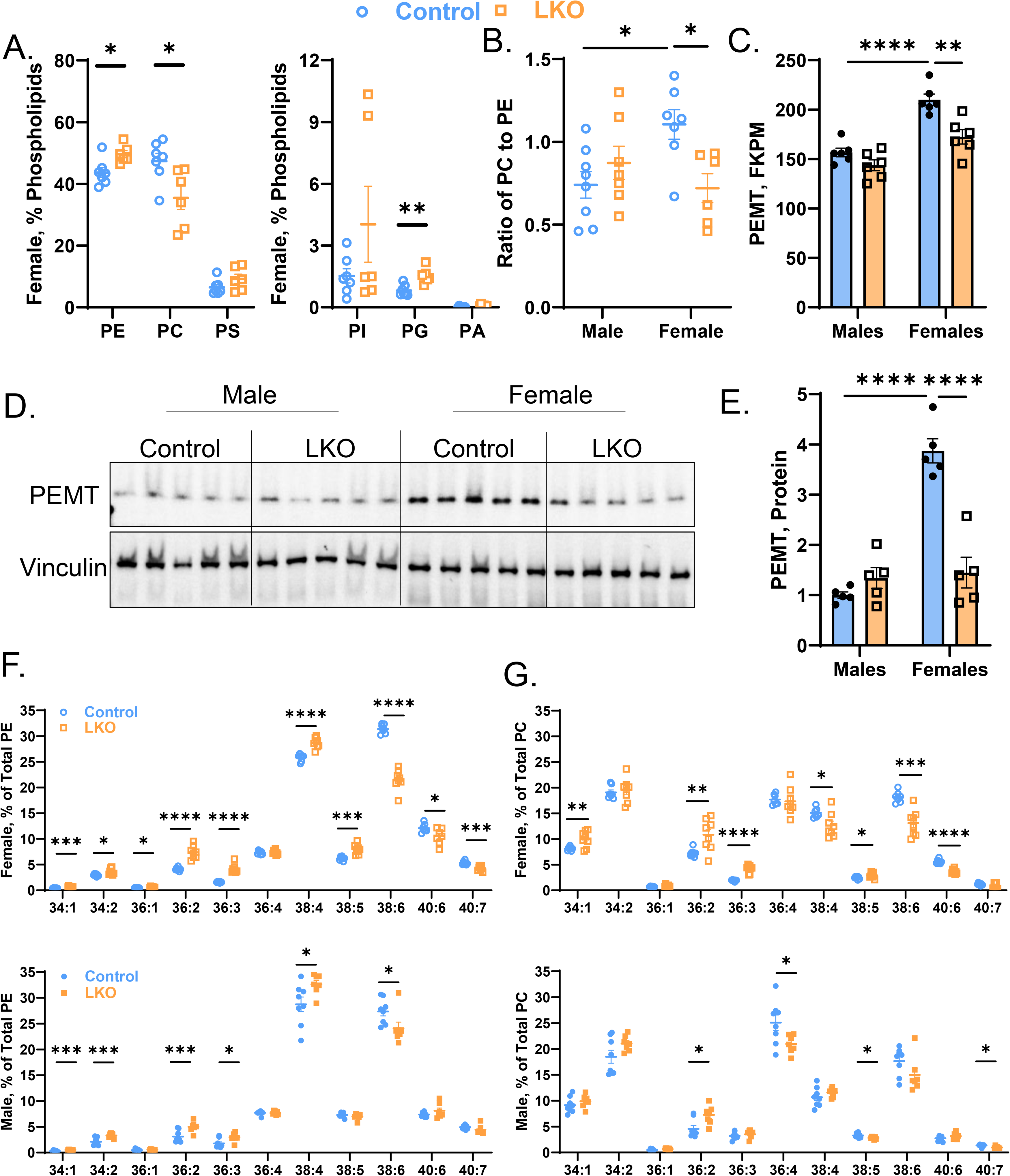
Female LKO Mice Have Increased PE and MUFA-Containing Phospholipids in the Liver. Male and female control and LKO mice were fed a HFD for 15-weeks. (**A**) Whole cell liver lysates were subjected to LC-MS/MS based lipidomics. Phospholipid species (PE, PC, PS, PI, PG, PA) displayed as percent (%) of total phospholipids (n=7-8). (**B**) A ratio of total PC to PE across the four groups (n=7-8). (**C**) Total read counts of *Pemt* from bulk RNA-sequencing data provided in Fig. 6 (FPKM=fragments per kilobase million; n=6). (**D**, **E**) Immunoblotting (**D**) followed by densitometry (**E**) for PEMT protein (n=5). Vinculin is used as a loading control. (**F**, **G**) The fatty acyl composition of PE (**F**) and PC (**G**) from female (top) and male (bottom) mice. All data are presented as % of total phospholipid (n=7-8). Significance was determined by an unpaired Student’s t-test (panels A, F, G) or by a two-way ANOVA with Tukey’s multiple comparison *post hoc* analysis (panels B, C, E). *P<0.05; **P<0.01; ***P<0.001; ****P<0.0001.

Aside from the total phospholipid pool, we next asked if CPT1a deficiency would alter the acyl-CoA composition of PE and PC. Male and female LKO mice exhibited increased MUFA (34:1, 34:2, 36:1, 36:2)-, ω6-PUFA-(36:3, 38:4)-, and EPA (38:5)-containing PE (**Fig. 3F**) and PC (**Fig. 3G**), while DHA-containing (38:6, 40:6) phospholipids were selectively reduced (**Fig. 3F, G**). Female LKO mice exhibited the most profound differences in PUFA-containing phospholipids, particularly the significant reductions in 38:6 PE (*P*=2.1E-08; **Fig. 3F**) and PC (*P*=2.9E-04; **Fig. 3G**) levels in the liver. Altogether, female LKO mice had increased hepatic PE levels at the expense of PC, which coincided with decreased PEMT protein levels in the liver. Moreover, both male and female LKO mice had increased MUFA-containing phospholipids at the expense of DHA-containing phospholipids, which tended to be more prominent in LKO females.

### MUFA-Containing Phospholipids Are Increased at the Expense of DHA-Containing Phospholipids in LD Fractions Collected From LKO Mice

Given the shift towards MUFA-containing PE and PC in whole cell lysates (**Fig. 3**), we asked whether these changes were also occurring within the LD phospholipid monolayer. We first isolated LDs from HFD-fed control and LKO mice by sucrose gradient centrifugation (17). Using non-LD and LD fractions, we immunoblotted for LD (perilipin 2 [PLIN2], PLIN5), mitochondria (voltage-dependent anion channel, VDAC), and cytosolic (glyceraldehyde 3-phopshate dehydrogenase, GAPDH) proteins. We confirmed the presence of PLIN2 and PLIN5 on LD-fractions, while these fractions were largely absent of the mitochondrial and cytosolic proteins, VDAC and GAPDH, respectively (**Fig. 4A**). Both PLIN2 and PLIN5 were more abundant on LD-fractions from female LKO mice suggesting more PLIN2 and 5 protein were coating these LDs in female LKO mice (**Fig. 4A**). We then extracted lipids from the LD fractions and quantified the acyl-CoA composition of PE and PC using a previously established targeted LC-MS/MS method (17). Consistent with our observation in whole liver lysates (**Fig. 3**), male and female LKO mice had increased MUFA-containing PC (**Fig. 4B**) and PE (**Fig. 4C**) at the expense of DHA-containing phospholipids, particularly 38:6 PE (**Fig. 4B, C**).

**Figure 4.**
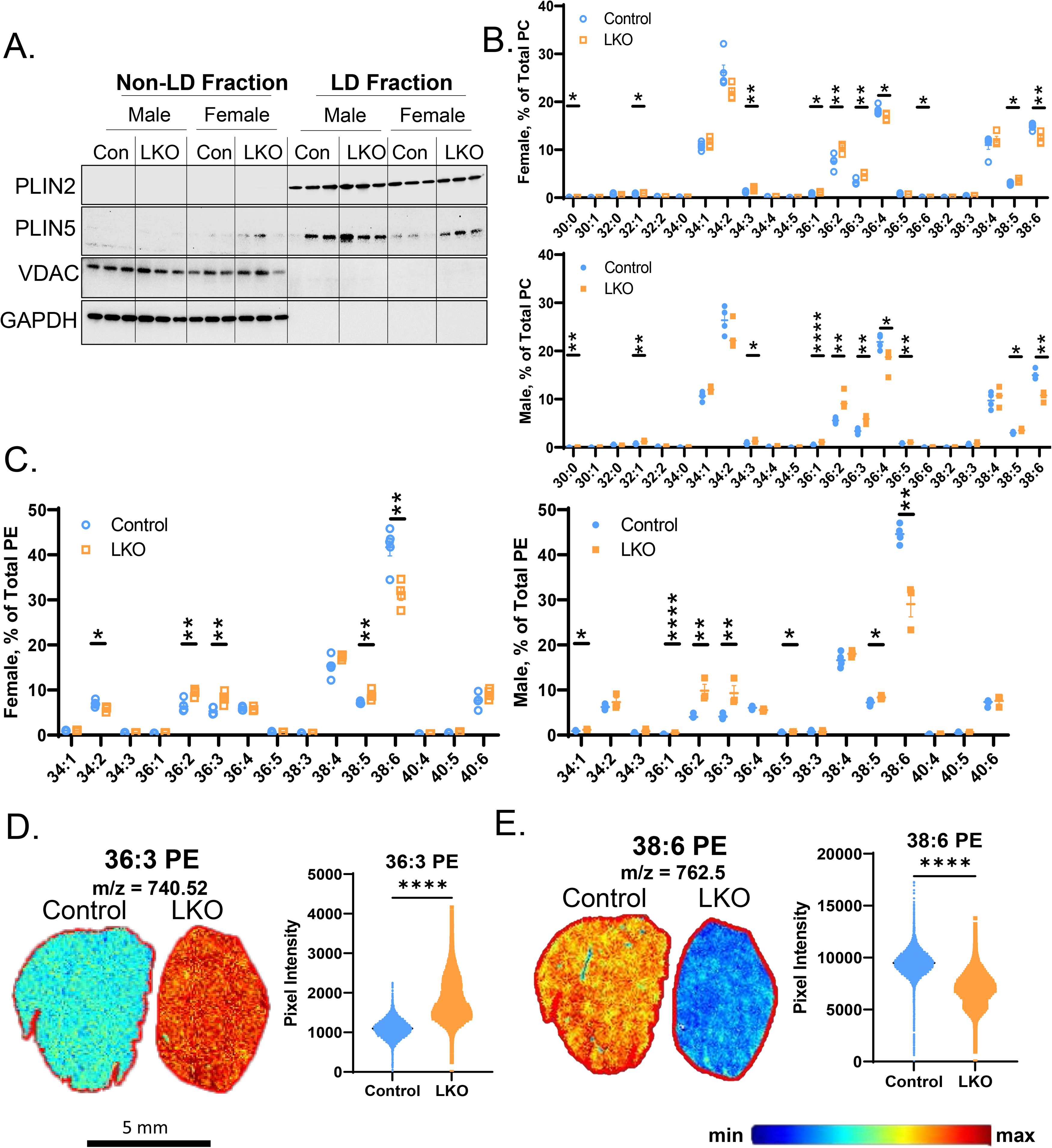
Lipid Droplets Isolated from LKO Mice Have Increased MUFA-Containing Phospholipids. Male and female control and LKO mice were fed a HFD for 15-weeks. (**A**) Non-LD and LD fractions were isolated from livers and immunoblotted for PLIN2, PLIN5, VDAC, and GAPDH (n=3). (**B**, **C**) Lipid droplet fractions were subjected to targeted semi-quantification of PC (**B**) and PE (**C**; n=3-5). (**D**, **E**) MALDI-MSI was completed on whole livers from female control and LKO mice (n=4). A representative spatial map and pixel intensity are shown for 36:3 PE (**D**) and 38:6 PE (**E**). Significance was determined by unpaired Student’s t-tests. *P<0.05; **P<0.01; ***P<0.001; ****P<0.0001.

Phospholipids exhibit varying spatial intensities across the periportal-pericentral axis in response to genetic- and dietary manipulation (42, 49). While periportal hepatocytes are thought to undergo greater FAO rates, they also contain more arachidonic acid- and DHA-containing phospholipids (e.g. 38:6 PC) (42, 49). Therefore, we asked whether the panlobular microvesicular steatosis observed in female LKO mice may lead to changes in the spatial distribution of these key MUFA- and PUFA-containing phospholipids. Utilizing MALDI-MSI technology, we observed that female LKO mice had significantly increased 36:3 PE intensity, while the intensity for the DHA-containing phospholipid, 38:6 PE, was significantly reduced across the entire liver lobule (**Fig. 4D**). While there was a tendency for 38:6 PE to exhibit spatial localization across the liver lobule in control mice, the intensity was dramatically reduced across the entire lobule in both male and female LKO mice (**Fig. 4D, S12**). A compilation of all MALDI-MSI spatial maps for 36:3 PE and 38:4 PE across males and females has been provided in **Fig. S12**. Collectively, whole cell lysates and LDs isolated from LKO mice favor MUFA-containing at the expense of DHA-containing PE and PC, which appears to be pan-lobular rather than spatially localized across the periportal-pericentral axis.

### PPAR Signaling and PUFA Biosynthetic Genes are Upregulated in LKO Mice

To attempt to tease out the mechanisms explaining the sexually dimorphic response to *Cpt1a* deficiency, we completed bulk RNA-sequencing on whole livers from HFD-fed male and female mice. We identified 577 (male) and 748 (female) genes that met pre-determined statistical (p-adjusted value ≤0.05) and effect size cut-offs (Log2 normalized fold-change [FC] of ±0.58). Partial least squares-discriminant analysis showed that the transcriptome of LKO mice was profoundly different from that of WT mice (both in males and females), accounting for up to 12.16% on PC2 (**Fig. S13A**). Moreover, sex alone accounted for up to 24.61% of the total variation on PC1 between male and female mice (**Fig. S13A**). The most downregulated gene in both male and female LKO mice was *Cpt1a*, while several other genes involved in *de novo* lipogenesis (*Srebf1*, *Acly*) and carbohydrate metabolism (*Khk*, *Tkfc, Pklr, Pygl*) were significantly downregulated compared to control mice (**Fig. 5A, B**). Laminin subunit beta-3 (*Lamb3*), which is involved in Akt-mediated tumorigenesis (50), was the most significantly upregulated (Log_2_ FC=2.39; Padj=2.05E-73) gene in response to LKO in male mice. In female LKO mice, the most significantly upregulated gene (Log_2_ FC=3.61; Padj=1.18E-58) was cell death-inducing DFFA-like effector C (*Cidec*), a gene encoding a LD-tethering protein involved in the fusion and growth of LDs (51). Genes encoding proteins that influence LD turnover (*Plin2*, *Hilpda*, *G0S2*) and fatty acid metabolism (*Acot2*, *Cyp4a14*, *Cyp4a12*) were collectively upregulated in LKO mice across both sexes (**Fig. 5A, B**). Notably, inflammatory genes (*Ly6d*, *Mmp12*, *Cxcl2*) were selectively upregulated only in female LKO mice (**Fig. 5B**), likely contributing to the worsened liver injury in these animals (**Fig. 2D**).

**Figure 5.**
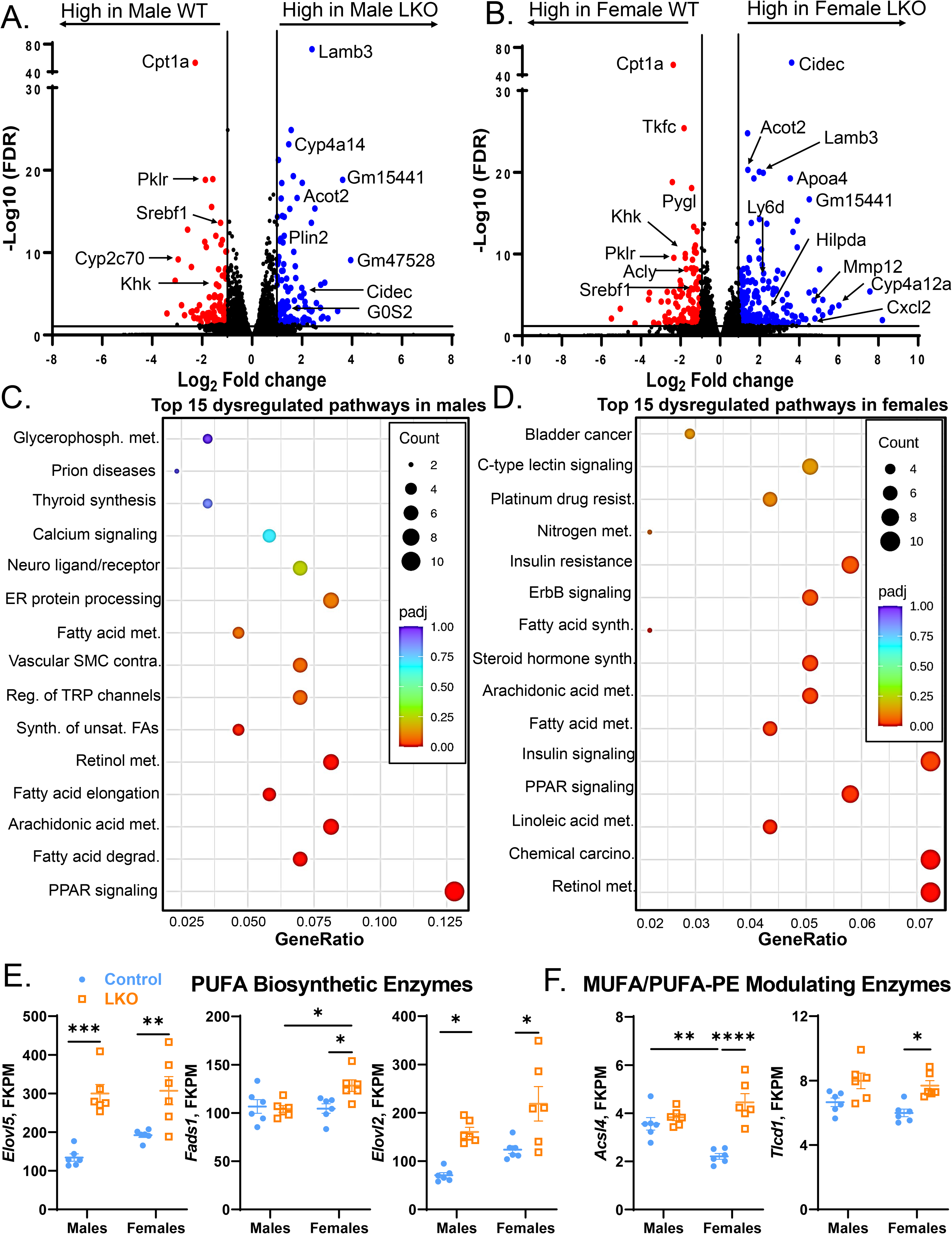
PPAR Signaling and Lipid Droplet Genes Are Elevated in LKO Mice. Bulk RNA sequencing was completed on the livers of male and female HFD-control and LKO mice. (**A**, **B**) Volcano plots in males (**A**) and females (**B**) highlighting all genes increased (in blue) or decreased (in red) with *Cpt1a*-deficiency. The horizontal black bar denotes the significance cutoff FDR=0.05. The vertical black bars denote a minimal threshold for the effect size of 1.5 (Log2 fold change=±0.58). (**C**, **D**) KEGG analysis dot plots for the top 15 dysregulated pathways in WT and LKO male (**C**) and female (D) mice. (**E**, **F**) Total read counts (in FPKM) for genes involved in the biosynthesis of PUFAs (**E**; *Elovl5*, *Fads1*, *Elovl2*) and fatty acyl remodeling of phospholipids (**F**; *Acsl4*, *Tlcd1*). Significance was determined by an unpaired Student’s t-test (panels E, F). *P<0.05; **P<0.01; ***P<0.001; ****P<0.0001.

To determine pathways differentially affected by *Cpt1a*-deletion in the liver, we completed a Kyoto Encyclopedia of Genes and Genomes (KEGG) enrichment analysis for both male and female mice. The top 5 dysregulated pathways comparing male control and LKO mice were related to peroxisome proliferator-activated receptor (PPAR) signaling, fatty acid degradation, arachidonic acid metabolism, fatty acid elongation, and retinol metabolism (**Fig. 5C**). Similarly, in female mice, the top 5 pathways affected were related to retinol metabolism, chemical carcinogenesis, linoleic acid metabolism, and PPAR and insulin signaling (**Fig. 5D**). Since both male and females had enrichment of PPAR signaling pathways, we measured individual PPAR isoforms (α, δ, γ1, γ2) using normalized read counts from RNA-sequencing and by qPCR. Along with impairments in long chain FAO with *Cpt1a*-deficiency, RNA levels of the major transcription factor controlling FAO gene transcription, PPARα, was elevated in both male and female LKO mice (significance by genotype = 0.0005; **Fig. S13B**). Moreover, only female LKO had significant elevations in PPARγ1 (**Fig. S13B**), while PPARδ and PPARγ2 were unaffected.

Aside from PPAR-signaling, the KEGG enrichment analysis also identified several pathways related to PUFA metabolism. The synthesis of PUFAs, such as DHA, are mediated by a series of reactions that involve the desaturation and elongation of substrate lipids. The enzymes catalyzing these reactions include members of the fatty acid desaturase (*Fads1, 2*) and elongase gene families (*Elovl2, 5*). We examined normalized read counts of PUFA biosynthetic genes in livers collected from female control and LKO mice fed a HFD. Despite significant reductions in DHA-containing phospholipids in both whole liver lysates (**Fig. 3F, G**) and in LDs (**Fig. 4B, C**), the biosynthetic enzymes *Elovl5*, *Fads1*, and *Elovl2* were transcriptionally elevated in LKO mice relative to control mice (**Fig. 5E; S13C**).

Given the shift from DHA to MUFA-containing phospholipids, we also measured RNA levels of genes known to regulate the acyl-CoA composition of phospholipids. The long-chain acyl-CoA synthetase 4 (*Acsl4*) gene facilitates the synthesis of long-chain PUFA-CoAs for esterification into phospholipids, albeit with a preference for arachidonic acid and ω3-PUFAs (52). In female LKO mice, *Acsl4* gene expression was significantly elevated as compared to control mice (**Fig. 5F**). No changes were observed in RNA levels of 1-acyl-*sn*-glycerol-3-phosphate acyltransferase (*Agpat3*) or lysophosphatidylcholine acyltransferase (*Lpcat3*), two genes shown to facilitate DHA (53)- and arachidonic acid (54)-incorporation into PE lipids, respectively (**Fig. S13C**). The *sn*-1 position of PE contains saturated fatty acids (SFAs) or MUFAs, and recent work identified transmembrane proteins containing TRM-Lag1p-CLN8 domains 1 and 2 (*Tlcd1*, *2*) as selective inducers of MUFA incorporation into *sn*-1 position of PEs (55). Consistent with previous reports that *Tlcd1* and *Tlcd2* increase MUFA-PEs and drive worsened liver injury in mice (55), female LKO mice exhibited elevated *Tlcd1* RNA levels as compared to female control mice (**Fig. 5F; S13C**). Altogether, male and female LKO mice respond to *Cpt1a*-deletion by upregulating PPARα signaling (and PPARγ1 in females) and its downstream targets, some of which are involved in LD hydrolysis (*Cidec*, *Plin2*, *G0s2*; **Fig. 5A-D**) (56, 57). Moreover, genes involved in PUFA biosynthesis (*Elovl5*, *Fads1*, *Elovl2*) and in MUFA incorporation into PE (*Tlcd1*) were also elevated in female LKO mice (**Fig. 5E**).

### PKA Signaling and Triglyceride Hydrolysis is Impaired in LKO Mice

To identify altered signal transduction that might influence downstream lipid and gene expression changes observed in LKO mice, we quantified the activity of 144 serine/threonine kinases (STK) in liver samples from HFD-fed male and female mice using PamGene technology in real-time (22, 25). We determined individual substrate phosphorylation from the PamGene analysis into heatmaps showing the top peptides that exhibited differential phosphorylation compared to controls (**Fig. S14A, B**). We then used bioinformatics to identify upstream kinases responsible for the observed difference in phosphorylation across all peptides. The activity of protein kinase A (PKA) was one of the most repressed STKs in LKO mice in both male and female mice. Peacock plots (**Fig. 6A**; **S15A**), which show the confidence of our bioinformatics analysis relative to the number of random sampling over 2000 iterations and overall kinase activity across all PKA-peptides (22, 25) (**Fig. 6B; S15B**), demonstrate with high confidence that PKA activity was impaired in both LKO groups. A major role of PKA during fasting is to phosphorylate proteins involved in the lipolytic machinery of LDs, including perilipins and hormone sensitive lipase (HSL) in adipocytes. Individual peptides were then compiled in a waterfall plot across male (**Fig. 6C**) and female mice (**Fig. S15C**), and we discovered the hormone sensitive lipase (HSL) peptide (encoding serine 950, 951, and threonine 955) and the β2-adrenergic receptor (β2-AR) peptide (encoding serine 345 and 346) was significantly reduced in male LKO but not female LKO mice (**Fig. 6C; S15C**). We then immunoblotted for perilipins (PLIN2, PLIN5), lipases (adipose triglyceride lipase, ATGL; HSL, monoacylglycerol lipase, MGL), and lipase co-regulators (comparative gene identification 58, CGI-58; G0/G1 switch 2, G0S2) known to regulate LD lipolysis (**Fig. 6D**). Consistent with changes at the RNA level (**Fig. 5**), both male and female LKO mice had elevated PLIN2 and PLIN5 protein (**Fig. 6D, E**). Total protein levels of ATGL, HSL, and MGL were largely unchanged in male LKO mice; however, female LKO mice had elevated ATGL (∼30-fold) and MGL (∼6-fold) protein levels, relative to female control mice (**Fig. 6D, E**). Protein levels of the PPARα-target gene (58) and potent inhibitor of ATGL (59), G0S2, was also elevated (4-fold) in male and female LKO mice (**Fig. 6D, E**). Consistent with impaired PKA signaling and increased protein levels of PLIN 2, 5, and G0S2, triglyceride hydrolysis was reduced in mice lacking *Cpt1a* in the liver (**Fig. 6E**).

**Figure 6.**
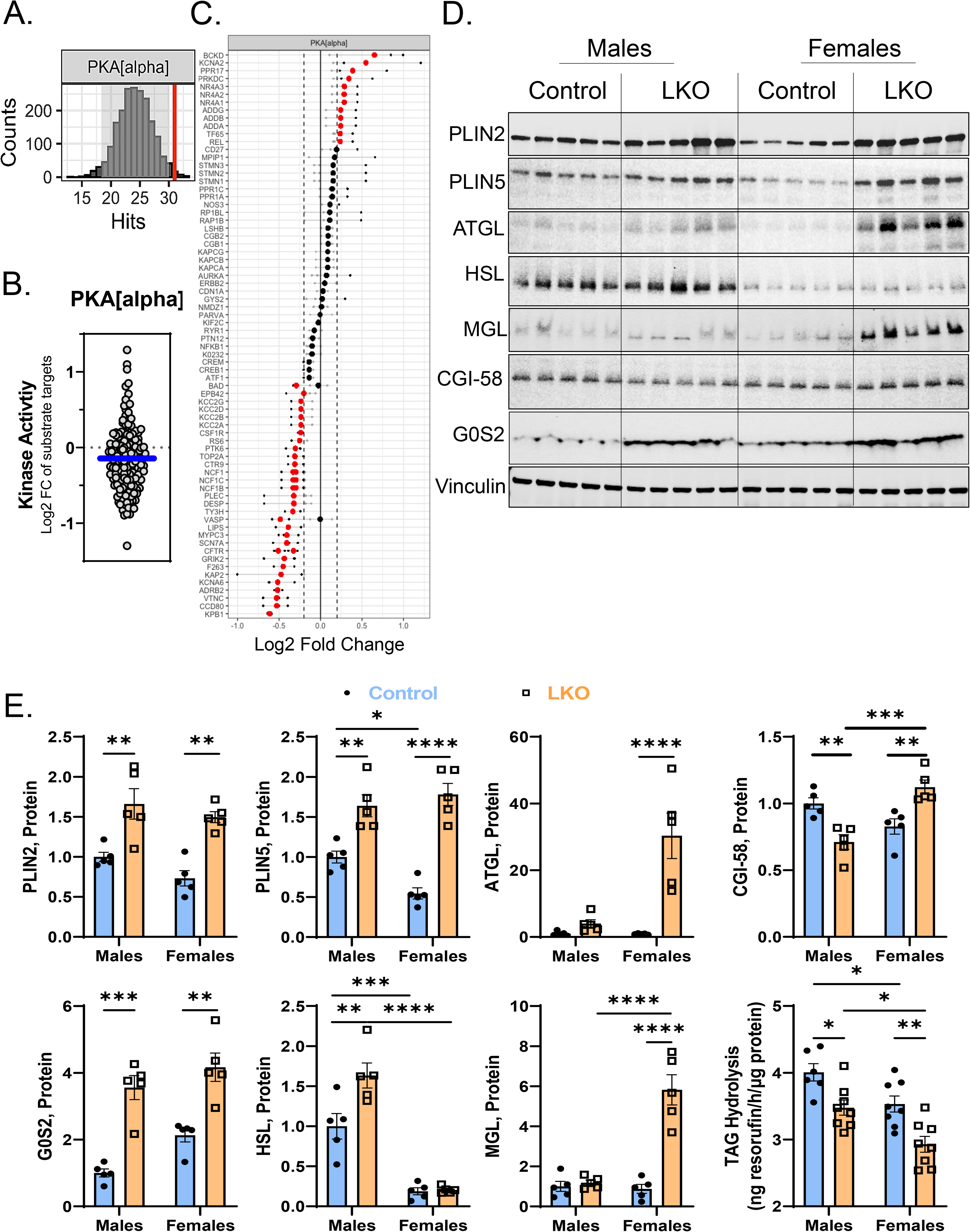
PKA Signaling and Triglyceride Hydrolysis Are Impaired in LKO Mice. A serine-threonine kinome analysis was completed on pooled liver samples from male and female HFD-control and LKO mice. A total of 6 mice per sex and genotype were pooled and ran in triplicate on the PamGene PamStation. (**A**-**C**) PKA peacock (**A**), MEOW (**B**), and substrate waterfall plots (**C**) are presented comparing male control and LKO mice. The waterfall plot shows all known peptides that are phosphorylated by PKA and the extent by which their phosphorylation, in real-time, is increased or decreased (in red) with *Cpt1a*-deficiency. (**D**, **E**) Immunoblotting (**D**) followed by densitometry (**E**) for proteins (PLIN2, PLIN5, ATGL, HSL, MGL, CGI-58, G0S2) associated with regulating the size and turnover of LDs (n=5). Vinculin is used as a loading control. (**F**) Triglyceride hydrolysis was completed using liver lysates from HFD-fed control and LKO mice using resorufin ester as substrate (n=8). Significance was determined by a two-way ANOVA with Tukey’s multiple comparison *post hoc* analysis. *P<0.05; **P<0.01; ***P<0.001; ****P<0.0001.

## DISCUSSION

Here we report a novel role for CPT1a controlling sexually dimorphic SLD. First, we show that ablation of CPT1a in the liver of male mice promotes modest (although not significant) LD accumulation in periportal hepatocytes while total hepatic lipids (triglycerides, cholesterol) across the lobule were not different compared to male control mice. In fact, male LKO mice were protected from liver dysfunction (via ALT levels) compared to control mice. However, female control mice were largely protected from diet-induced liver dysfunction, which was completely lost in female mice lacking CPT1a in the liver. The exacerbation of SLD in female LKO mice was characterized by the accumulation of LDs within periportal and pericentral hepatocytes, consistent with significant elevations in triglycerides, cholesterol, and circulating ALT levels. Estrogen has been shown to stimulate FAO (60) and *Cpt1a* gene transcription via the estrogen receptor α (61). It is plausible that estrogen mediates its hepatoprotective properties through a CPT1a-dependent FAO mechanism. However, we did not detect differences in CPT1a RNA or protein levels across male and female mice. Given that CPT1a activity is subjected to allosteric regulation by malonyl-CoA, future studies should determine if CPT1a enzymatic activity is regulated in a sexually dimorphic manner.

Next, we show that *Cpt1a* deficiency in female mice is associated with alterations in the total phospholipid pool from PC to PE, as well as the acyl-CoA composition of these two major phospholipids. The PEMT enzyme facilitates the methylation of PE to generate PC, accounting for ∼30% of total liver PC biosynthesis in mammalian hepatocytes, while the remaining ∼70% of PC is synthesized by the conversion of choline to PC (62, 63). PEMT confers specificity for PUFA-containing PE as a substrate for PC synthesis, while MUFA-containing PCs tend to be derived largely from the CDP-choline pathway (62, 64). A reduction in PEMT has been shown to deplete hepatic and plasma DHA-containing PC (65), promote steatosis, inflammation, and fibrosis in mouse models of SLD (66-68), and associate with an increased risk of worsened liver dysfunction in humans (69, 70). Notably, PEMT is responsive to estrogen due to three evolutionarily conserved estrogen regulatory motifs within the promoter (71), and loss of ovarian sex hormones by ovariectomy promotes SLD and reduces *Pemt* gene expression in the liver (72). These publications largely support our findings where we show female LKO lose their protection from diet-induced SLD, which associates with lower PEMT protein and PC:PE ratios compared to female control mice. Further, male and female LKO mice have reduced DHA-containing PC and PE, indicating other mechanisms independent of PEMT are likely driving the shifts in acyl-CoA phospholipid composition.

To further understand how *Cpt1a* shapes phospholipid metabolism, we measured the expression of genes involved in the PUFA-biosynthetic pathway. Despite reductions in DHA-containing phospholipids, we observed significant increases in the fatty acid elongase gene family (*Elovl5*, *Elovl2*). The ELOVL2 enzyme facilitates the elongation of 20- and 22-carbon PUFAs (73), while ELOVL5 has a broader fatty acid specificity (74). Depletion of DHA, through genetic deletion of ELOVL5 or AGPAT3, drives SLD via activation of sterol response element binding transcription factor-1 (SREBP1)-mediated *de novo* lipogenesis (53, 74). On the contrary, supplementation of DHA strongly suppresses SREBP1 activation (74, 75). Given this negative feedback loop between DHA and SREBP1, we expected an increase in *Srepf1* and its downstream target genes, but *Srebf1* was consistently downregulated across male and female LKO mice, as were its canonical target genes *Acly* and *Fasn*. Given that this pathway is also positively regulated by insulin levels (76), the low insulin levels observed in LKO mice likely contribute to the reductions in lipogenic gene expression observed in these animals.

Aside from reduced SREBP1-mediated lipogenesis, our studies show that loss of CPT1a in the liver of male and female mice stimulates PPAR-signaling. We found that the PPARα isoform, which transcriptionally regulates genes involved in peroxisomal biogenesis and fatty acid metabolism, was elevated across male and female LKO mice. This is consistent with other reports in skeletal muscle showing compensatory increases in peroxisomal FAO in response to impaired mitochondrial FAO (77). Further, an increase in peroxisome and LD number can be driven by MUFA supplementation, which in turn elevates the phospholipid MUFA:PUFA ratio and reduces the capacity for lipid oxidation (78). On the contrary, impaired peroxisomal β-oxidation promotes the accumulation of PUFA-CoAs and represses large LD formation (79). Therefore, enhanced PPARα signaling in LKO mice may accelerate DHA catabolism in peroxisomes leading to a shift in MUFA:PUFA-containing phospholipids in the LD monolayer, which results in smaller and more stable LDs.

Another key observation in female LKO mice was the significant elevation in *Pparγ1* RNA, which coincided with significant upregulation of *Cidec*, an established PPARγ target gene (80). While PPARγ has historically been studied in adipose tissue, laboratories have shown that gain- and loss-of-function of PPARγ in the liver exacerbates and ameliorates diet-induced liver dysfunction (81, 82), which is mediated, in part, by *Cidec* (80). CIDEC is an LD-associated protein that facilitates the transfer of neutral lipids from smaller to larger LDs, thereby inhibiting lipolysis and controlling the expansion of these organelles (51). Counterintuitive to its role in LD expansion, forced expression in the liver phenocopies the histopathology observed in female LKO mice – the accumulation of small LD vacuoles throughout the liver lobule (80). Future studies are required to fully understand the role of *Cidec* in mediating microvesicular steatosis, a phenotype observed in individuals with impaired FAO (83).

To assess which signaling pathways might be altered with CPT1a ablation, we performed an unbiased characterization of STK activity in male and female mice. This analysis revealed significant suppression of cyclic adenosine monophosphate (cAMP)-dependent protein kinases, including protein kinase X (PRKX) and PKA in LKO mice. PKA functions as a heterotetramer which consists of 2 regulatory and 2 catalytic subunits, one of which is PRKX. An increase in intracellular cAMP, as observed during fasting, results in a conformational change of the protein leading to activation and subsequent phosphorylation of lipolytic proteins, including PLIN5 (84) and HSL (85), to drive LD catabolism. Inhibition of this pathway further promotes the accumulation of LDs in mice (86). Consistent with reduced PKA activity, we observed significant reductions in triglyceride hydrolysis from liver lysates of LKO mice. This reduction in hydrolysis can be attributed to loss of PKA activity and an increase in protein of known lipase inhibitors (PLIN2, PLIN5, G0S2), despite significant elevations in lipases themselves (ATGL, MGL). Collectively, a reduction in the PKA-mediated liberation of fatty acids from LDs likely serves as a negative feedback loop to protect hepatocytes from lipotoxicity during periods of impaired FAO.

It is important to note that some of the results highlighted here are not in complete agreement with a previous study showing the impact of liver-specific CPT1a-deficiency on SLD in mice. The manuscript by Sun et al. showed male CPT1a LKO mice had significantly lower body weight, and improved glucose and insulin tolerance in response to a 45% kcal from fat HFD (34). While we were unable to replicate the lowering of body weight in LKO mice, we did observe significant reductions in circulating insulin levels in male LKO mice fed a 60% kcal from fat HFD, consistent with improvements in insulin sensitivity. Other notable differences include the observation that male LKO mice had significant elevations in hepatic triglycerides in response to HFD feeding (34). In our studies, we did observe slight periportal steatosis in male LKO mice; however, hepatic lipid data from male mice were variable and insignificant compared to male control mice. In the previous report, female mice were not included, and it is unclear if mice were necropsied in the fed or fasted state (34). Overall, while many of the phenotypes described by Sun et al. (34) was replicated here; a few distinct differences were noted and should be considered moving forward.

In this study, we show that female mice are dependent on CPT1a-mediated FAO for protection from HFD-induced SLD. The exacerbation of SLD observed with CPT1a deficiency in female mice is associated with significant reductions in hepatic PC synthesis via the PEMT-dependent pathway. Modulation of PPARα and PKA signaling pathways may serve as compensatory mechanisms to protect hepatocytes from lipotoxicity in the face of impaired FAO. Future studies should consider sexually dimorphic responses to FAO when designing more personalized therapeutics for the management of SLD.

## ABBREVIATIONS

CPT1a: Carnitine Palmitoyltransferase 1a
SLD: Steatotic Liver Disease
PKA: Protein Kinase A
LD: Lipid droplet
DHA: Docosahexaenoic acid
MUFA: Monounsaturated fatty acid
PUFA: Polyunsaturated fatty acid
MALDI-MSI: Matrix-assisted laser desorption ionization for mass spectrometry imaging
FAO: Fatty acid oxidation
ALT: Alanine aminotransferase
HFD: High fat diet
LFD: Low fat diet
PPAR: Peroxisome proliferator-activated receptor

## DATA AVAILABILITY

All data are contained in the entirety of this manuscript. Additional information and requests for resources and reagents should be directed to and fulfilled by the Lead Contact, Robert N. Helsley. Bulk RNA-sequencing data have been deposited in GEO under accession code GSE225769.

## ACKNOWLEDGEMENTS

We would like to acknowledge and thank the laboratory of Dr. Peter Carmeliet (Katholieke Universiteit Leuven) for sharing the *Cpt1a* floxed mice. We would also like to thank Drs. René Jacobs (University of Alberta) and Jun Liu (Mayo Clinic) for sharing their respective PEMT and G0S2 antibodies.

## AUTHOR CONTRIBUTIONS

**Mikala M. Zelows**: Conceptualization, Methodology, Validation, Formal analysis, Investigation, Data Curation, Writing – Original Draft, Writing – Review & Editing, Visualization. **Corissa Cady**: Investigation, Writing – Review & Editing. **Nikitha Dharanipragada**: Methodology, Formal analysis, Investigation, Data Curation, Writing – Review & Editing. **Anna Mead**: Methodology, Formal analysis, Investigation, Data Curation, Writing – Review & Editing. **Zachary Kipp**: Methodology, Software, Validation, Formal analysis, Investigation, Data Curation, Writing – Review & Editing. **Evelyn Bates**: Methodology, Software, Validation, Formal analysis, Investigation, Data Curation, Writing – Review & Editing. **Venkateshwari Varadharajan**: Formal analysis, Data Curation, Writing – Review & Editing. **Rakhee Banerjee**: Formal analysis, Data Curation, Writing – Review & Editing. **Se-Hyung Park**: Investigation, Writing – Review & Editing. **Nathan R. Shelman**: Formal analysis, Data Curation, Writing – Review & Editing. **Harrison A. Clarke**: Formal analysis, Data Curation, Writing – Review & Editing. **Tara R. Hawkinson**: Formal analysis, Data Curation, Writing – Review & Editing. **Terrymar Medina**: Formal analysis, Data Curation, Writing – Review & Editing. **Ramon C. Sun**: Conceptualization, Methodology, Software, Validation, Formal analysis, Investigation, Resources, Data Curation, Writing – Original Draft, Writing – Review & Editing, Visualization, Supervision. **Todd A. Lydic**: Formal analysis, Investigation, Data Curation, Writing – Original Draft, Writing – Review & Editing. **Terry Hinds Jr.**: Methodology, Software, Validation, Formal analysis, Investigation, Resources, Data Curation, Writing – Original Draft, Writing – Review & Editing, Supervision. **J. Mark Brown**: Data Curation, Methodology, Validation, Formal analysis, Resources, Data Curation, & Funding acquisition**. Samir Softic**: Resources, Writing – Original Draft, Writing – Review & Editing, Supervision, Project administration, Funding acquisition. **Gregory A. Graf**: Conceptualization, Resources, Data Curation, Writing – Original Draft, Writing – Review & Editing, Supervision, Project administration, Funding acquisition. **Robert N. Helsley**: Conceptualization, Data Curation, Methodology, Validation, Formal analysis, Investigation, Resources, Data Curation, Writing – Original Draft, Writing – Review & Editing, Visualization, Supervision, Project administration, Funding acquisition.

## TABLES WITH TITLES AND LEGENDS

**Supplemental Table 1.** STK Peptide Substrates

**Supplemental Table 2.** STK Scores and Rankings Male LKO versus Male Control

**Supplemental Table 3.** STK Scores and Rankings Female LKO versus Female Control

**Supplemental Table 4.** Mouse primers used for real-time PCR.

**Supplemental Table 5.** Antibodies used for immunoblotting.

## SUPPLEMENTAL INFORMATION TITLES AND LEGENDS

**Supplemental Figure 1. CPT1a Deletion in the Liver Reduces Fasting Ketones In Response to LFD-Feeding.** (**A**) An overview of the experimental design. (**B**-**D**) Liver CPT1a RNA and protein levels were measured by qPCR (**B**) and western blot (**C**) followed by densitometry (**D**), respectively (n=5). Vinculin is used as a loading control for immunoblotting. (**E**) RNA levels of *Cpt1b* were measured in livers of LFD-fed control and LKO mice by qPCR (n=8). For all qPCR analyses, housekeeping genes *Tbp* and *Hprt* were averaged and used for normalization. (**F**) β-hydroxybutyrate levels were measured from plasma of fasted mice fed LFD (n=7-8). Significance was determined by two-way ANOVA with Tukey’s multiple comparison *post hoc* analysis. *P<0.05; **P<0.01; ***P<0.001; ****P<0.0001.

**Supplemental Figure 2. CPT1b RNA and Protein Levels Are Low in Liver.** (**A**) Total read counts for *Cpt1a* and *Cpt1b* in FPKM (fragments per kilobase million) from bulk RNA-sequencing data on HFD-fed control male livers (n=6). (**B**) Total read counts for *Cpt1b* in male control and LKO mice fed a HFD (n=6). (**C**) A comparison of CPT1b protein expression from liver and brown adipose tissue (BAT) of female control mice fed HFD. Vinculin was used as a loading control. Significance was determined by an unpaired Student’s t-test. *P<0.05; ****P<0.0001.

**Supplemental Figure 3. Liver-specific CPT1a Deletion Has No Effect on Body Weight at Baseline and Response to LFD-Feeding.** (**A**-**C**) Baseline body weights (**A**), percent (%) fat (**B**) and lean (**C**) mass in control and LKO mice at 6-8 weeks of age (n=15-20). (**D**, **E**) Percent body weight (**D**) and fat mass throughout the study in response to LFD-feeding (n=8-12). Significance was determined by two-way ANOVA with Tukey’s multiple comparison *post hoc* analysis. ****P<0.0001.

**Supplemental Figure 4. LFD-Fed Control and LKO Mice Exhibit Similar Glucose Tolerance and Insulin Sensitivity.** After 10- and 12-weeks of LFD-feeding, male and female control and LKO mice underwent intraperitoneal glucose (IPGTT) and insulin tolerance tests (IPITT), respectively. (**A**-**D**) Glucose levels in response to an exogenous bolus of glucose (**A**) and insulin (**C**), as well as their area under the curves, were quantified (**B**, **D**; n=8-12). Significance was determined by two-way ANOVA with Tukey’s multiple comparison *post hoc* analysis. The ANOVA results have been provided. **P<0.01.

**Supplemental Figure 5. HFD-Fed Male LKO Mice Have Lower Circulating Insulin Levels and Improved HOMA-IR.** After 10- and 12-weeks of HFD-feeding, male and female control and LKO mice underwent intraperitoneal glucose (IPGTT) and insulin tolerance tests (IPITT), respectively. (**A**-**C**) Fasting glucose (**A**; n=8-15), insulin (**B**; n=7-8), and HOMA-IR (**C**; homeostatic model assessment for insulin resistance) was calculated (n=7-8). (**D**, **E**) Glucose levels in response to an exogenous bolus of glucose (**D**) and insulin (**E**), as well as their area under the curves were quantified (**D**, **E**; n=8-13). Significance was determined by two-way ANOVA with Tukey’s multiple comparison *post hoc* analysis. The ANOVA results have been provided. ***P<0.001; ****P<0.0001.

**Supplemental Figure 6. Female LKO Mice Consume More Calories than Control Mice.** After 7- weeks of HFD-feeding, male and female control and LKO mice were subjected to indirect calorimetry. At Day 4, the mice were fasted overnight. (**A**-**F**) Cumulative caloric intake (in kcal) over the course of 5- days (**A**, **D**) or over the course of 3-days (**B**, **E**) and throughout the light/dark cycle (**C**, **F**) in male (**A-C**) and female (**D-F**) mice. Significance was determined by two-way ANOVA with Tukey’s multiple comparison *post hoc* analysis (n=4). The ANOVA results have been provided. *P<0.05.

**Supplemental Figure 7. Female LKO Mice are More Active than Control Mice.** After 7-weeks of HFD-feeding, male and female control and LKO mice were subjected to indirect calorimetry. At Day 4, the mice were fasted overnight. (**A**-**F**) Cumulative activity (in meters) over the course of 5-days (**A**, **D**) or over the course of 3-days (**B**, **E**) and throughout the light/dark cycle (**C**, **F**) in male (**A-C**) and female (**D-F**) mice. Significance was determined by two-way ANOVA with Tukey’s multiple comparison *post hoc* analysis (n=4). The ANOVA results have been provided. *P<0.05; ****P<0.0001.

**Supplemental Figure 8. Female LKO Mice Expend More Energy than Control Mice.** After 7-weeks of HFD-feeding, male and female control and LKO mice were subjected to indirect calorimetry. At Day 4, the mice were fasted overnight. (**A**, **B**) Respiratory exchange ratio (RER) over the course of 3 days and throughout the light/dark cycle in male (**A**) and female (**B**) mice. (**C**, **D**) Energy expenditure (kcal/h) over the course of 3 days and throughout the light/dark cycle in male (**C**) and female (**D**) mice. Significance was determined by two-way ANOVA with Tukey’s multiple comparison *post hoc* analysis (n=4). The ANOVA results have been provided. **P<0.01; ***P<0.001.

**Supplemental Figure 9. Female LKO Mice Accumulate More Hepatic Cholesterol in Response to Fasting-Induced Steatosis.** Male and female control and LKO mice were fed a LFD for 15-weeks and were necropsied after a 16 hour fast. (**A**-**C**) Liver weights normalized to body weight (**A**; LW:BW, %; n=8-11), and hepatic triglycerides (**B**; n=8) and total cholesterol (**C**; n=8) levels were quantified enzymatically. (**D**) Serum ALT levels were quantified (n=8). Significance was determined by two-way ANOVA with Tukey’s multiple comparison *post hoc* analysis. *P<0.05; **P<0.01; ****P<0.0001.

**Supplemental Figure 10. Female LKO Mice Tend to Have a Greater Frequency of Small Lipid Droplets in Pericentral Hepatocytes than Control Mice.** Male and female control and LKO mice were fed a HFD for 15-weeks and were necropsied after a 16 hour fast. (**A**, **B**) Total lipid droplet diameters were quantified in a blinded fashion and presented as percent (%) frequency in periportal (**A**) and pericentral (**B**) hepatocytes. The frequency classes were broken down as follows (μm): 0-5, 5-10, 10-15, 15-20, 20-25, 25-30, 30-35, 35-40, 40-45, 45-50, 50-150.

**Supplemental Figure 11. HFD-Fed Female LKO Mice Have Reduced Hepatic PC.** (**A**) Hepatic phospholipid levels measured by tandem-MS in HFD-fed male mice (n=7-8). (**B**) Hepatic PC levels measured enzymatically across male and females (n=7-8). Significance was determined by two-way ANOVA with Tukey’s multiple comparison *post hoc* analysis. **P<0.01.

**Supplemental Figure 12. Spatial Intensities for 36:3 and 38:6 PE.** (**A**) MALDI-MSI spatial intensities for 36:3 and 38:6 PE across the liver lobule from all male and female mice. Scale bar = 7 mm.

**Supplemental Figure 13. Female LKO Upregulate PPARα and PPARγ Gene Expression.** (**A**) Principal component analysis for bulk RNA-sequencing data from HFD-fed mice (n=6). (**B**) RNA levels of *Pparα*, *Pparδ*, *Pparγ1*, and *Pparγ2* measured by RNA-sequencing (*Pparα*, *Pparδ*) or by qPCR (*Pparγ1*, and *Pparγ2*). (**C**) RNA levels of *Fads2*, *Lpcat3*, and *Tlcd2* measured by qPCR. Significance was determined by two-way ANOVA with Tukey’s multiple comparison *post hoc* analysis (n=6-8). The ANOVA results have been provided. *P<0.05; **P<0.01.

**Supplemental Figure 14. Top Peptides Differentially Altered with CPT1a-deficiency.** A serine-threonine kinome analysis was completed on pooled liver samples from female HFD-control and LKO mice. A total of 6 mice per sex and genotype were pooled and ran in triplicate on the PamGene PamStation. (**A**, **B**) Heatmaps showing the top peptides differently phosphorylated in response to CPT1a deletion in liver lysates from male (**A**) and female (**B**) mice.

**Supplemental Figure 15. Female LKO Mice Also Exhibit Impaired PKA Activity in the Liver.** A serine-threonine kinome analysis was completed on pooled liver samples from female HFD-control and LKO mice. A total of 6 mice per sex and genotype were pooled and ran in triplicate on the PamGene PamStation. (**A**-**C**) PKA peacock (**A**), MEOW (**B**), and waterfall plots (**C**) are presented comparing female control and LKO mice. The waterfall plot shows all known peptides that are phosphorylated by PKA, and the extent by which their phosphorylation, in real-time, is increased or decreased (in red) with *Cpt1a*-deficiency.

