## Supplemental Table 5 for "Loss of Carnitine Palmitoyltransferase 1a Reduces Docosahexaenoic Acid-Containing Phospholipids and Drives Sexually Dimorphic Liver Disease in Mice"

**Supplemental Table 5.** Antibodies used for immunoblotting.

| Protein | Catalog Information | Species & Clonality | Dilution |
|---|---|---|---|
| CPT1a | 128568 | Mouse Monoclonal | 1:1000 |
| CPT1b | 22170-1-AP | Rabbit Polyclonal | 1:1000 |
| Vinculin | NB600-1293 | Mouse Monoclonal | 1:1000 |
| PEMT | Gift from Dr. René Jacobs (rabbit) | | 1:1000 |
| PLIN2 | NB110-40877 | Rabbit Polyclonal | 1:1000 |
| PLIN5 | GP31 | Pig Polyclonal | 1:1000 |
| VDAC | CS-4866S | Rabbit Polyclonal | 1:1000 |
| GAPDH | 5174T | Rabbit Monoclonal | 1:1000 |
| ATGL | 2138 | Rabbit Polyclonal | 1:1000 |
| CGI-58 | NB110-41576 | Rabbit Polyclonal | 1:1000 |
| G0S2 | Gift from Dr. Jun Liu (rabbit) | | 1:1000 |
| HSL | 4107 | Rabbit Polyclonal | 1:1000 |
| MGL | sc-398942 | Mouse Monoclonal | 1:1000 |
| PKA Substrate | 9624 | Mouse Monoclonal | 1:1000 |
