## Supplemental Table 1 for "Loss of Carnitine Palmitoyltransferase 1a Reduces Docosahexaenoic Acid-Containing Phospholipids and Drives Sexually Dimorphic Liver Disease in Mice"

### **Supplemental Table 1.** STK Peptide Substrates.

| Row, Col | ID | Sequence | Ser | Thr | SpotConc | UniprotAccession | Description |
|---|---|---|---|---|---|---|---|
| 1,1 | pTY3H_64_78 | RFIGRRQ(pS)LIEDARK | [71] | [] | 50 | P07101 | Tyrosine 3-monooxygenase (EC 1.14.16.2) (Tyrosine 3-hydroxylase) (TH). |
| 1,2 | ATF2_47_59 | VADQTPTPTRFLK | [] | [51, 53, 55] | 1000 | P15336 | Cyclic AMP-dependent transcription factor ATF-2 (Activatingtranscription factor 2) (cAMP response element-binding |
| 1,3 | CDN1A_139_151 | GRKRRQTSMTDFY | [146] | [145, 148] | 1000 | P38936 | Cyclin-dependent kinase inhibitor 1 (p21) (CDK-interacting protein 1) (Melanoma differentiation-associated protein 6) (MDA-6). |
| 1,4 | FIBA_569_581 | EFPSRGKSSSYSK | [572, 576, 577, 578, 580] | [] | 1000 | P02671 | Fibrinogen alpha chain precursor [Contains: Fibrinopeptide A]. |
| 1,5 | IKKB_173_185_C179A | LDQGSLATSFVGT | [177, 181] | [180, 185] | 1000 | O14920 | Inhibitor of nuclear factor kappa-B kinase subunit beta (I-kappa-B-kinase beta) (IkBKB) (IKK-beta) (IKK-B) (EC=2.7.11.10) (I-kappa-B kinase 2) (IKK2) (Nuclear factor NF-kappa-B inhibitor kinase beta) (NFKBIKB). |
| 1,6 | LIPS_944_956 | GFHPRRSSQGATQ | [950, 951] | [955] | 1000 | Q05469 | Hormone-sensitive lipase (HSL) (EC=3.1.1.79). |
| 1,7 | MYBB_513_525 | DNTPHTPTPFKNA | [] | [515, 518, 520] | 1000 | P10244 | Myb-related protein B (B-Myb). |
| 1,8 | PLEK_106_118 | GQKFARKSTRRSI | [113, 117] | [114] | 1000 | P08567 | Pleckstrin (Platelet p47 protein). |
| 1,9 | RBL2_632_644 | DEICIAGSPLTPR | [639] | [642] | 1000 | Q08999 | Retinoblastoma-like protein 2 (130 kDa retinoblastoma-associatedprotein) (p130) (PRB2) (RBR-2). |
| 1,10 | VIGLN_289_301 | EEKKKKTTTIAVE | [] | [295, 296, 297] | 1000 | Q00341 | Vigilin (High density lipoprotein-binding protein) (HDL-bindingprotein). |
| 1,11 | GRIK2_708_720 | FMSSRRQSVLVKS | [710, 711, 715, 720] | [] | 200 | Q13002 | Glutamate receptor, ionotropic kainate 2 precursor (Glutamate receptor6) (GluR-6) (GluR6) (Excitatory amino acid receptor 4) (EAA4). |
| 1,12 | RADI_559_569 | RDKYKTLRQIR | [] | [564] | 200 | P35241 | Radixin. |
| 2,1 | ACM1_421_433 | CNKAFRDTFRLLL | [] | [428] | 1000 | P11229 | Muscarinic acetylcholine receptor M1. |
| 2,2 | ATM_1972_1984 | KRSLAFEEGSQST | [1974, 1981, 1983] | [1984] | 1000 | Q13315 | Serine-protein kinase ATM (EC=2.7.11.1) (Ataxia telangiectasia mutated) (A-T, mutated). |
| 2,3 | CDN1B_151_163 | IRKRPATDDSSTQ | [160, 161] | [157, 162] | 1000 | P46527 | Cyclin-dependent kinase inhibitor 1B (Cyclin-dependent kinase inhibitor p27) (p27Kip1). |
| 2,4 | FOXO3_25_37 | QSRPRSCTWPLQR | [26, 30] | [32] | 1000 | O43524 | Forkhead box protein O3 (Forkhead in rhabdomyosarcoma-like 1) (AF6q21protein). |
| 2,5 | IKKB_686_698 | QLMSQPSTASNSL | [689, 692, 695, 697] | [693] | 1000 | O14920 | Inhibitor of nuclear factor kappa-B kinase subunit beta (I-kappa-B-kinase beta) (IkBKB) (IKK-beta) (IKK-B) (EC=2.7.11.10) (I-kappa-B kinase 2) (IKK2) (Nuclear factor NF-kappa-B inhibitor kinase beta) (NFKBIKB). |
| 2,6 | LMNA_192_204 | DAENRLQTMKEEL | [] | [199] | 1000 | P02545 | Lamin-A/C (70 kDa lamin) (Renal carcinoma antigen NY-REN-32). |
| 2,7 | MYC_51_63 | KKFELLPTPPLSP | [62] | [58] | 1000 | P01106 | Myc proto-oncogene protein (c-Myc) (Transcription factor p64). |
| 2,8 | PP2AB_297_309 | EPHVTRRTPDYFL | [] | [301, 304] | 1000 | P62714 | Serine/threonine-protein phosphatase 2A catalytic subunit beta isoform(EC 3.1.3.16) (PP2A-beta). |
| 2,9 | RBL2_655_667 | GLGRSITSPTTLY | [659, 662] | [661, 664, 665] | 1000 | Q08999 | Retinoblastoma-like protein 2 (130 kDa retinoblastoma-associatedprotein) (p130) (PRB2) (RBR-2). |
| 2,10 | YAP1_121_133 | QHVRAHSSPASLQ | [127, 128, 131] | [] | 1000 | P46937 | 65 kDa Yes-associated protein (YAP65). |
| 2,11 | KAP2_92_104 | SRFNRRVSVCAET | [92, 99] | [104] | 200 | P13861 | cAMP-dependent protein kinase type II-alpha regulatory subunit. |
| 2,12 | RS6_228_240 | IAKRRRLSSLRAS | [235, 236, 240] | [] | 200 | P62753 | 40S ribosomal protein S6 (Phosphoprotein NP33). |
| 3,1 | ACM1_444_456 | KIPKRPGSVHRTP | [451] | [455] | 1000 | P11229 | Muscarinic acetylcholine receptor M1. |
| 3,2 | BAD_112_124 | RELRRMSDEFVDS | [118, 124] | [] | 1000 | Q92934 | Bcl2 antagonist of cell death (BAD) (Bcl-2-binding component 6) (Bcl-XL/Bcl-2-associated death promoter) (Bcl-2-like 8 protein). |
| 3,3 | CENPA_1_14 | MGPRRRSRKPEAPR | [7] | [] | 1000 | P49450 | Histone H3-like centromeric protein A (Centromere protein A) (CENP-A)(Centromere autoantigen A). |

| | | | | | | | |
|---|---|---|---|---|---|---|---|
| 3,4 | FRAP_2443_2455 | RTRTDSYSAGQSV | [2448, 2450, 2454] | [2444, 2446] | 1000 | P42345 | FKBP12-rapamycin complex-associated protein (FK506-binding protein 12-rapamycin complex-associated protein 1) (Rapamycin target protein) (RAPT1) (Mammalian target of rapamycin) (mTOR). |
| 3,5 | K6PL_766_778 | LEHVTRRTLSMDK | [775] | [770, 773] | 1000 | P17858 | 6-phosphofructokinase, liver type (EC 2.7.1.11) (Phosphofructokinase1) (Phosphohexokinase) (Phosphofructo-1-kinase isozyme B) (PFK-B). |
| 3,6 | LMNB1_16_28 | GGPTTPLSPTRLS | [23, 28] | [19, 20, 25] | 1000 | P20700 | Lamin-B1. |
| 3,7 | NEK2_172_184 | FAKTFVGTPYYMS | [184] | [175, 179] | 1000 | P51955 | Serine/threonine-protein kinase Nek2 (EC 2.7.11.1) (NimA-relatedprotein kinase 2) (NimA-like protein kinase 1) (HSPK 21). |
| 3,8 | PPR1A_28_40 | QIRRRRPTPATLV | [] | [35, 38] | 1000 | Q13522 | Protein phosphatase 1 regulatory subunit 1A (Protein phosphataseinhibitor 1) (IPP-1) (I-1). |
| 3,9 | RBL2_959_971 | DRTSRDSSPVMRS | [962, 965, 966, 971] | [961] | 1000 | Q08999 | Retinoblastoma-like protein 2 (130 kDa retinoblastoma-associatedprotein) (p130) (PRB2) (RBR-2). |
| 3,10 | ADRB2_338_350 | ELLCLRRSSLKAY | [345, 346] | [] | 200 | P07550 | Beta-2 adrenergic receptor (Beta-2 adrenoceptor) (Beta-2adrenoreceptor). |
| 3,11 | KCC2G_278_289 | VASMMHRQETVE | [280] | [287] | 200 | Q13555 | Calcium/calmodulin-dependent protein kinase type II gamma chain(EC 2.7.11.17) (CaM-kinase II gamma chain) (CaM kinase II subunitgamma) (CaMK-II subunit gamma). |
| 3,12 | RYR1_4317_4329 | VRRLRRLTAREAA | [] | [4324] | 200 | P21817 | Ryanodine receptor 1 (Skeletal muscle-type ryanodine receptor) (RyR1)(RYR-1) (Skeletal muscle calcium release channel). |
| 4,1 | ACM4_456_468 | CNATFKKTFRHLL | [] | [459, 463] | 1000 | P08173 | Muscarinic acetylcholine receptor M4. |
| 4,2 | BAD_69_81 | IRSRHSSYPAGTE | [71, 74, 75] | [80] | 1000 | Q92934 | Bcl2 antagonist of cell death (BAD) (Bcl-2-binding component 6) (Bcl-XL/Bcl-2-associated death promoter) (Bcl-2-like 8 protein). |
| 4,3 | COF1_17_29 | DMKVRKSSTPEEV | [23, 24] | [25] | 1000 | P23528 | Cofilin-1 (Cofilin, non-muscle isoform) (18 kDa phosphoprotein) (p18). |
| 4,4 | FRAP_2475_2487 | VPESIHSFIGDGL | [2478, 2481] | [] | 1000 | P42345 | FKBP12-rapamycin complex-associated protein (FK506-binding protein 12-rapamycin complex-associated protein 1) (Rapamycin target protein) (RAPT1) (Mammalian target of rapamycin) (mTOR). |
| 4,5 | KAPCG_192_206 | VKGRTWTLCGTPEYL | [] | [196, 198, 202] | 1000 | P22612 | cAMP-dependent protein kinase catalytic subunit gamma (EC 2.7.11.11)(PKA C-gamma). |
| 4,6 | MARCS_152_164 | KKKKKRFSFKKSF | [159, 163] | [] | 1000 | P29966 | Myristoylated alanine-rich C-kinase substrate (MARCKS) (Protein kinaseC substrate, 80 kDa protein, light chain) (PKCSL) (80K-L protein). |
| 4,7 | NEK3_158_170 | FACTYVGTPYYVP | [] | [161, 165] | 1000 | P51956 | Serine/threonine-protein kinase Nek3 (EC 2.7.11.1) (NimA-relatedprotein kinase 3) (HSPK 36). |
| 4,8 | PRKDC_2618_2630 | TRTQEGSLSARWP | [2624, 2626] | [2618, 2620] | 1000 | P78527 | DNA-dependent protein kinase catalytic subunit (DNA-PK catalytic subunit) (DNA-PKcs) (EC=2.7.11.1) (DNPK1) (p460). |
| 4,9 | REL_260_272 | KMQLRRPSDQEVS | [267, 272] | [] | 1000 | Q04864 | C-Rel proto-oncogene protein (C-Rel protein). |
| 4,10 | ART_025_CXGLRRWSLGGLRRWSL | GLRRWSLGGLRRWSL | NA | NA | 200 | NA | NA |
| 4,11 | KCNA1_438_450 | DSDLSRRSSSTMS | [439, 442, 445, 446, 447, 450] | [448] | 200 | Q09470 | Potassium voltage-gated channel subfamily A member 1 (Voltage-gatedpotassium channel subunit Kv1.1) (HUKI) (HBK1). |
| 4,12 | SCN7A_898_910 | KNGCRRGSSLGQI | [905, 906] | [] | 200 | Q01118 | Sodium channel protein type 7 subunit alpha (Sodium channel proteintype VII subunit alpha) (Putative voltage-gated sodium channel subunitalpha Nax) (Sodium channel protein cardiac and skeletal muscle |
| 5,1 | ACM5_494_506 | CYALCNRTFRKTF | [] | [501, 505] | 1000 | P08912 | Muscarinic acetylcholine receptor M5. |
| 5,2 | BAD_93_105 | FRGRSRSAPPNLW | [97, 99] | [] | 1000 | Q92934 | Bcl2 antagonist of cell death (BAD) (Bcl-2-binding component 6) (Bcl-XL/Bcl-2-associated death promoter) (Bcl-2-like 8 protein). |
| 5,3 | CSF1R_701_713 | NIHLEKKYVRRDS | [713] | [] | 1000 | P07333 | Macrophage colony-stimulating factor 1 receptor precursor(EC 2.7.10.1) (CSF-1-R) (Fms proto-oncogene) (c-fms) (CD115 antigen). |
| 5,4 | GPR6_349_361 | QSKVPFRSRSPSE | [350, 356, 358, 360] | [] | 1000 | P46095 | Sphingosine 1-phosphate receptor GPR6 (G-protein coupled receptor 6). |
| 5,5 | KCNA2_442_454 | PDLKKSRSASTIS | [447, 449, 451, 454] | [452] | 1000 | P16389 | Potassium voltage-gated channel subfamily A member 2 (Voltage-gatedpotassium channel subunit Kv1.2) (HBK5) (NGK1) (HUKIV). |

| | | | | | | | |
|---|---|---|---|---|---|---|---|
| 5,6 | MARCS_160_172 | FKKSFKLSGFSFK | [163, 167, 170] | [] | 1000 | P29966 | Myristoylated alanine-rich C-kinase substrate (MARCKS) (Protein kinaseC substrate, 80 kDa protein, light chain) (PKCSL) (80K-L protein). |
| 5,7 | NMDZ1_890_902 | SFKRRRSSKDTST | [890, 896, 897, 901] | [900, 902] | 1000 | Q05586 | Glutamate [NMDA] receptor subunit zeta-1 precursor (N-methyl-D-aspartate receptor subunit NR1). |
| 5,8 | PTK6_436_448 | ALRERLSSFTSYE | [442, 443, 446] | [445] | 1000 | Q13882 | Tyrosine-protein kinase 6 (EC 2.7.10.2) (Breast tumor kinase)(Tyrosine-protein kinase BRK). |
| 5,9 | SRC_413_425 | LIEDNEYTARQGA | [] | [420] | 1000 | P12931 | Proto-oncogene tyrosine-protein kinase Src (EC 2.7.10.2) (p60-Src) (c-Src) (pp60c-src). |
| 5,10 | CAC1C_1974_1986 | ASLGRRASFHLEC | [1975, 1981] | [] | 200 | Q13936 | Voltage-dependent L-type calcium channel subunit alpha-1C (Voltage-gated calcium channel subunit alpha Cav1.2) (Calcium channel, L type,alpha-1 polypeptide, isoform 1, cardiac muscle). |
| 5,11 | KPB1_1011_1023 | QVEFRRLSISAES | [1018, 1020, 1023] | [] | 200 | P46020 | Phosphorylase b kinase regulatory subunit alpha, skeletal muscleisoform (Phosphorylase kinase alpha M subunit). |
| 5,12 | SRC8_CHICK_423_435 | KTPSSPVYQDAVS | [426, 427, 435] | [424] | 200 | Q01406 | Src substrate protein p85 (p80) (Cortactin). |
| 6,1 | ACM5_498_510 | CNRTFRKTFKMLL | [] | [501, 505] | 1000 | P08912 | Muscarinic acetylcholine receptor M5. |
| 6,2 | BCKD_45_57 | ERSKTVTSFYNQS | [47, 52, 57] | [49, 51] | 1000 | O14874 | [3-methyl-2-oxobutanoate dehydrogenase [lipoamide]] kinase,mitochondrial precursor (EC 2.7.11.4) (Branched-chain alpha-ketoaciddehydrogenase kinase) (BCKDHKIN) (BCKD-kinase). |
| 6,3 | CSK21_355_367 | ISSVPTPSPLGPL | [356, 357, 362] | [360] | 1000 | P68400 | Casein kinase II subunit alpha (EC 2.7.11.1) (CK II). |
| 6,4 | GPSM2_394_406 | PKLGRRHSMENME | [401] | [] | 1000 | P81274 | G-protein-signaling modulator 2 (Mosaic protein LGN). |
| 6,5 | KCNA3_461_473 | EELRKARSNSTLS | [468, 470, 473] | [471] | 1000 | P22001 | Potassium voltage-gated channel subfamily A member 3 (Voltage-gatedpotassium channel subunit Kv1.3) (HPCN3) (HGK5) (HuKIII) (HLK3). |
| 6,6 | MBP_222_234 | HFFKNIVTPRTPP | [] | [229, 232] | 1000 | P02686 | Myelin basic protein (MBP) (Myelin A1 protein) (Myelin membraneencephalitogenic protein). |
| 6,7 | NOS3_1171_1183 | SRIRTQSFSLQER | [1171, 1177, 1179] | [1175] | 1000 | P29474 | Nitric oxide synthase, endothelial (EC=1.14.13.39) (Endothelial NOS) (eNOS) (EC-NOS) (NOS type III) (NOSIII) (Constitutive NOS) (cNOS). |
| 6,8 | RAB1A_187_199 | KSNVKIQSTPVKQ | [188, 194] | [195] | 1000 | P62820 | Ras-related protein Rab-1A (YPT1-related protein). |
| 6,9 | STK6_283_295 | SSRRTTLCGTLDY | [283, 284] | [287, 288, 292] | 1000 | O14965 | Serine/threonine-protein kinase 6 (EC 2.7.11.1) (Aurora kinase A)(Aurora-A) (Serine/threonine kinase 15) (Aurora/IPL1-related kinase 1)(Aurora-related kinase 1) (hARK1) (Breast tumor-amplified kinase). |
| 6,10 | CFTR_730_742 | EPLERRLSLVPDS | [737, 742] | [] | 200 | P13569 | Cystic fibrosis transmembrane conductance regulator (CFTR) (cAMP-dependent chloride channel) (ATP-binding cassette transporter sub- |
| 6,11 | MYPC3_268_280 | LSAFRRTSLAGGG | [269, 275] | [274] | 200 | Q14896 | Myosin-binding protein C, cardiac-type (Cardiac MyBP-C) (C-protein,cardiac muscle isoform). |
| 6,12 | VASP_271_283 | LARRRKATQVGEK | [] | [278] | 200 | P50552 | Vasodilator-stimulated phosphoprotein (VASP). |
| 7,1 | ADDB_696_708 | GSPSKSPSKKKKK | [697, 699, 701, 703] | [] | 1000 | P35612 | Beta-adducin (Erythrocyte adducin subunit beta) |
| 7,2 | BRCA1_1451_1463 | EKAVLTSQKSSEY | [1457, 1460, 1461] | [1456] | 1000 | P38398 | Breast cancer type 1 susceptibility protein (RING finger protein 53). |
| 7,3 | DCX_49_61 | HFDERDKTSRNMR | [57] | [56] | 1000 | O43602 | Neuronal migration protein doublecortin (Lissencephalin-X) (Lis-X)(Doublin). |
| 7,4 | GSUB_61_73 | KKPRRKDTPALHI | [] | [68] | 1000 | O96001 | G-substrate. |
| 7,5 | KCNB1_489_501 | KWTKRTLSETSSS | [496, 499, 500, 501] | [491, 494, 498] | 1000 | Q14721 | Potassium voltage-gated channel subfamily B member 1 (Voltage-gatedpotassium channel subunit Kv2.1) (h-DRK1). |
| 7,6 | MK10_214_226 | AGTSFMMTPYVVT | [217] | [216, 221, 226] | 1000 | P53779 | Mitogen-activated protein kinase 10 (EC 2.7.11.24) (Stress-activated protein kinase JNK3) (c-Jun N-terminal kinase 3) (MAP kinase p49 3F12). |
| 7,7 | NR4A1_344_356 | GRRGRLPSKPKQP | [351] | [] | 1000 | P22736 | Nuclear receptor subfamily 4 group A member 1 (Orphan nuclear receptorHMR) (Early response protein NAK1) (TR3 orphan receptor) (ST- |
| 7,8 | RAF1_253_265 | QRQRSTSTPNVHM | [257, 259] | [258, 260] | 1000 | P04049 | RAF proto-oncogene serine/threonine-protein kinase (EC 2.7.11.1) (Raf-1) (C-RAF) (cRaf). |

| | | | | | | | |
|---|---|---|---|---|---|---|---|
| 7,9 | STMN2_90_102 | AAGERRKSQEAQV | [97] | [] | 1000 | Q93045 | Stathmin-2 (Protein SCG10) (Superior cervical ganglion-10 protein). |
| 7,10 | CGHB_109_121 | QCALCRRSTTDCG | [116] | [117, 118] | 200 | P01233 | Choriogonadotropin subunit beta precursor (CG-beta) (Chorionicgonadotrophin chain beta). |
| 7,11 | NCF1_296_308 | RGAPPRRSSIRNA | [303, 304] | [] | 200 | P14598 | Neutrophil cytosol factor 1 (NCF-1) (Neutrophil NADPH oxidase factor1) (47 kDa neutrophil oxidase factor) (p47-phox) (NCF-47K) (47 kDaautosomal chronic granulomatous disease protein) (Nox organizer 2)(Nox-organizing protein 2) (SH3 and PX domain-containing protein |
| 7,12 | VTNC_390_402 | NQNSRRPSRATWL | [393, 397] | [400] | 200 | P04004 | Vitronectin precursor (Serum-spreading factor) (S-protein) (V75)[Contains: Vitronectin V65 subunit; Vitronectin V10 |
| 8,1 | ADDB_706_718 | KKKFRTPSFLKKS | [713, 718] | [711] | 1000 | P35612 | Beta-adducin (Erythrocyte adducin subunit beta) |
| 8,2 | C1R_201_213 | ASGYISSLEYPRS | [202, 206, 207, 213] | [] | 1000 | P00736 | Complement C1r subcomponent (EC=3.4.21.41) |
| 8,3 | ELK1_356_368 | LLPTHTLTPVLLT | [] | [359, 361, 363, 368] | 1000 | P19419 | ETS domain-containing protein Elk-1. |
| 8,4 | GYS2_1_13 | MLRGRSLSVTSLG | [6, 8, 11] | [10] | 1000 | P54840 | Glycogen [starch] synthase, liver (EC 2.4.1.11). |
| 8,5 | KIF11_919_931 | LDIPTGTTPQRKS | [931] | [923, 925, 926] | 1000 | P52732 | Kinesin-like protein KIF11 (Kinesin-related motor protein Eg5)(Kinesin-like spindle protein HKSP) (Thyroid receptor-interactingprotein 5) (TRIP-5) (Kinesin-like protein 1). |
| 8,6 | MP2K1_281_293 | GDAAETPPRPRTP | [] | [286, 292] | 1000 | Q02750 | Dual specificity mitogen-activated protein kinase kinase 1(EC 2.7.12.2) (MAP kinase kinase 1) (MAPKK 1) (ERK activator kinase 1)(MAPK/ERK |
| 8,7 | NTRK3_824_836 | LHALGKATPIYLD | [] | [831] | 1000 | Q16288 | NT-3 growth factor receptor precursor (EC 2.7.10.1) (Neurotrophictyrosine kinase receptor type 3) (TrkC tyrosine kinase) |
| 8,8 | RAP1B_172_184 | PGKARKKSSCQLL | [179, 180] | [] | 1000 | P61224 | Ras-related protein Rap-1b precursor (GTP-binding protein smg p21B). |
| 8,9 | TAU_524_536 | GSRSRTPSLPTPP | [525, 527, 531] | [529, 534] | 1000 | P10636 | Microtubule-associated protein tau (Neurofibrillary tangle protein)(Paired helical filament-tau) (PHF-tau). |
| 8,10 | CREB1_126_138 | EILSRRPSYRKIL | [129, 133] | [] | 200 | P16220 | cAMP response element-binding protein (CREB). |
| 8,11 | NCF1_321_333 | QDAYRRNSVRFLQ | [328] | [] | 200 | P14598 | Neutrophil cytosol factor 1 (NCF-1) (Neutrophil NADPH oxidase factor1) (47 kDa neutrophil oxidase factor) (p47-phox) (NCF-47K) (47 kDaautosomal chronic granulomatous disease protein) (Nox organizer 2)(Nox-organizing protein 2) (SH3 and PX domain-containing protein |
| 8,12 | CFTR_761_773 | LQARRRQSVLNLM | [768] | [] | 200 | P13569 | Cystic fibrosis transmembrane conductance regulator (CFTR) (cAMP-dependent chloride channel) (ATP-binding cassette transporter sub- |
| 9,1 | AKT1_301_313 | KDGATMKTFCGTP | [] | [305, 308, 312] | 1000 | P31749 | RAC-alpha serine/threonine-protein kinase (EC 2.7.11.1) (RAC-PK-alpha)(Protein kinase B) (PKB) (C-AKT). |
| 9,2 | CA2D1_494_506 | LEDIKRLTPRFTL | [] | [501, 505] | 1000 | P54289 | Voltage-dependent calcium channel subunit alpha-2/delta-1 precursor(Voltage-gated calcium channel subunit alpha-2/delta-1) [Contains:Voltage-dependent calcium channel subunit alpha-2-1; |
| 9,3 | ELK1_410_422 | ISVDGLSTPVVLS | [411, 416, 422] | [417] | 1000 | P19419 | ETS domain-containing protein Elk-1. |
| 9,4 | H2B1B_27_40 | GKKRKRSRKESYSI | [33, 37, 39] | [] | 1000 | P33778 | Histone H2B type 1-B (H2B.f) (H2B/f) (H2B.1). |
| 9,5 | KIF2C_105_118_S106G | EGLRSRSTRMSTVS | [109, 111, 115, 118] | [112, 116] | 1000 | Q99661 | Kinesin-like protein KIF2C (Mitotic centromere-associated kinesin)(MCAK) (Kinesin-like protein 6). |
| 9,6 | MP2K1_287_299 | PPRPRTPGRPLSS | [298, 299] | [292] | 1000 | Q02750 | Dual specificity mitogen-activated protein kinase kinase 1(EC 2.7.12.2) (MAP kinase kinase 1) (MAPKK 1) (ERK activator kinase 1)(MAPK/ERK |
| 9,7 | P53_12_24 | PPLSQETFSDLWK | [15, 20] | [18] | 1000 | P04637 | Cellular tumor antigen p53 (Tumor suppressor p53) (Phosphoprotein p53)(Antigen NY-CO-13). |
| 9,8 | RB_242_254 | AVIPINGSPRTPR | [249] | [252] | 1000 | P06400 | Retinoblastoma-associated protein (PP110) (P105-RB) (RB). |
| 9,9 | TLE2_246_258 | EPPSPATTPCGKV | [249] | [252, 253] | 1000 | Q04725 | Transducin-like enhancer protein 2 (ESG2). |

| | | | | | | | |
|---|---|---|---|---|---|---|---|
| 9,10 | DESP_2842_2854 | RSGSRRGSFDATG | [2843, 2845, 2849] | [2853] | 200 | P15924 | Desmoplakin (DP) (250/210 kDa paraneoplastic pemphigus antigen). |
| 9,11 | NFKB1_330_342 | FVQLRRKSDLETS | [337, 342] | [341] | 200 | P19838 | Nuclear factor NF-kappa-B p105 subunit (DNA-binding factor KBF1) (EBP-1) [Contains: Nuclear factor NF-kappa-B p50 subunit]. |
| 9,12 | F263_454_466 | NPLMRRNSVTPLA | [461] | [463] | 200 | Q16875 | 6-phosphofructo-2-kinase/fructose-2,6-biphosphatase 3 (6PF-2-K/Fru-2,6-P2ASE brain/placenta-type isozyme) (iPFK-2) (Renal carcinomaantigen NY-REN-56) [Includes: 6-phosphofructo-2-kinase (EC 2.7.1.105);Fructose-2,6-bisphosphatase (EC 3.1.3.46)]. |
| 10,1 | ANDR_785_797 | VRMRHLSQEFGWL | [791] | [] | 1000 | P10275 | Androgen receptor (Dihydrotestosterone receptor) (Nuclear receptor subfamily 3 group C member 4). |
| 10,2 | CD27_212_224 | HQRRKYRSNKGES | [219, 224] | [] | 1000 | P26842 | CD27 antigen precursor (CD27L receptor) (T-cell activation antigenCD27) (T14) (Tumor necrosis factor receptor superfamily |
| 10,3 | ERBB2_679_691 | QQKIRKYTMRRLL | [] | [686] | 1000 | P04626 | Receptor tyrosine-protein kinase erbB-2 precursor (EC 2.7.10.1)(p185erbB2) (C-erbB-2) (NEU proto-oncogene) (Tyrosine kinase-type cellsurface receptor HER2) (MLN 19) (CD340 antigen). |
| 10,4 | H32_3_18 | RTKQTARKSTGGKAPR | [11] | [4, 7, 12] | 1000 | Q71DI3 | Histone H3.2 (H3/m) (H3/o). |
| 10,5 | KPCB_19_31_A25S | RFARKGSLRQKNV | [25] | [] | 1000 | P05771 | Protein kinase C beta type (EC 2.7.11.13) (PKC-beta) (PKC-B). |
| 10,6 | MPH6_140_152 | EDENGDITPIKAK | [] | [147] | 1000 | Q99547 | M-phase phosphoprotein 6. |
| 10,7 | P53_308_323 | LPNNTSSSPQPKKKPL | [313, 314, 315] | [312] | 1000 | P04637 | Cellular tumor antigen p53 (Tumor suppressor p53) (Phosphoprotein p53)(Antigen NY-CO-13). |
| 10,8 | RB_350_362 | SFETQRTPRKSNL | [350, 360] | [353, 356] | 1000 | P06400 | Retinoblastoma-associated protein (PP110) (P105-RB) (RB). |
| 10,9 | TOP2A_1463_1475 | RRKRKPSTSDDSD | [1469, 1471, 1474] | [1470] | 1000 | P11388 | DNA topoisomerase 2-alpha (EC=5.99.1.3) (DNA topoisomerase II, alpha isozyme). |
| 10,10 | E1A_ADE05_212_224 | AILRRPTSPVSRE | [219, 222] | [218] | 200 | P03255 | Early E1A 32 kDa protein. |
| 10,11 | PLM_76_88 | EEGTFRSSIRRLS | [82, 83, 88] | [79] | 200 | O00168 | Phospholemman precursor (FXYD domain-containing ion transportregulator 1). |
| 10,12 | KAP3_107_119 | NRFTRRASVCAEA | [114] | [110] | 200 | P31323 | cAMP-dependent protein kinase type II-beta regulatory subunit. |
| 11,1 | ANXA1_209_221 | AGERRKGTDVNVF | [] | [216] | 1000 | P04083 | Annexin A1 (Annexin-1) (Annexin I) (Lipocortin I) (Calpactin II)(Chromobindin-9) (p35) (Phospholipase A2 inhibitory protein). |
| 11,2 | CDC2_154_169 | GIPIRVYTHEVVTLWY | [] | [161, 166] | 1000 | P06493 | Cell division control protein 2 homolog (EC 2.7.11.22) (EC 2.7.11.23) (p34 protein kinase) (Cyclin-dependent kinase 1) (CDK1). |
| 11,3 | ERF_519_531 | GEAGGPLTPRRVS | [531] | [526] | 1000 | P50548 | ETS domain-containing transcription factor ERF (Ets2 repressorfactor). |
| 11,4 | IF4E_203_215 | TATKSGSTTKNRF | [207, 209] | [203, 205, 210, 211] | 1000 | P06730 | Eukaryotic translation initiation factor 4E (eIF-4E) (eIF4E) (mRNA cap-binding protein) (eIF-4F 25 kDa subunit). |
| 11,5 | KPCB_626_639 | AENFDRFFTRHPPV | [] | [634] | 1000 | P05771-2 | Protein kinase C beta type (EC 2.7.11.13) (PKC-beta) (PKC-B). |
| 11,6 | MPIP1_172_184 | FTQRQNSAPARML | [178] | [173] | 1000 | P30304 | M-phase inducer phosphatase 1 (EC=3.1.3.48) (Dual specificity phosphatase Cdc25A). |
| 11,7 | PDE5A_95_107 | GTPTRKISASEFD | [102, 104] | [96, 98] | 1000 | O76074 | cGMP-specific 3',5'-cyclic phosphodiesterase (EC 3.1.4.35) (CGB-PDE)(cGMP-binding cGMP-specific phosphodiesterase). |
| 11,8 | RB_774_786 | TRPPTLSPIPHIP | [780] | [774, 778] | 1000 | P06400 | Retinoblastoma-associated protein (PP110) (P105-RB) (RB). |
| 11,9 | VASP_150_162 | EHIERRVSNAGGP | [157] | [] | 1000 | P50552 | Vasodilator-stimulated phosphoprotein (VASP). |
| 11,10 | EPB42_241_253 | LLNKRRGSVPILR | [248] | [] | 200 | P16452 | Erythrocyte membrane protein band 4.2 (Erythrocyte protein 4.2)(P4.2). |
| 11,11 | PTN12_32_44 | FMRLRRLSTKYRT | [39] | [40, 44] | 200 | Q05209 | Tyrosine-protein phosphatase non-receptor type 12 (EC 3.1.3.48)(Protein-tyrosine phosphatase G1) (PTPG1) (PTP-PEST). |
| 11,12 | KCNA6_504_516 | ANRERRPSYLPTP | [511] | [515] | 200 | P17658 | Potassium voltage-gated channel subfamily A member 6 (Voltage-gatedpotassium channel subunit Kv1.6) (HBK2). |

| | | | | | | | |
|---|---|---|---|---|---|---|---|
| 12,1 | pVASP_150_164 | EHIERRV(pS)NAGG PPA | [157] | [] | 50 | P50552 | Vasodilator-stimulated phosphoprotein (VASP). |
| 12,2 | CDK7_163_175 | GSPNRAYTHQVVT | [164] | [170, 175] | 1000 | P50613 | Cell division protein kinase 7 (EC 2.7.11.22) (EC 2.7.11.23) (CDK-activating kinase) (CAK) (TFIIH basal transcription factor complexkinase subunit) (39 kDa protein kinase) (P39 Mo15) (STK1) (CAK1). |
| 12,3 | ESR1_160_172 | GGRERLASTNDKG | [167] | [168] | 1000 | P03372 | Estrogen receptor (ER) (Estradiol receptor) (ER-alpha) (Nuclearreceptor subfamily 3 group A member 1). |
| 12,4 | IKBA_26_38 | LDDRHDSGLDSMK | [32, 36] | [] | 1000 | P25963 | NF-kappa-B inhibitor alpha (I-kappa-B-alpha) (IkappaBalpha) (IkB-alpha) (Major histocompatibility complex enhancer-binding protein MAD3). |
| 12,5 | KS6A1_374_386 | QLFRGFSFVATGL | [380] | [384] | 1000 | Q15418 | Ribosomal protein S6 kinase alpha-1 (S6K-alpha 1) (EC=2.7.11.1) (90 kDa ribosomal protein S6 kinase 1) (p90-RSK 1) (pp90RSK1) (p90S6K) (Ribosomal S6 kinase 1) (RSK-1) (MAP kinase-activated protein kinase |
| 12,6 | MPIP3_208_220 | RSGLYRSPSMPEN | [209, 214, 216] | [] | 1000 | P30307 | M-phase inducer phosphatase 3 (EC 3.1.3.48) (Dual specificityphosphatase Cdc25C). |
| 12,7 | PDPK1_27_39 | SMVRTQTESSTPP | [27, 35, 36] | [31, 33, 37] | 1000 | O15530 | 3-phosphoinositide-dependent protein kinase 1 (EC 2.7.11.1) (hPDK1). |
| 12,8 | RB_803_815 | NIYISPLKSPYKI | [807, 811] | [] | 1000 | P06400 | Retinoblastoma-associated protein (PP110) (P105-RB) (RB). |
| 12,9 | VASP_232_244 | GAKLRKVSKQEEA | [239] | [] | 1000 | P50552 | Vasodilator-stimulated phosphoprotein (VASP). |
| 12,10 | GBRB2_427_439 | SRLRRRASQLKIT | [427, 434] | [439] | 200 | P47870 | Gamma-aminobutyric acid receptor subunit beta-2 precursor (GABA(A)receptor subunit beta-2). |
| 12,11 | PYGL_8_20 | QEKRRQISIRGIV | [15] | [] | 200 | P06737 | Glycogen phosphorylase, liver form (EC 2.4.1.1). |
| 12,12 | TY3H_65_77 | FIGRRQSLIEDAR | [71] | [] | 200 | P07101 | Tyrosine 3-monooxygenase (EC 1.14.16.2) (Tyrosine 3-hydroxylase) (TH). |
