## Supplemental Table 2 for "Loss of Carnitine Palmitoyltransferase 1a Reduces Docosahexaenoic Acid-Containing Phospholipids and Drives Sexually Dimorphic Liver Disease in Mice"

**Supplemental Table 2.** STK Scores and Rankings Male LKO versus Control.

| Chip | Kinase Uniprot ID | Kinase Name | UKA Mean Final Score | KRSA Z-Score |
|---|---|---|---|---|
| STK | P51817 | PRKX | 3.111809995 | 2.529633673 |
| STK | Q13976 | PKG1 | 3.158651268 | 2.445281341 |
| STK | P17612 | PKA[alpha] | 3.443324027 | 2.356425466 |
| STK | Q86V86 | Pim3 | 1.308285831 | 2.223978899 |
| STK | Q16539 | MAPK14 | 0.802383313 | -2.156149343 |
| STK | Q13237 | PKG2 | 3.512189444 | 2.135018211 |
| STK | Q16644 | MAPKAPK3 | 1.204716045 | 1.925710638 |
| STK | P11802 | CDK4 | 0.556943338 | -1.847471793 |
| STK | P49137 | MAPKAPK2 | 1.256764996 | 1.772399097 |
| STK | Q9Y6S9 | RSKL2 | 1.342948469 | 1.723372384 |
| STK | P45983 | JNK1 | 0.306887231 | -1.717927171 |
| STK | P45984 | JNK2 | 0.363192669 | -1.64946 |
| STK | P53779 | JNK3 | 0.354832933 | -1.637249591 |
| STK | P11309 | Pim1 | 1.080952117 | 1.601519942 |
| STK | Q9UBS0 | p70S6K[beta] | 3.192738193 | 1.575058299 |
| STK | Q00535 | CDK5 | 0.729472959 | -1.525831101 |
| STK | P24941 | CDK2 | 0.309493576 | -1.388225822 |
| STK | P49840 | GSK3[alpha] | 0.159335565 | -1.276965997 |
| STK | Q96L96 | AlphaK1 | 0.920202592 | 1.24626505 |
| STK | O75676 | MSK2 | 0.113929424 | 1.244412482 |
| STK | Q00526 | CDK3 | 0.961761287 | -1.239957819 |
| STK | Q92772 | CDKL2 | 1.574792073 | 1.232795042 |
| STK | P42345 | mTOR/FRAP | 0.394560245 | -1.184449408 |
| STK | Q00534 | CDK6 | 0.539734886 | -1.163128249 |
| STK | P06493 | CDC2/CDK1 | 1.005935185 | -1.144363283 |
| STK | P68400 | CK2[alpha]1 | 1.353155423 | -1.142067594 |
| STK | Q9HBY8 | SGK2 | 1.492588211 | 1.127073508 |
| STK | P41279 | COT | 2.193056892 | -1.111654847 |
| STK | Q15759 | p38[beta] | 0.473979192 | -1.034682702 |
| STK | O75582 | MSK1 | 0.279723604 | 1.003195413 |
| STK | Q13464 | ROCK1 | 0.835902842 | 0.956265635 |
| STK | P53778 | p38[gamma] | 0.265015714 | -0.954828781 |
| STK | O14965 | AurA/Aur2 | 0.015671136 | -0.946553347 |
| STK | P28482 | ERK2 | 0.321431267 | -0.940762732 |
| STK | Q00532 | CDKL1 | 0.953171277 | -0.936833094 |
| STK | Q9P1W9 | Pim2 | 1.654101631 | 0.918937472 |
| STK | Q96Q40 | PFTAIRE2 | 1.275658344 | -0.871056317 |
| STK | O60285 | NuaK1 | 1.347129647 | -0.870020344 |
| STK | P50613 | CDK7 | 0.297692008 | 0.858574411 |
| STK | P48730 | CK1[delta] | 3.308379687 | 0.844231234 |
| STK | Q05513 | PKC[zeta] | 0.217103998 | -0.8273527 |
| STK | Q15139 | PKD1 | 2.288134594 | 0.818032911 |

| | | | | |
|---|---|---|---|---|
| STK | Q9BWU1 | CDK11 | 0.395290293 | -0.810860528 |
| STK | Q05655 | PKC[delta] | 1.648729418 | 0.802004893 |
| STK | Q9UIK4 | DAPK2 | 1.452185287 | -0.801664316 |
| STK | P17252 | PKC[alpha] | 1.440882864 | 0.784405507 |
| STK | O14757 | CHK1 | 0.812534559 | 0.783460579 |
| STK | P27361 | ERK1 | 0.355786452 | -0.767228048 |
| STK | Q9UQM7 | CaMK2[alpha] | 2.06321783 | 0.727911225 |
| STK | Q13131 | AMPK[alpha]1 | 1.559729428 | 0.712480458 |
| STK | Q14164 | IKK[epsilon] | 0.517511478 | 0.707545496 |
| STK | O76039 | CDKL5 | 0.924661328 | 0.693455456 |
| STK | O94921 | PFTAIRE1 | 1.576398879 | -0.686932972 |
| STK | P31751 | Akt2/PKB[beta] | 2.23528387 | 0.684157997 |
| STK | Q13627 | DYRK1A | 0.878424816 | 0.682118833 |
| STK | P23443 | p70S6K | 1.259665334 | 0.657977832 |
| STK | O43293 | DAPK3 | 1.154695654 | -0.651974963 |
| STK | Q13535 | ATR | 1.532426525 | -0.645059086 |
| STK | Q02156 | PKC[epsilon] | 1.151991027 | 0.641736374 |
| STK | Q13164 | ERK5 | 0.857342792 | -0.541072671 |
| STK | P15056 | BRAF | 1.687822644 | -0.520283125 |
| STK | P49674 | CK1[epsilon] | 2.331926044 | 0.514907145 |
| STK | Q15418 | RSK3 | 1.312410551 | -0.50241818 |
| STK | P04049 | RAF1 | 0.773789417 | -0.502183724 |
| STK | Q8NI60 | ADCK3 | 0.298337511 | -0.497094973 |
| STK | P49841 | GSK3[beta] | 0.159837603 | -0.485460908 |
| STK | O96017 | CHK2 | 1.523917342 | 0.466538477 |
| STK | P05771 | PKC[beta] | 0.252050689 | -0.463127953 |
| STK | O14920 | IKK[beta] | 1.669783497 | 0.428529467 |
| STK | P51812 | RSK2 | 1.006270391 | 0.417001964 |
| STK | Q16512 | PKN1/PRK1 | 0.521640261 | 0.406786354 |
| STK | O75116 | ROCK2 | 1.020509203 | 0.387205506 |
| STK | P05129 | PKC[gamma] | 0.708389234 | -0.385589925 |
| STK | P24723 | PKC[eta] | 0.702617592 | 0.373518155 |
| STK | O15264 | p38[delta] | 0.173671854 | -0.354647515 |
| STK | Q15131 | CDK10 | 1.261302099 | 0.346928941 |
| STK | P31749 | Akt1/PKB[alpha] | 2.740098368 | 0.292863781 |
| STK | Q00537 | PCTAIRE2 | 0.883977021 | 0.287881876 |
| STK | Q8IWB6 | SgK307 | 1.178288967 | -0.280337815 |
| STK | Q8TD08 | ERK7 | 0.814559844 | 0.262998604 |
| STK | O43930 | PRKY | 1.710450348 | 0.23740726 |
| STK | P48729 | CK1[alpha] | 2.05883531 | -0.22299891 |
| STK | O15111 | IKK[alpha] | 3.117044722 | 0.20052033 |
| STK | P50750 | CDK9 | 0.164390034 | 0.132980272 |
| STK | Q16566 | CaMK4 | 2.593234543 | 0.108770163 |

| | | | | |
|---|---|---|---|---|
| STK | Q9UHD2 | TBK1 | 1.042579242 | -0.08094341 |
| STK | Q04759 | PKC[theta] | 0.918340295 | -0.07287373 |
| STK | P16066 | ANP[alpha] | 1.365475179 | 0.052870348 |
| STK | Q15349 | RSK1/p90RSK | 1.62507002 | -0.049420269 |
| STK | Q13153 | PAK1 | 0.080711017 | -0.032328312 |
| STK | Q96GD4 | AurB/Aur1 | 0.116466043 | -0.023903083 |
| STK | P41743 | PKC[iota] | 1.121345481 | 0.022648981 |
| STK | P53355 | DAPK1 | 0.598446019 | 0.001821918 |
