## Supplemental Table 3 for "Loss of Carnitine Palmitoyltransferase 1a Reduces Docosahexaenoic Acid-Containing Phospholipids and Drives Sexually Dimorphic Liver Disease in Mice"

**Supplemental Table 3.** STK Scores and Rankings Female LKO versus Control.

| Chip | Kinase Uniprot ID | Kinase Name | UKA Mean Final Score | KRSA Z-Score |
|---|---|---|---|---|
| STK | Q9UBS0 | p70S6K[beta] | 1.017341984 | 2.933655215 |
| STK | P51817 | PRKX | 1.942685529 | 2.666579126 |
| STK | P27361 | ERK1 | 0.399523427 | -2.567561766 |
| STK | Q9UHD2 | TBK1 | 0.363808154 | -2.51196494 |
| STK | P28482 | ERK2 | 0.42285169 | -2.457333075 |
| STK | Q16539 | MAPK14 | 0.634491863 | -2.431932472 |
| STK | P17612 | PKA[alpha] | 3.194708056 | 2.217966944 |
| STK | P45983 | JNK1 | 0.717784731 | -2.03310928 |
| STK | P31749 | Akt1/PKB[alpha] | 2.241460632 | 1.933808204 |
| STK | P53779 | JNK3 | 0.71979217 | -1.888316325 |
| STK | P06493 | CDC2/CDK1 | 0.592942691 | -1.885753009 |
| STK | O15264 | p38[delta] | 0.574215613 | -1.883733192 |
| STK | Q13976 | PKG1 | 1.761794596 | 1.861818926 |
| STK | Q9Y6S9 | RSKL2 | 1.110588827 | 1.837524663 |
| STK | P24941 | CDK2 | 0.628888811 | -1.773208064 |
| STK | P41743 | PKC[iota] | 0.562723253 | -1.762719071 |
| STK | Q16644 | MAPKAPK3 | 0.969017277 | 1.686049842 |
| STK | Q86V86 | Pim3 | 1.052172601 | 1.667473768 |
| STK | Q96L96 | AlphaK1 | 8.70E-04 | -1.626209413 |
| STK | P45984 | JNK2 | 0.799735289 | -1.620462457 |
| STK | Q14164 | IKK[epsilon] | 1.173806428 | -1.604896154 |
| STK | Q9UIK4 | DAPK2 | 0.979700427 | -1.481603616 |
| STK | Q15759 | p38[beta] | 0.472812624 | -1.466695976 |
| STK | O96017 | CHK2 | 1.354050318 | 1.449775095 |
| STK | Q8NI60 | ADCK3 | 0.86262824 | 1.41117678 |
| STK | P11309 | Pim1 | 1.141398696 | 1.392256907 |
| STK | P49840 | GSK3[alpha] | 1.83955768 | -1.347157303 |
| STK | P53778 | p38[gamma] | 0.476992767 | -1.291731828 |
| STK | Q13464 | ROCK1 | 0.456646091 | -1.285532889 |
| STK | O94921 | PFTAIRE1 | 1.614393972 | 1.262696415 |
| STK | O75676 | MSK2 | 1.754244576 | 1.254151907 |
| STK | P15056 | BRAF | 1.516327136 | -1.249582085 |
| STK | Q9HBY8 | SGK2 | 1.67746092 | 1.208933976 |
| STK | P51812 | RSK2 | 2.088153928 | 1.145524947 |
| STK | P48729 | CK1[alpha] | 0.72670781 | -1.129774788 |
| STK | P16066 | ANP[alpha] | 0.30772192 | -1.129290353 |
| STK | P49137 | MAPKAPK2 | 1.353509862 | 1.125169053 |
| STK | P49674 | CK1[epsilon] | 0.34452163 | -1.1196131 |
| STK | P31751 | Akt2/PKB[beta] | 2.298553684 | 1.098341817 |
| STK | P41279 | COT | 1.712197337 | -1.093630141 |
| STK | P49841 | GSK3[beta] | 1.845537345 | -1.088881206 |
| STK | Q00534 | CDK6 | 0.837452808 | -1.08457661 |
| STK | O75582 | MSK1 | 1.750172301 | 1.018943256 |
| STK | O14965 | AurA/Aur2 | 0.6957088 | 0.965605408 |
| STK | Q00532 | CDKL1 | 0.580687662 | -0.924763198 |
| STK | P48730 | CK1[delta] | 0.469754592 | -0.922241172 |
| STK | Q8IWB6 | SgK307 | 0.207710433 | -0.909748426 |
| STK | Q16566 | CaMK4 | 1.027563715 | -0.908173196 |

| | | | | |
|---|---|---|---|---|
| STK | O75116 | ROCK2 | 0.966224207 | -0.900332679 |
| STK | Q00535 | CDK5 | 0.852275386 | -0.892377812 |
| STK | P50613 | CDK7 | 0.885639837 | -0.883741579 |
| STK | O60285 | NuaK1 | 1.155036027 | -0.883713334 |
| STK | Q13237 | PKG2 | 1.651737962 | 0.8763904 |
| STK | Q13153 | PAK1 | 0.484710401 | -0.861035142 |
| STK | Q04759 | PKC[theta] | 2.022212117 | 0.860735607 |
| STK | Q13164 | ERK5 | 0.429144742 | -0.84791894 |
| STK | P23443 | p70S6K | 1.066225223 | 0.812421518 |
| STK | O43930 | PRKY | 0.633005657 | 0.759369219 |
| STK | Q9BWU1 | CDK11 | 0.947924679 | -0.727907435 |
| STK | Q00526 | CDK3 | 1.323591928 | -0.723564057 |
| STK | Q9P1W9 | Pim2 | 1.039275033 | 0.715581707 |
| STK | Q9UQM7 | CaMK2[alpha] | 2.028239866 | 0.705704408 |
| STK | Q13627 | DYRK1A | 0.784072915 | 0.705088328 |
| STK | Q8TD08 | ERK7 | 0.263284866 | -0.696495476 |
| STK | O43293 | DAPK3 | 0.58743602 | -0.651338556 |
| STK | Q00537 | PCTAIRE2 | 0.408775068 | -0.60623194 |
| STK | O15111 | IKK[alpha] | 1.229498358 | -0.511868562 |
| STK | P17252 | PKC[alpha] | 1.953322049 | 0.493057703 |
| STK | P42345 | mTOR/FRAP | 0.395380641 | -0.462426092 |
| STK | P05771 | PKC[beta] | 0.679332593 | -0.459115533 |
| STK | O14920 | IKK[beta] | 1.935033939 | 0.458253793 |
| STK | P24723 | PKC[eta] | 1.097582503 | 0.452689499 |
| STK | P04049 | RAF1 | 0.811003937 | -0.445676118 |
| STK | Q15131 | CDK10 | 0.869295638 | 0.421894836 |
| STK | Q05655 | PKC[delta] | 1.539266638 | 0.416866647 |
| STK | O14757 | CHK1 | 0.501981083 | -0.400881993 |
| STK | Q92772 | CDKL2 | 0.867130045 | 0.286952491 |
| STK | P68400 | CK2[alpha]1 | 1.658146824 | -0.284493007 |
| STK | Q16512 | PKN1/PRK1 | 2.586247777 | -0.241828628 |
| STK | Q15418 | RSK3 | 1.650810144 | 0.2382695 |
| STK | Q96Q40 | PFTAIRE2 | 0.837052384 | -0.219568702 |
| STK | O76039 | CDKL5 | 0.401753149 | 0.203647213 |
| STK | P50750 | CDK9 | 1.380166415 | 0.181682903 |
| STK | Q05513 | PKC[zeta] | 0.502175686 | -0.178486987 |
| STK | Q02156 | PKC[epsilon] | 1.269600342 | -0.175910746 |
| STK | Q13131 | AMPK[alpha]1 | 2.064292842 | 0.175882774 |
| STK | P05129 | PKC[gamma] | 1.122933401 | 0.150794753 |
| STK | P11802 | CDK4 | 0.82155846 | -0.045681402 |
| STK | Q15139 | PKD1 | 1.55465892 | 0.029114726 |
| STK | Q15349 | RSK1/p90RSK | 1.965969489 | 0.016764465 |
| STK | Q96GD4 | AurB/Aur1 | 0.721626555 | 0.011984258 |
| STK | P53355 | DAPK1 | 1.683487177 | 0.011740595 |
| STK | Q13535 | ATR | 0.446142034 | -0.004727717 |
