## Supplemental Table 4 for "Loss of Carnitine Palmitoyltransferase 1a Reduces Docosahexaenoic Acid-Containing Phospholipids and Drives Sexually Dimorphic Liver Disease in Mice"

**Supplemental Table 4.** Mouse primers used for real-time PCR.

| **Gene** | **Sequence (5’ – 3’)** | **Forward/Reverse** |
|---|---|---|
| *Cpt1a* | AGTGGCCTCACAGACTCCAG | Forward |
| | GCCCATGTTGTACAGCTTCC | Reverse |
| *Cpt1b* | GCTGCTTGCACATTTGTGTT | Forward |
| | TGAGTGACTGGTGGGAAGAA | Reverse |
| *Pparγ1* | AACAAGACTACCCTTTACTGAAATTACCA | Forward |
| | CACAGAGCTGATTCCGAAGTTG | Reverse |
| *Pparγ2* | CCAGAGCATGGTGCCTTCGCT | Forward |
| | CAGCAACCATTGGGTCAG | Reverse |
| *18S* | GTAACCCGTTGAACCCCATT | Forward |
| | CCATCCAATCGGTAGTAGCG | Reverse |
| *Hprt* | CACGCAACCAGGAAGTAGAA | Forward |
| | AGAGCGAGAACGAACAGATTA | Reverse |
