## Supplemental Figures for "Loss of Carnitine Palmitoyltransferase 1a Reduces Docosahexaenoic Acid-Containing Phospholipids and Drives Sexually Dimorphic Liver Disease in Mice"

Supplemental Figure 1

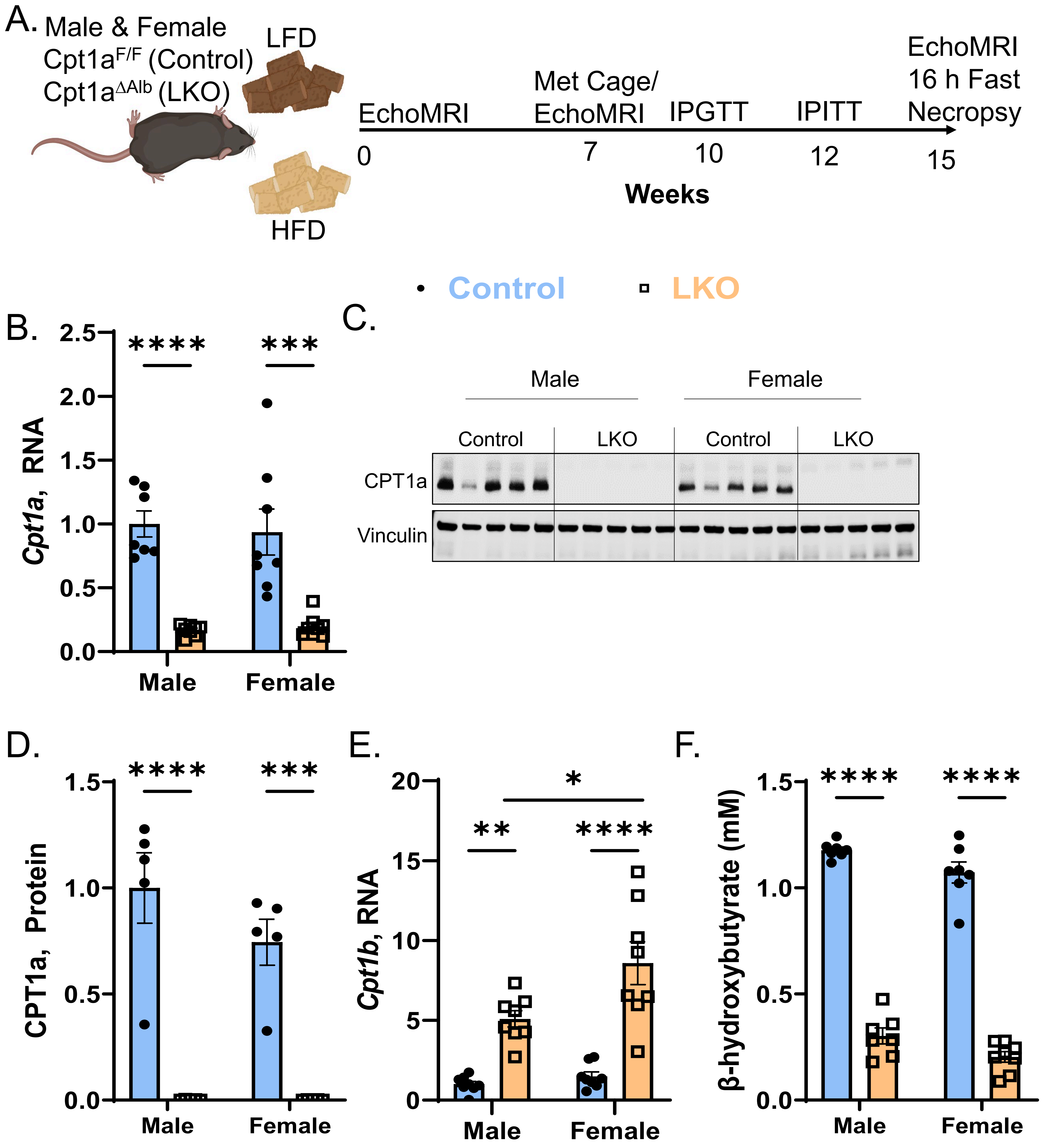

Supplemental Figure 2

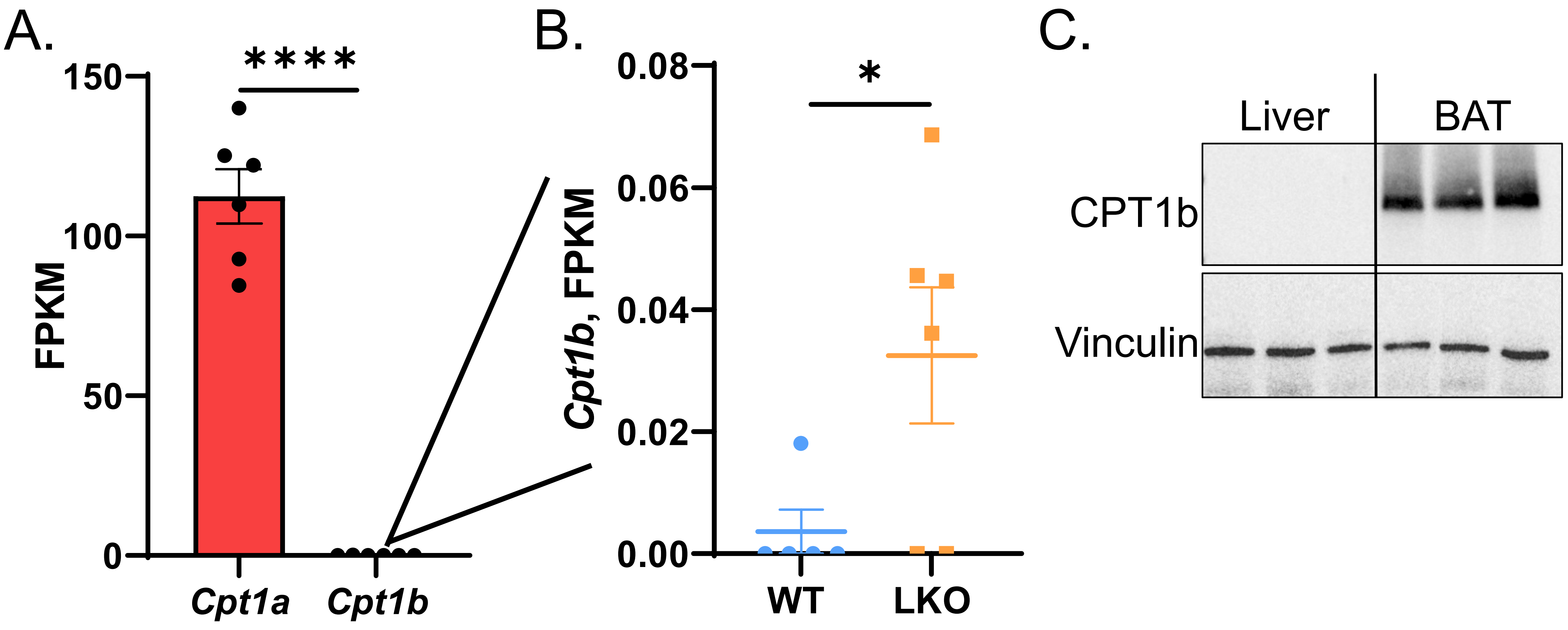

#### Supplemental Figure 3

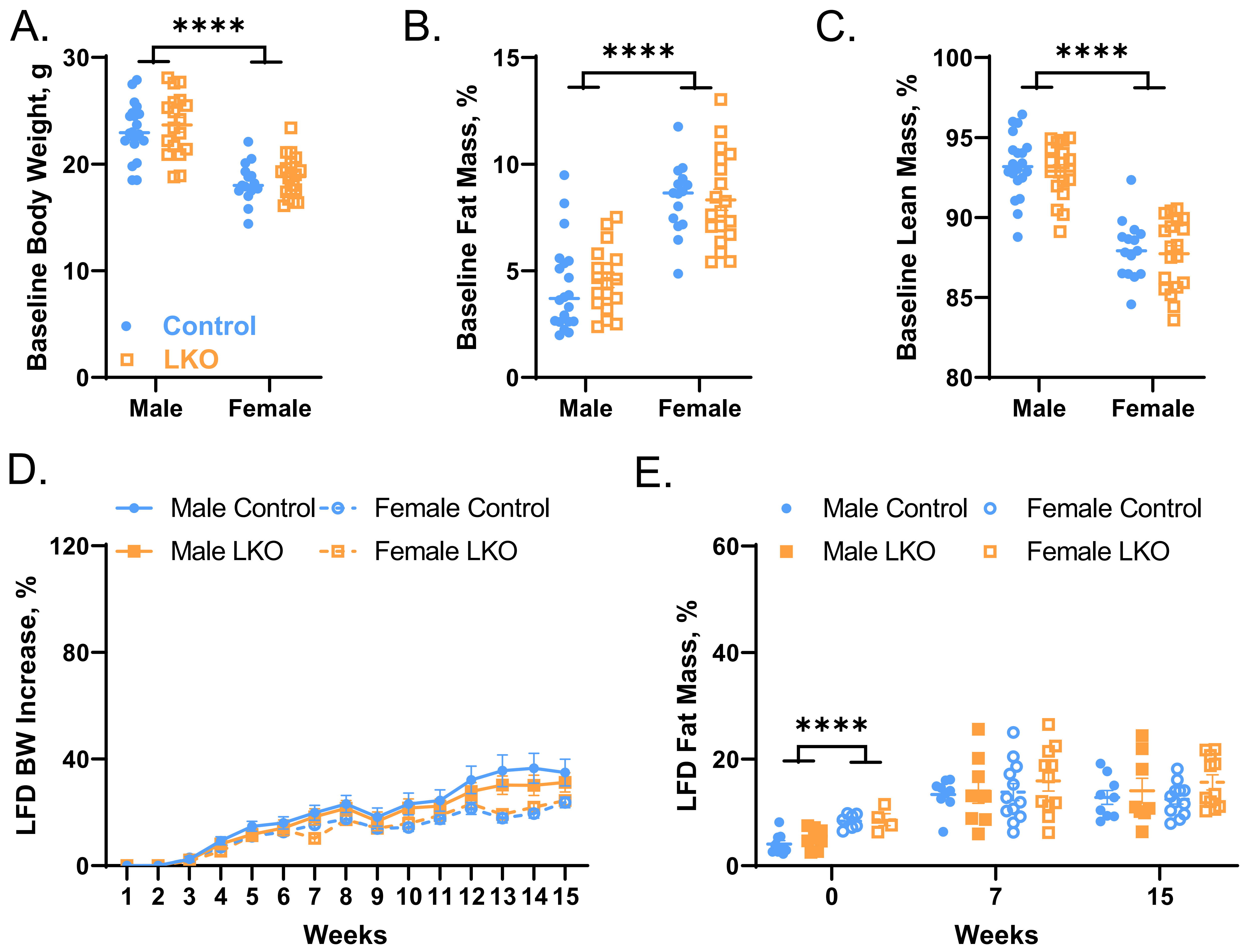

#### Supplemental Figure 4

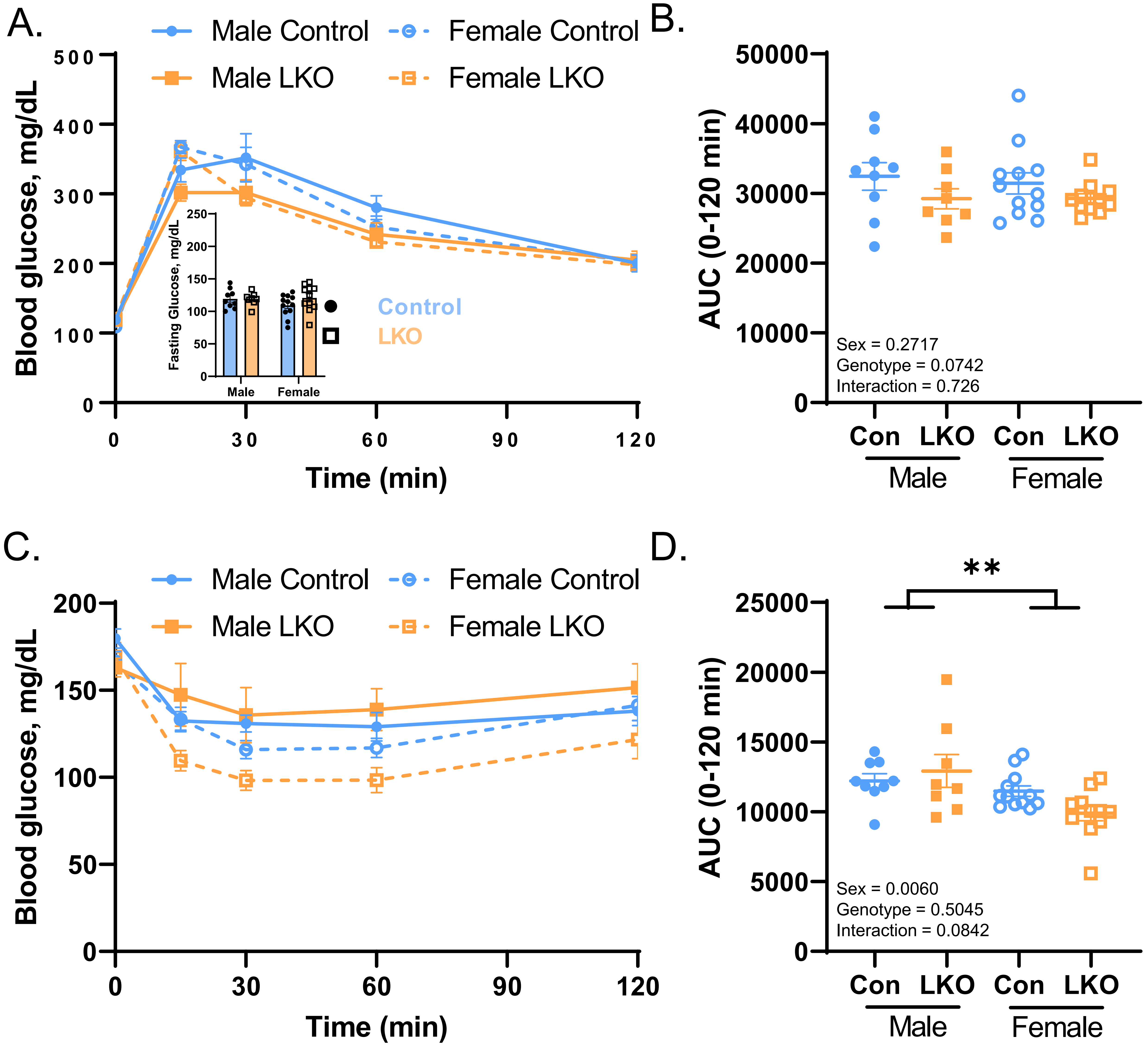

Supplemental Figure 5

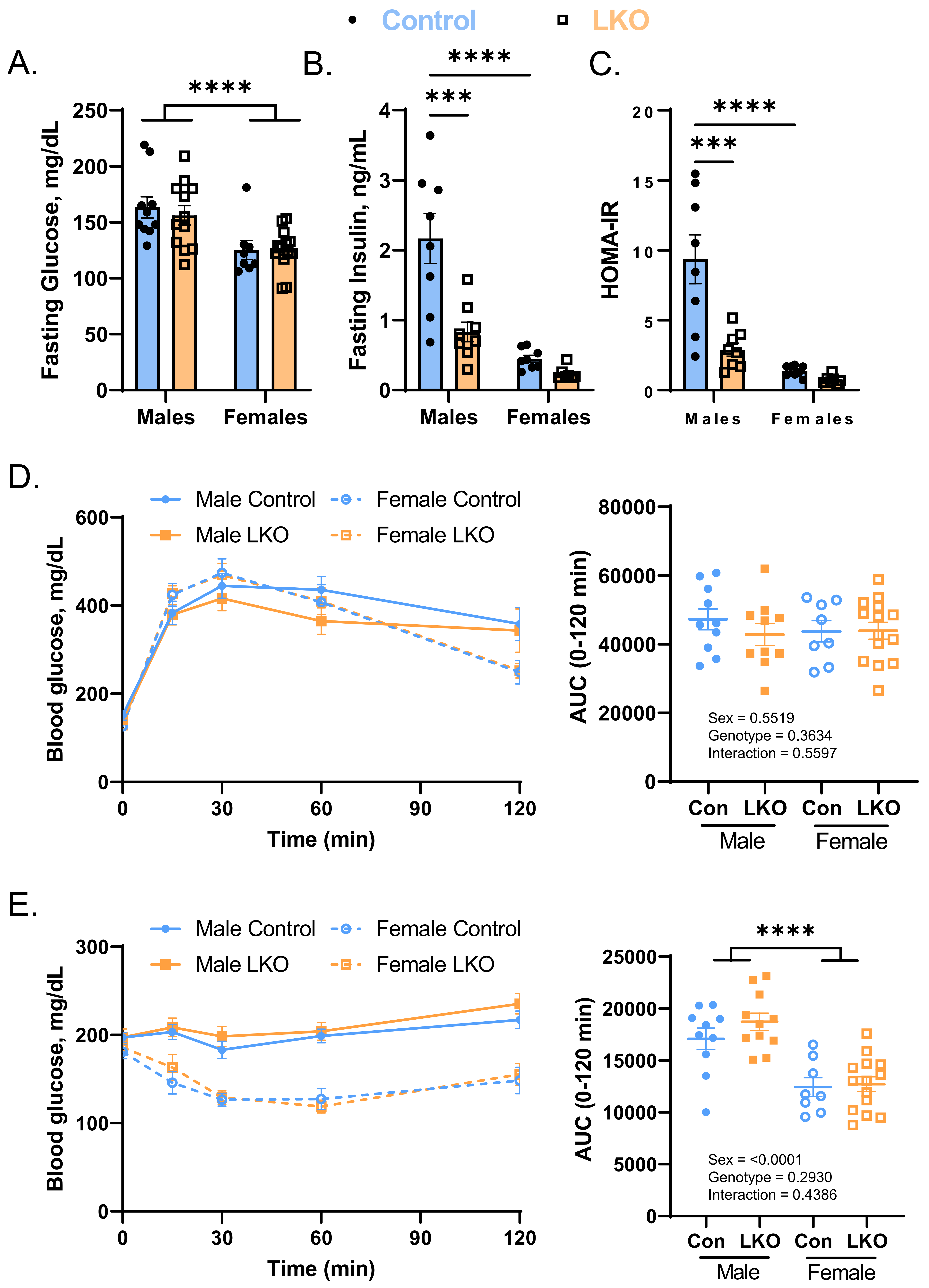

### Supplemental Figure 6

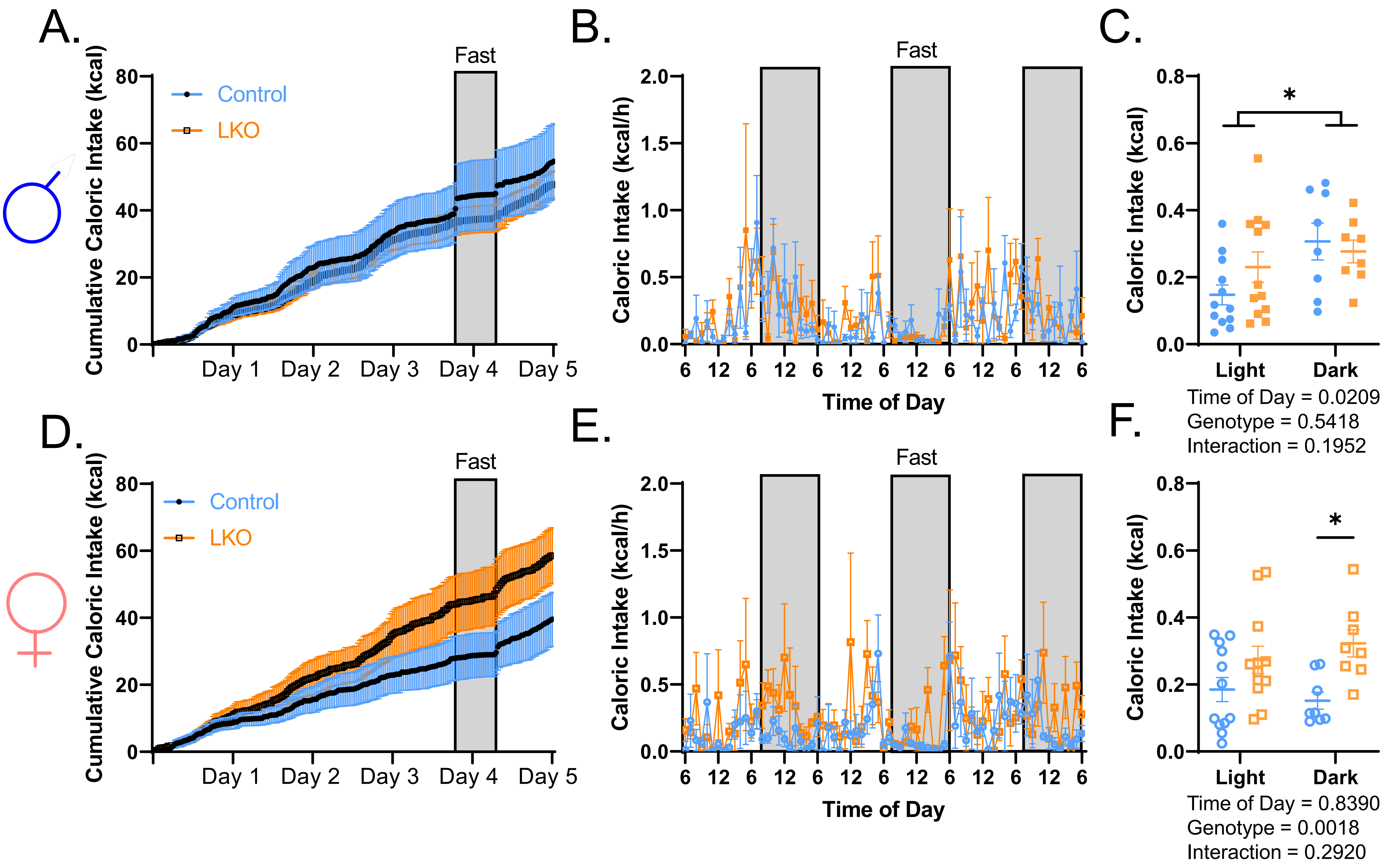

### Supplemental Figure 7

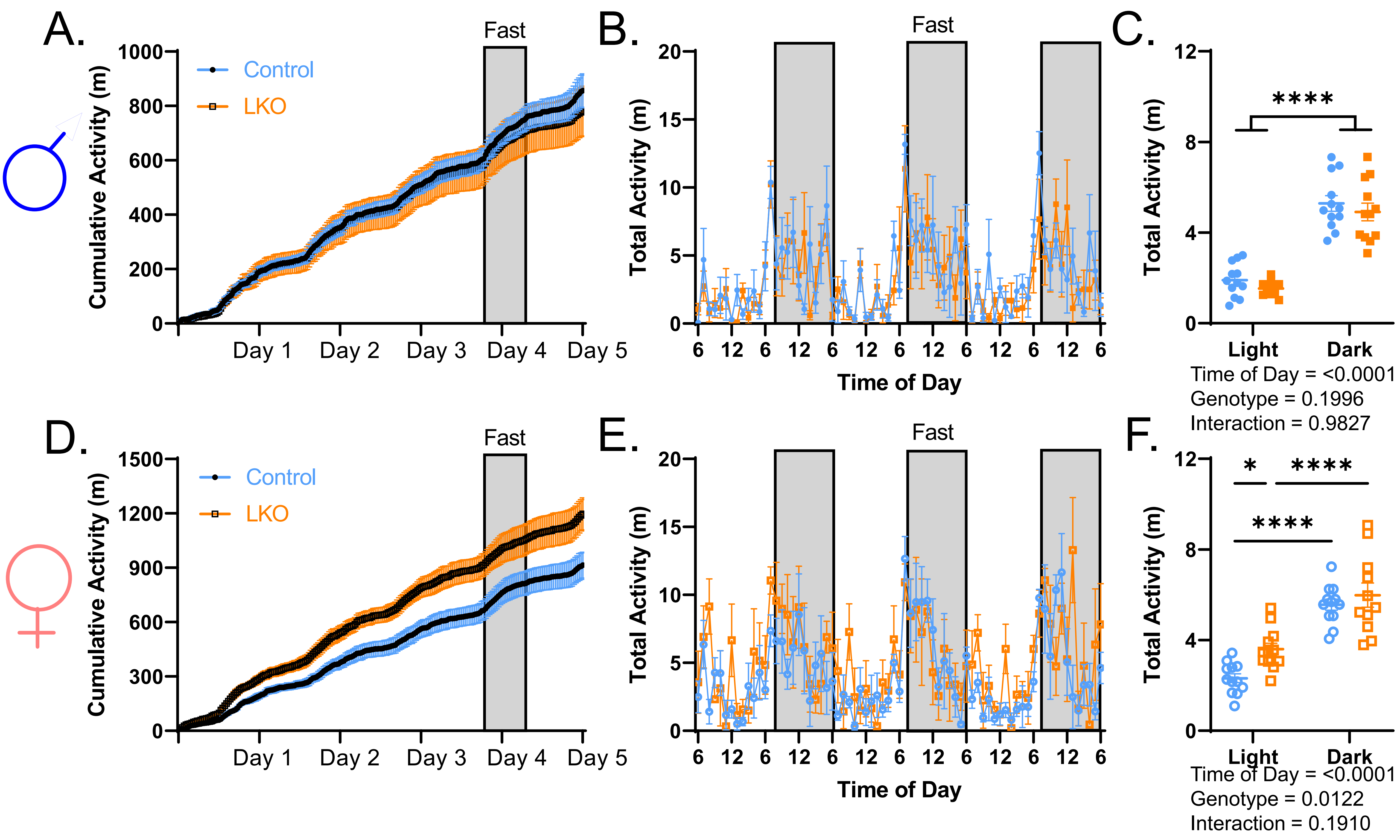

Supplemental Figure 8

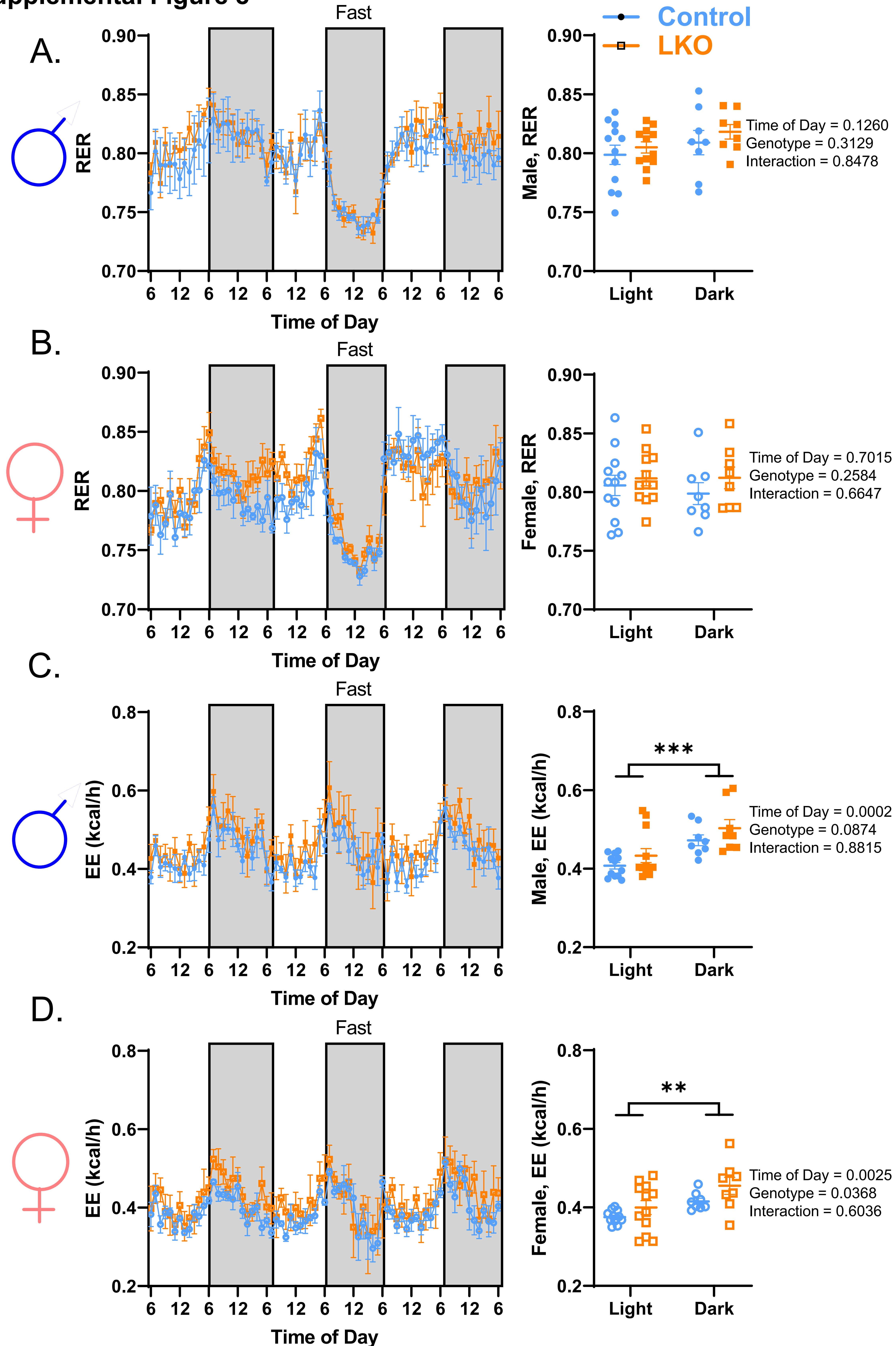

**Supplemental Figure 9**

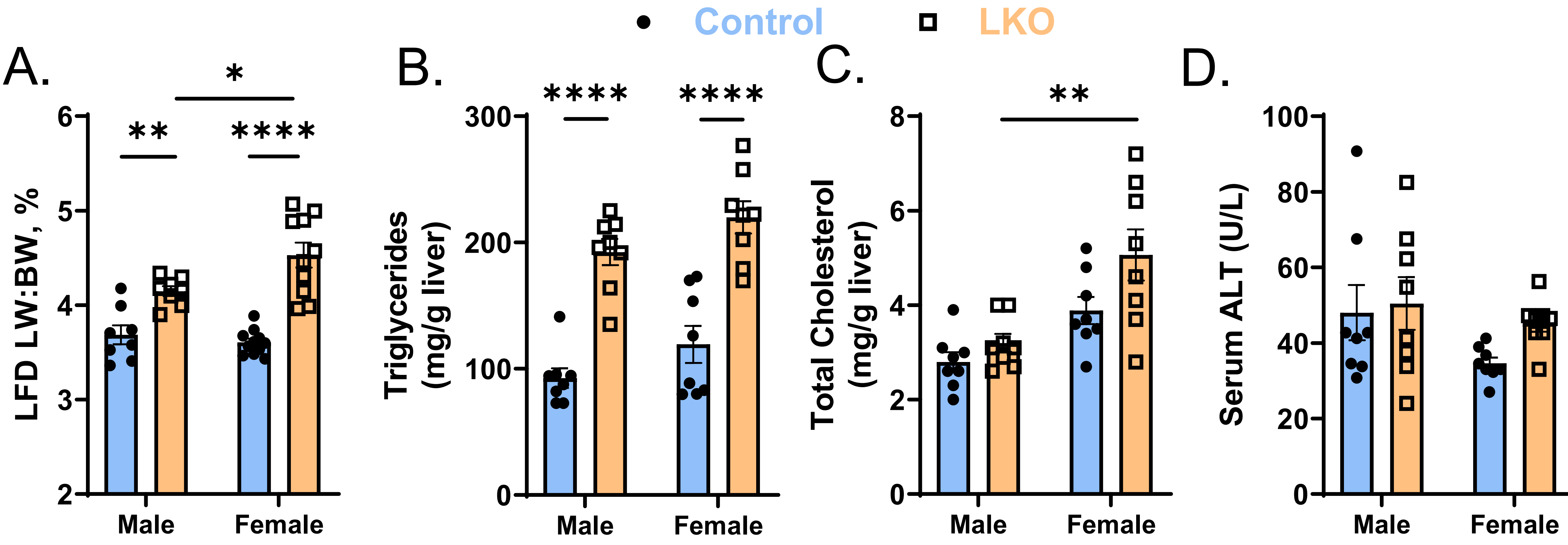

#### Supplemental Figure 10

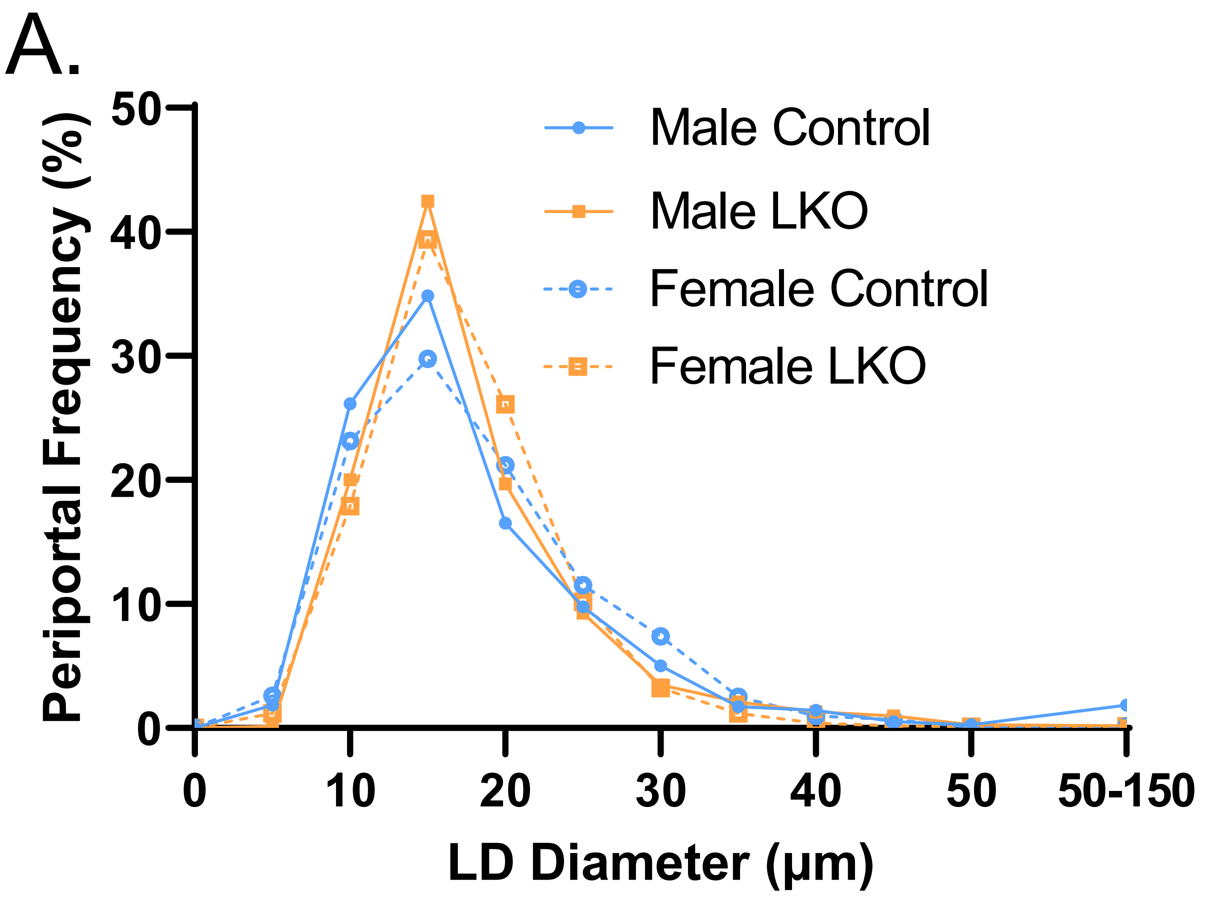


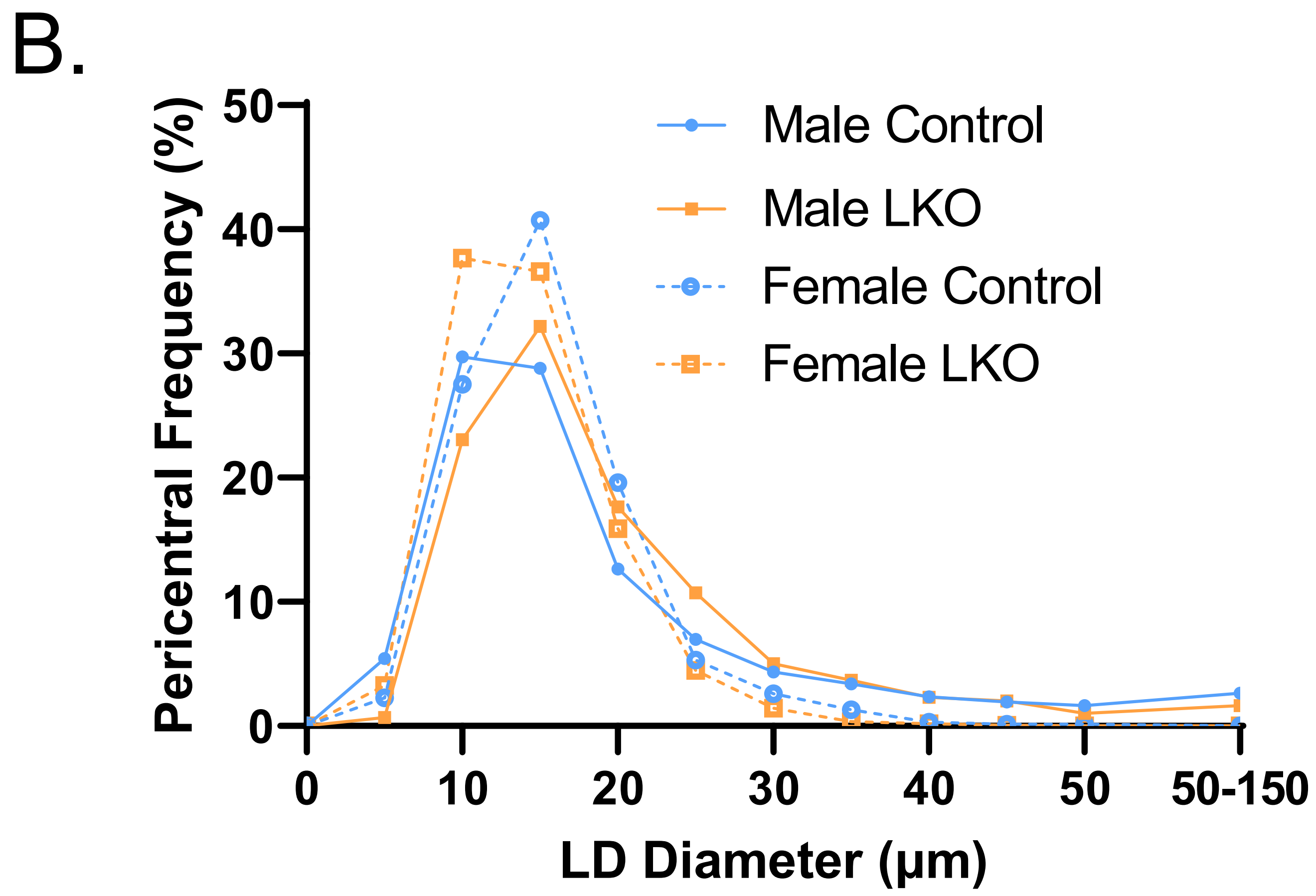

**Supplemental Figure 11**

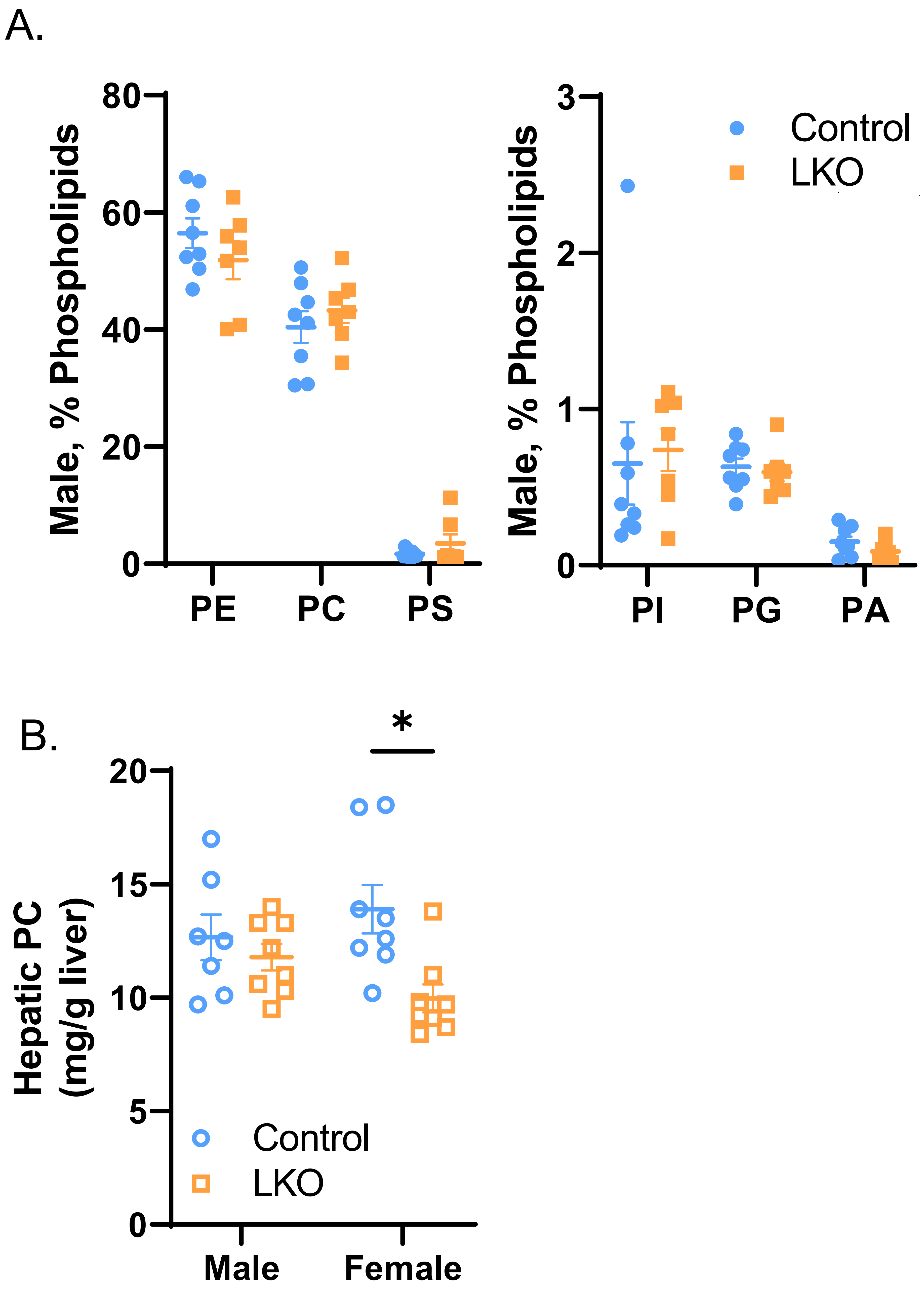

Supplemental Figure 12

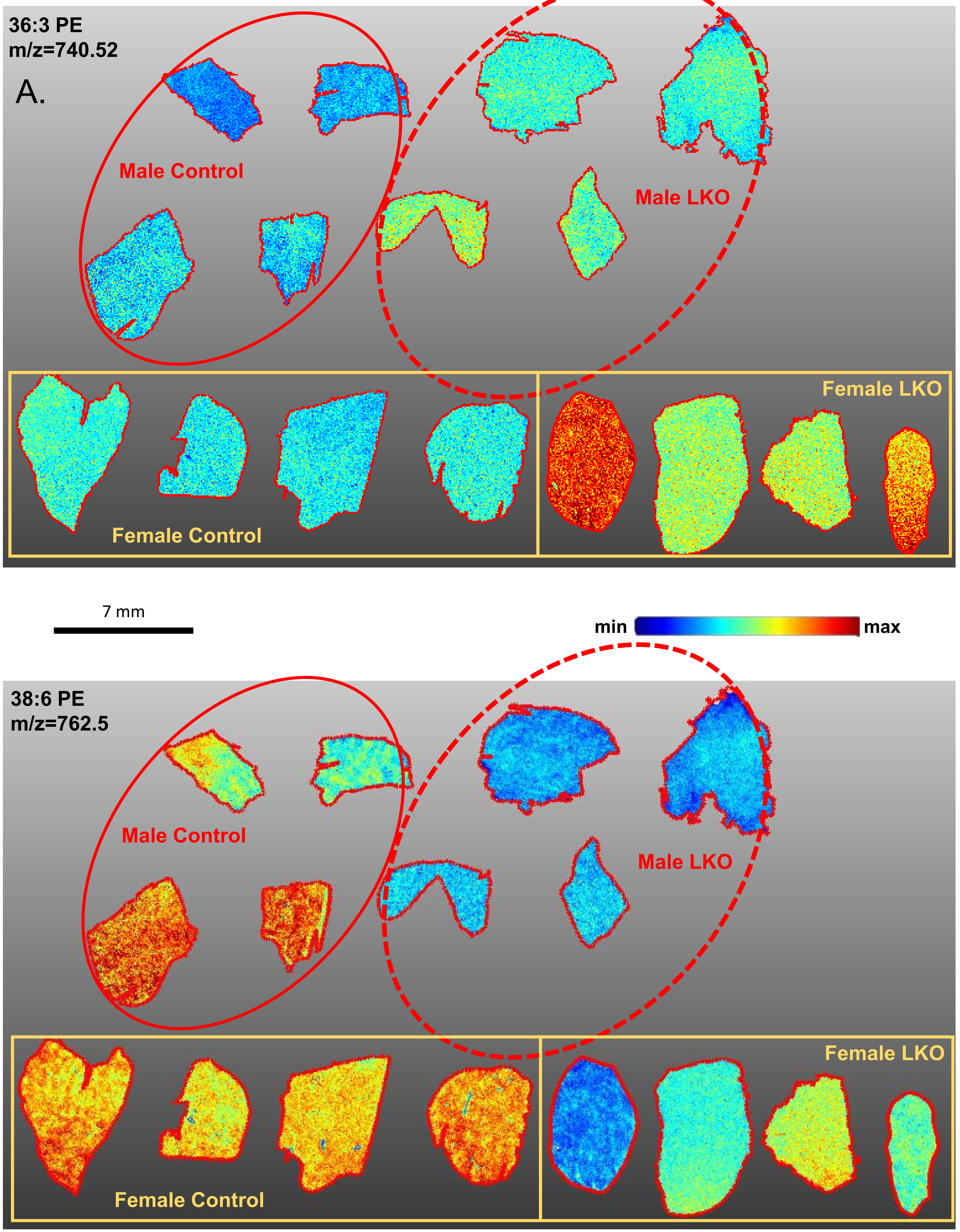

**Supplemental Figure 13**

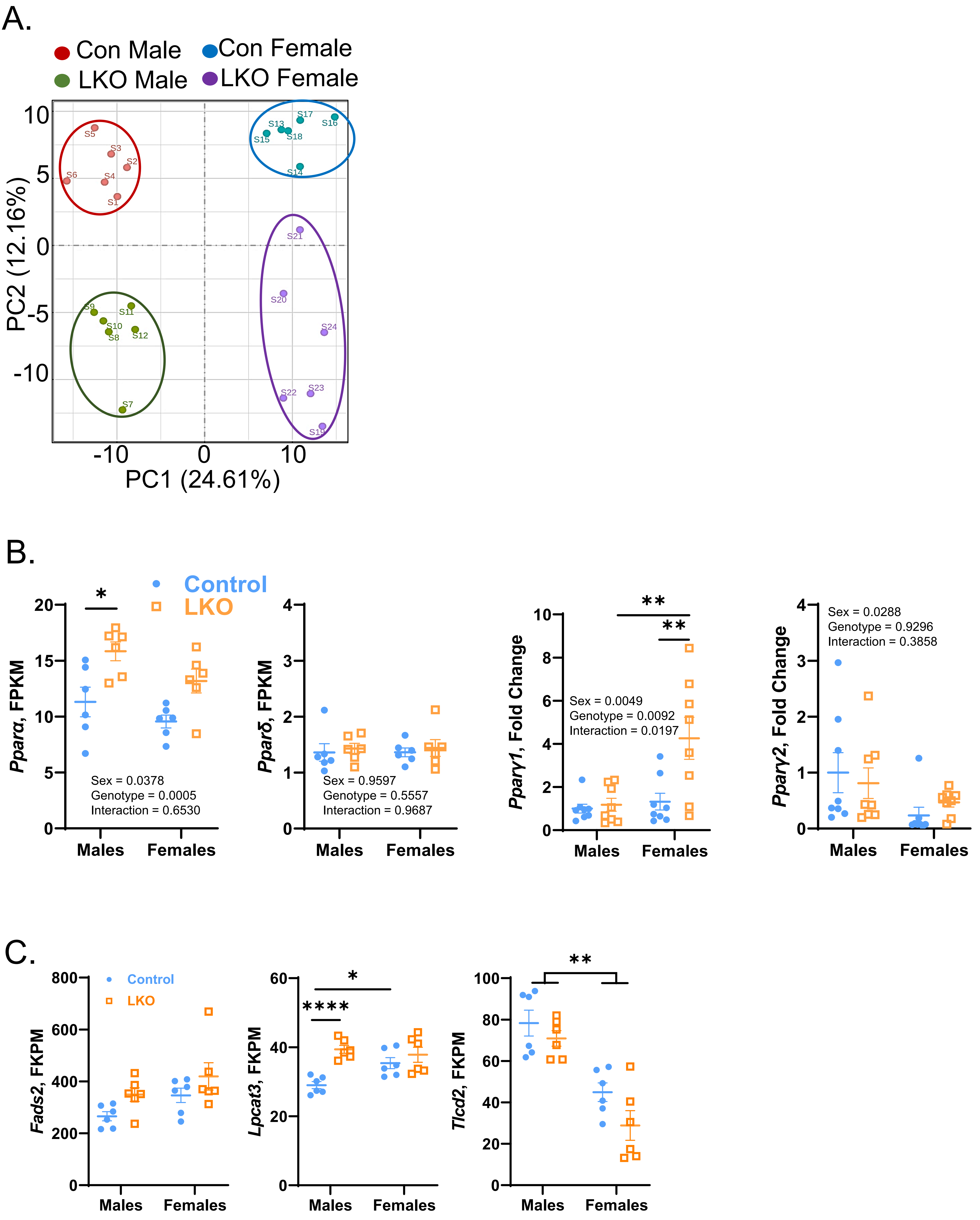

### Supplemental Figure 14

A.

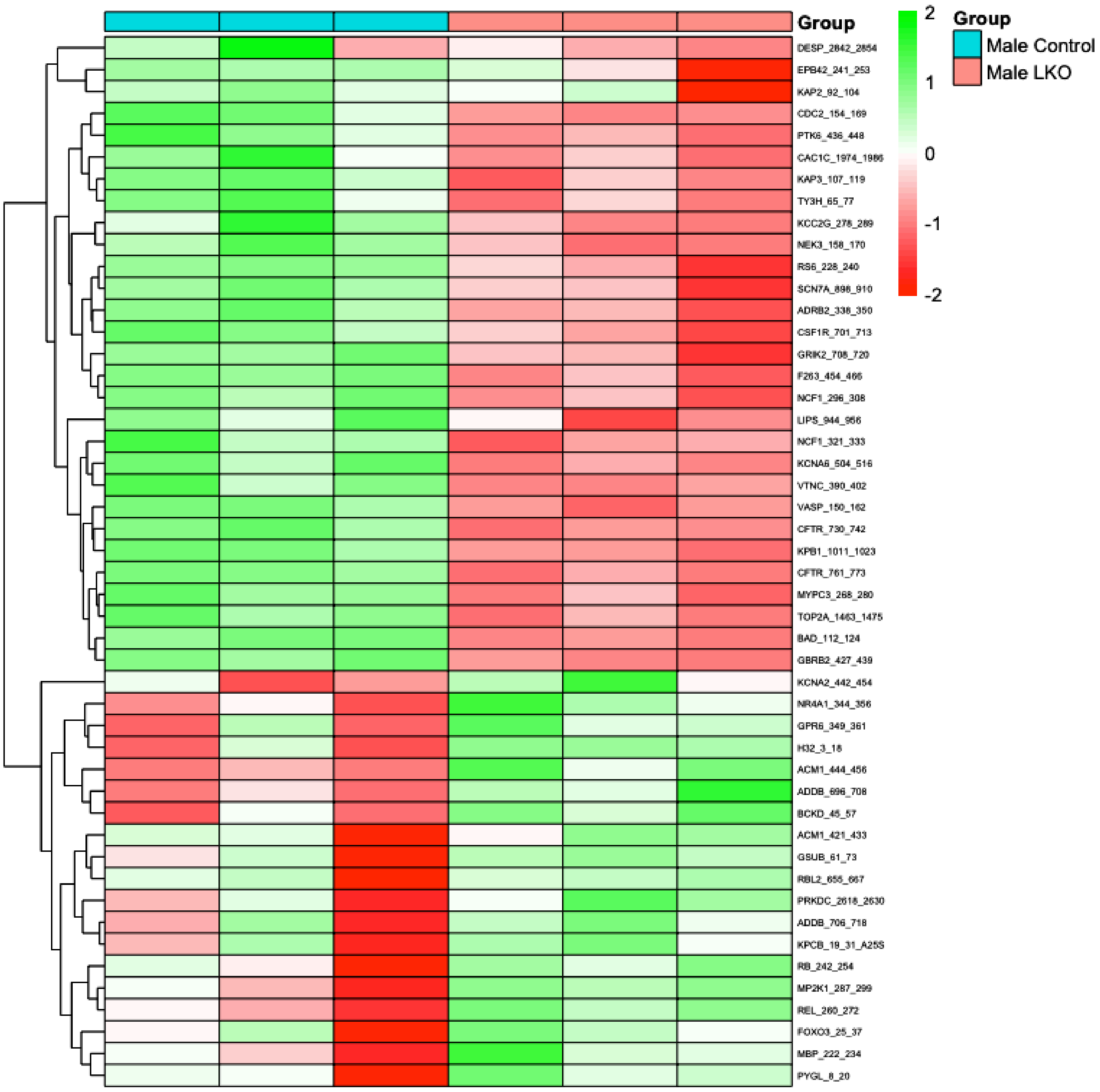


B.

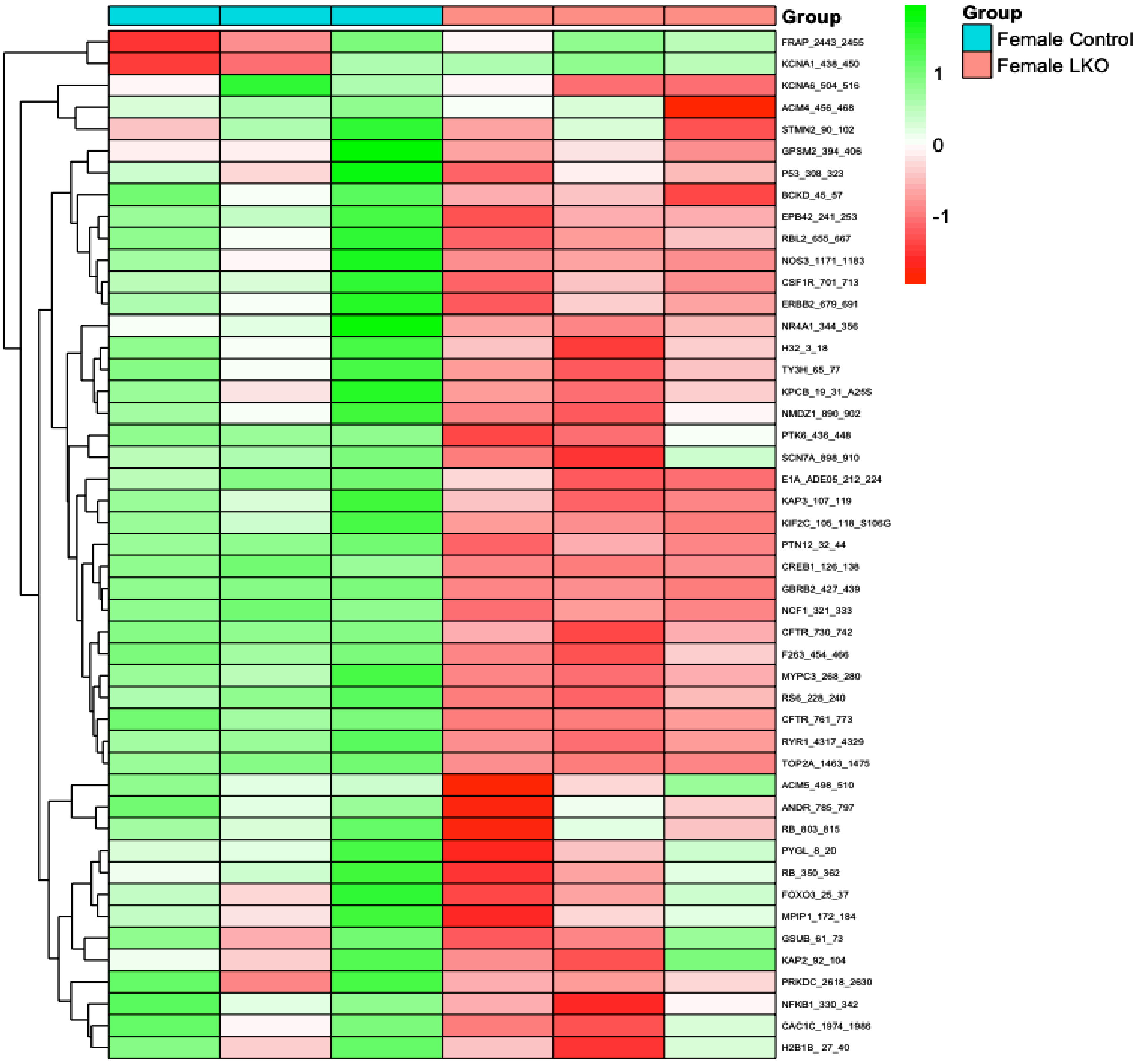

**Supplemental Figure 15**

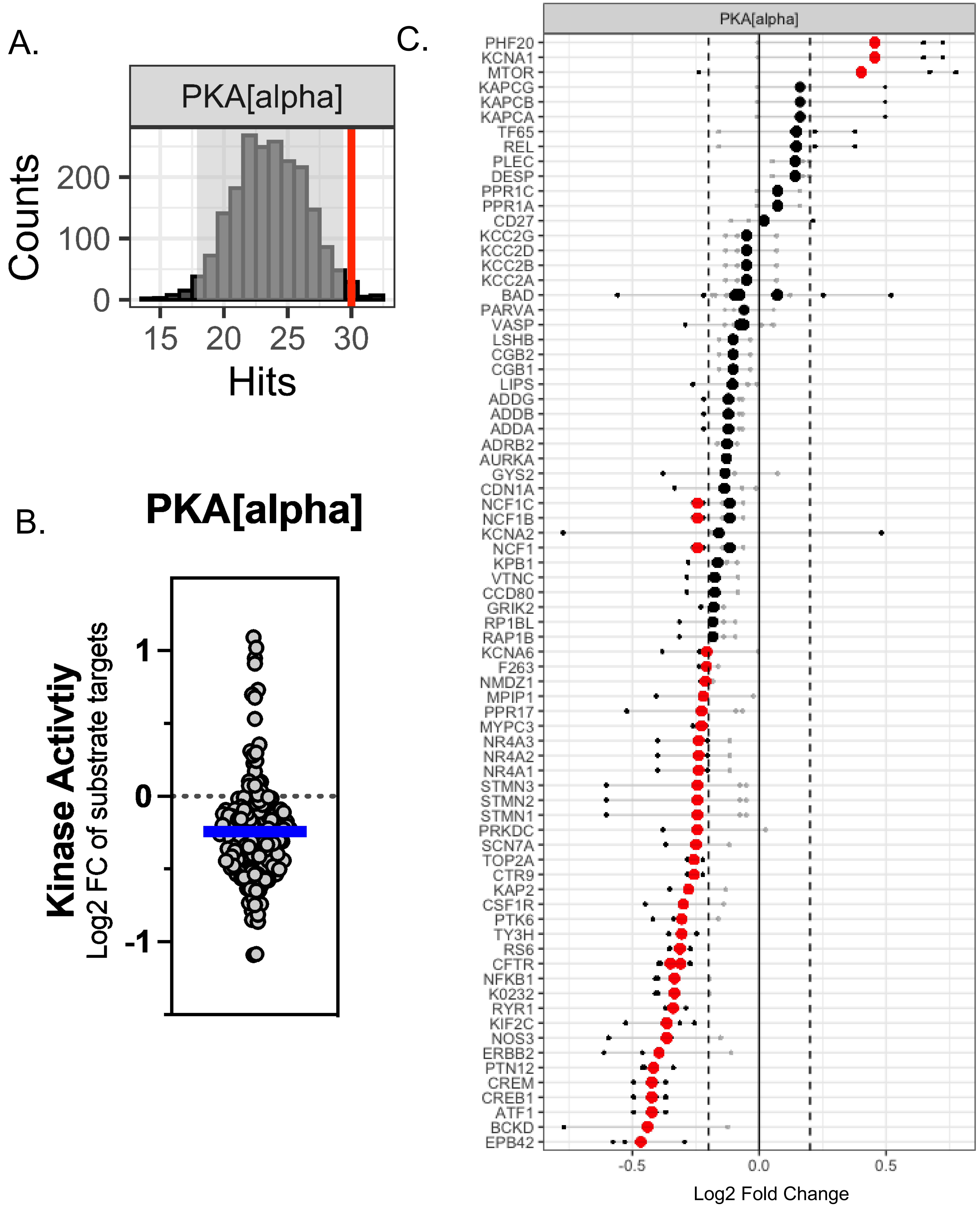
